# The double round-robin population unravels the genetic architecture of grain size in barley

**DOI:** 10.1101/2022.04.26.489546

**Authors:** Asis Shrestha, Francesco Cosenza, Delphine van Inghelandt, Po-Ya Wu, Jinquan Li, Federico A. Casale, Marius Weisweiler, Benjamin Stich

## Abstract

Grain number, size and weight primarily determine the yield of barley. Although the genes regulating grain number are well studied in barley, the genetic loci and the causal gene for sink capacity are poorly understood. Therefore, the primary objective of our work was to dissect the genetic architecture of grain size and weight in barley. We used a multi-parent population developed from a genetic cross between 23 diverse barley inbreds in a double round-robin design. Seed size-related parameters such as grain length, grain width, grain area and thousand-grain weight were evaluated in the HvDRR population comprising 45 recombinant inbred line sub-populations. We found significant genotypic variation for all seed size characters and observed 84 % or higher heritability across four environments. The results of the quantitative trait locus (QTL) detection indicate that the genetic architecture of grain size is more complex than reported previously. In addition, both cultivars and landraces contributed positive alleles at grain size QTLs. Candidate genes identified using genome-wide variant calling data for all parental inbred lines indicated overlapping and potential novel regulators of grain size in cereals. Furthermore, our results indicated that sink capacity was the primary determinant of grain weight in barley.

**Highlight:** Multi parent population uncovered the natural allelic series across quantitative loci associated with grain size and weight that will contribute to identifying causal genes and yield improvement in barley.

## Introduction

Barley (*Hordeum vulgare* L.) is one of the economically most important cereal crops after rice, wheat and maize (Newton *et al.*, 2011). Barley grain is primarily used in the feed and brewing industry (Dawson *et al.*, 2015). Large and plump grains contain a higher proportion of extractable sugar compared to small grains and, thus, are ideal for brewing (Walker *et al.*, 2013). Identifying the genetic regulators of natural grain size variation in barley is essential to creating additional phenotypic diversity by transgenesis (review in Scheben and Edwards, 2018). However, also classical breeding is facilitated by such information when used for marker-assisted selection.

Grain development is a complex process governed by multiple genes (Li *et al.*, 2018). The genetic basis of different grain size parameters is extensively studied in rice compared to other cereals. Several quantitative trait loci (QTLs) and the underlying genes controlling grain length (GL), grain width (GW) and weight have been isolated using map-based cloning in rice (Li *et al.*, 2019). Most of the genes control cell proliferation and cell expansion of the hulls (Li *et al.*, 2018). The described genes in rice are primarily transcriptional regulators or signaling proteins involved in G-protein, MAP kinase and hormonal signaling pathways (Li *et al.*, 2018).

Besides, a considerable effort has also been made to dissect the genetic architecture of kernel size in maize. Association and linkage mapping studies have identified genetic loci linked to grain size and weight in maize (Wang *et al.*, 2013; Liu *et al.*, 2014; Chen *et al.*, 2016; Liu *et al.*, 2017; Zhang *et al.*, 2017; Hao *et al.*, 2019; Li *et al.*, 2021). In addition, comparative genetics combined with genetic mapping were used to validate genes associated with kernel size in maize (Li *et al.*, 2010b,*a*; Liu *et al.*, 2020b). Comparative genomics and transgenic approaches were used to identify and validate the functional wheat orthologs of rice (Bednarek *et al.*, 2012; Ma *et al.*, 2016; Sajjad *et al.*, 2017) and Arabidopsis genes known to control seed size (Ma *et al.*, 2015a; Liu *et al.*, 2020a). Likewise, association mapping and linkage mapping studies, as well as functional validation of underlying genes, were performed in wheat (Kato *et al.*, 2000; Gegas *et al.*, 2010; Bennett *et al.*, 2012; Zhang *et al.*, 2012; Hu *et al.*, 2016; Simmonds *et al.*, 2016; Geng *et al.*, 2017; Li and Yang, 2017).

In contrast to other cereals, the genetic regulation of grain size is not well studied in barley. The genes controlling spikelet branching influence the average seed weight in barley (Zwirek *et al.*, 2019). Until now, five genes associated with spikelet branching, i.e. controlling 2 vs. 6 rowed spikes, were characterized in barley. The first characterized gene was a homeodomain-leucine zipper I-class type transcriptional regulator (*vrs1*) that suppressed the lateral spikelet fertility (Komatsuda *et al.*, 2007). Ramsay *et al.* (2011) showed that the allelic variation of the barley ortholog of maize teosinte branched 1 (*vrs5/int-c*) modifies *Vrs1* regulated lateral spikelet fertility. The mutant analysis also identified other row type genes such as *RAMOSA2/Vrs4* (Koppolu *et al.*, 2013), *SHI/Vrs2* (Koppolu *et al.*, 2013) and jumonji C-type H3k9me2/me3 demethylase/*Vrs3* (Bull *et al.*, 2017; van Esse *et al.*, 2017). Natural allelic variations were only reported for *Vrs1* and *Vrs5/Int-c* (Casas *et al.*, 2018).

Only a few mapping studies were performed to identify the genetic factors controlling grain size-related characters in barley. Walker *et al.* (2013) detected QTLs on all seven barley chromosomes in a double-haploid population for grain size characters such as GL, GW, test weight and grain volume. Pauli *et al.* (2014) identified five QTLs in an association mapping panel of the US spring barley collections for test weight of barley. Similarly, Watt *et al.* (2019) detected ten stable QTLs for GL, grain thickness, GW and plumpness. The same group performed positional cloning of a QTL associated with GL on Chr 2 and reduced the interval to three high confidence genes (Watt *et al.*, 2020). Pasam *et al.* (2012) identified 21 QTLs distributed across all seven chromosomes linked to thousand-grain weight (TGW) in an association mapping panel of spring barley. Zhou *et al.* (2016) detected two major QTLs for GL on Chr 3 and 4 in a recombinant inbred line population. Similarly, Xu *et al.* (2018) detected several common QTLs for grain size characters: GL, TGW, GW, GW/GL ratio, and grain area (GA) in a European barley cultivar association panel. Wang *et al.* (2019) identified 27 QTL hotspots (common QTLs shared between the characters) for eight-grain size parameters in a double-haploid population. Recently, in a double-haploid population, Wang et al. (2021) also detected four and three QTLs for GL and GW.

The studies mentioned above identifying genetic factors for seed size in barley were either performed in a single bi-parental population or an association panel. However, the main pitfall of bi-parental mapping is that it only utilizes a small fraction of species-wide allelic diversity and the limited number of recombination events within the population leads to a low resolution of mapping (Scott *et al.*, 2020). An alternative approach is to utilize genome-wide association studies (GWA) mapping, which improves the localization of QTLs. Nevertheless, the GWAS approach has limitations, including false marker-trait associations occurring due to confounding effects of population structure and kinship (Stich *et al.*, 2006, 2008). Also, GWA studies lack the power to detect the minor alleles in the population that might be associated with the trait of interest. Therefore, multi-parent populations (MPP) are utilized by plant geneticists and take advantage of linkage and association mapping (Stich, 2009). Nested association mapping (NAM) is a widely used MPP and developed by crossing a diverse set of inbreds to a common founder line producing multiple recombinant inbred line (RIL) families of bi-parental crosses (Yu *et al.*, 2008; Buckler *et al.*, 2009; Garin *et al.*, 2021). Stich (2009) and Klasen *et al.* (2012) investigated alternative crossing designs to create MPP populations concerning the power of QTL detection. The former computer simulation study found that a double round-robin crossing scheme to develop the MPP might be a resource-effective approach with respect to the number of required crosses without compromising the power of QTL detection compared to half diallel crossing schemes.

The study’s objective was

- to demonstrate the utility of the new MPP, the double round-robin population of barley (HvDRR) by exemplarily dissecting the genetic architecture of grain size in barley,
- to estimate the allelic series underlying each QTL,
- to identify candidate genes for the detected QTLs, and
- to identify promising QTL and sub-populations for fine mapping and QTL cloning.

## Materials and methods

### Plant materials and genotyping

We used 45 RIL populations developed from the genetic crosses among 23 spring barley inbreds (Casale *et al.*, 2021). The inbred lines used to develop the HvDRR population were selected from the barley diversity panel described by (Haseneyer *et al.*, 2010) to maximize a genotypic and trait diversity index (Weisweiler *et al.*, 2019). The 23 parental inbreds were crossed in a double round-robin design, where each inbred was crossed with four other inbreds (Fig.1 and Fig. S1). From each cross, a RIL population was developed following the single seed descent principle.

**Fig. 1:**
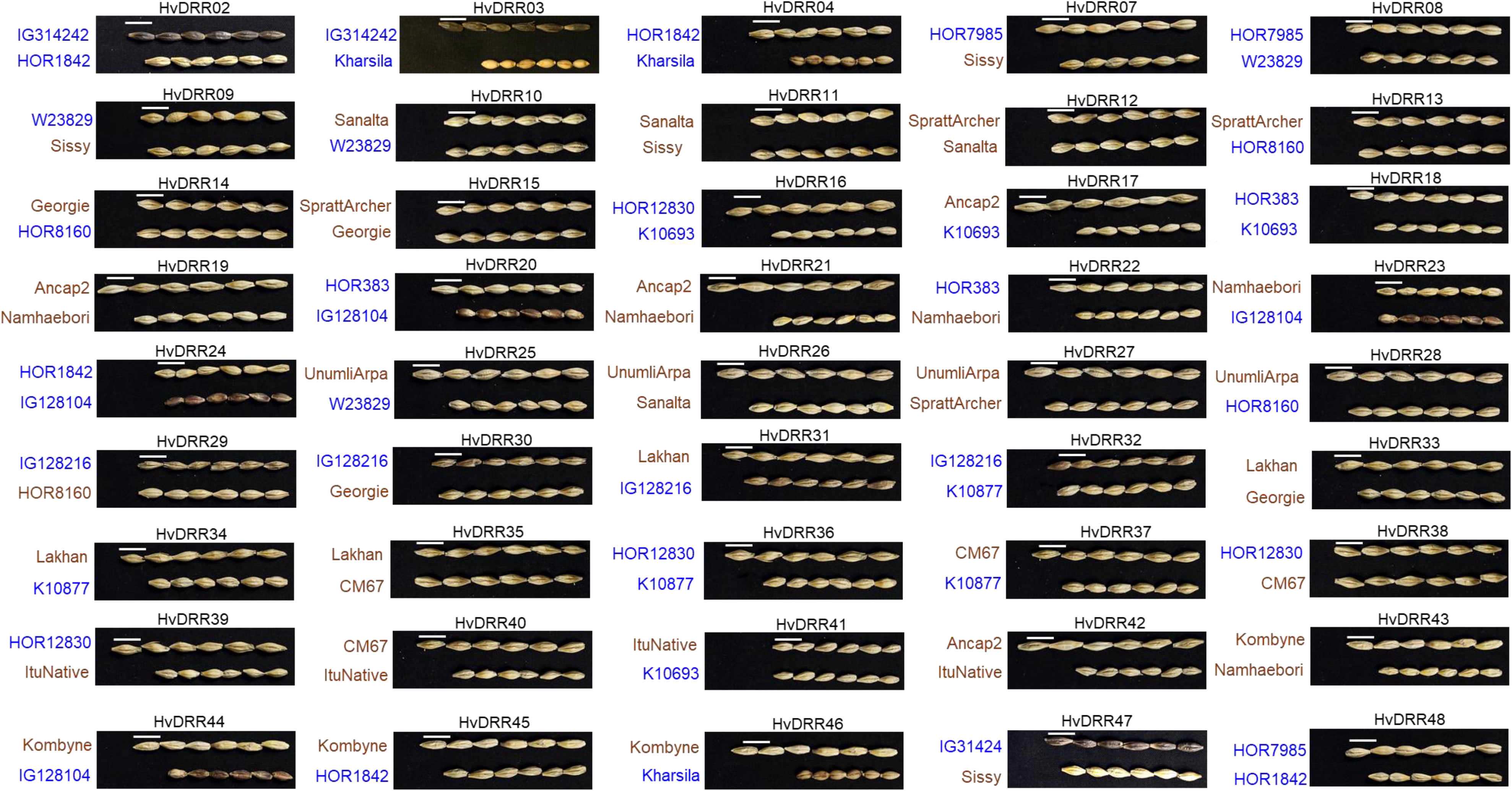
Image of grains of parental inbreds of 45 HvDRR sub-populations. The white scale bar indicates a length of 10 mm. Brown and blue colors designate germplasm type cultivars and landraces, respectively.

### Field experiment and data collection

Each RIL was grown in field experiments from 2017 to 2019. Data were collected from the field trials in Quedlinburg in 2018 and Cologne from 2017 to 2019, amounting to four different environments. We followed an augmented incomplete block design with one replicate for each RIL in each of the four environments. The RILs from a single HvDRR sub-population were sown in neighboring rows with 30 seeds per genotype in a 160 cm row. The parental inbreds were replicated as repeated checks across 16 to 20 rows randomized across the entire trial at each environment. Border plots were also maintained surrounding the entire trial. The ears were harvested and dried for a minimum of three days before threshing. MARViN seed analyzer (MARViNTECH GmbH, Germany) was used to estimate the seed size parameters for a random subset of around 60 seeds from each row of the field experiment. The camera mounted on the device recorded the image of the seed spread on the imaging platform. The seed image is then analyzed by the in-built software and returns seed size characters, including the count of seeds on the platform, GL (mm), GW (mm) and GA (mm^2^). Furthermore, TGW was measured for all rows.

### Genotyping

In F4 generation, 35 to 146 RILs from each of the 45 HvDRR sub-populations (Fig. S1) were genotyped using a 50K barley SNP array (Bayer *et al.*, 2017). The genetic map for individual HvDRR sub-populations and consensus map across the HvDRR population were established as described by Casale *et al.* (2021) and used for genomic prediction and linkage mapping.

### Statistical analyses

Dara were processed and statistical analyses was performed using R platform (R core team). A correction of field heterogeneity was performed at each environment based on the field map of the augmented design using the R package mvngGrAd package (Technow 2016). Adjusted entry means for original and field heterogeneity corrected values were estimated by using a mixed linear model with fixed genotype effects and random location and genotype:location effects:

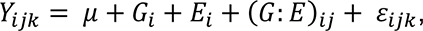

where *Y_ijk_* was the observed phenotypic value for *i*^*th*^ genotype in *j*^*th*^ environment for *k*^*th*^ replication, before or after field correction for the augmented design,

*μ* the general mean,

*G_i_* the effect of *i^th^* genotype,

*E_i_* the effect of *j^th^* environment,

(*G: *E**)*_ij_* the interaction of *i^th^* genotype with *j^th^* location and

*ε_ijk_* the random error term.

For broad sense heritability (*H*^2^) estimation on entry mean basis, the variance component of 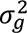 was calculated based on the above model, but with a random genotype effect and the following method was used:

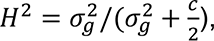

where *c* is the mean-variance of a difference of two adjusted entry means (Piepho and Möhring, 2007).

*H*^2^ across environments was highest for non-field heterogeneity adjusted data (Table S1). Therefore, the adjusted entry means across the four environments using the original data without field heterogeneity adjustment were used for subsequent analyses. Variance components were tested using a restricted likelihood ratio test using the R package RLRsim (Scheipl 2020).

In order to examine the mean differences between the segregating populations and the respective parental inbreds, the following mixed model was fitted to the data of all progenies and the two parental inbreds from all four environments:

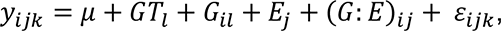

where *G*T*_i_* is the fixed effect of the germplasm type (either parental inbred or progeny).

### Genomic prediction

Genome-wide predictions of all phenotypes were performed according to Inghelandt *et al.* (2019) by genomic best linear unbiased prediction (GBLUP) using the following model (VanRaden, 2008):

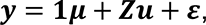

where *y* was the vector of the adjusted entry means of the corresponding phenotype, **1** the unit vector, *μ* the general mean, ***Z*** the design matrix that assigned the random effects to the genotypes and, ***u*** the vector of the genotypic effects that were assumed to be normally distributed with 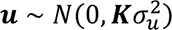 where ***K*** denotes the realized kinship matrix considering only additive effects calculated based on the SNP marker data mentioned above and 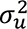 being the genotypic variance of the GBLUP model. In addition, ***ε*** was the vector of residuals following a normal distribution 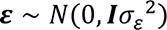, where ***I*** was the identity matrix and 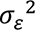 the residual variance.

Prediction ability was calculated as the Pearson correlation coefficient between observed and predicted phenotypes *r*(*y*, *ŷ*). Five-fold cross-validations (CVs) with 20 replications were performed to assess the model performance. The prediction ability was defined as the median of the predictions abilities across the 20 runs of 5 fold-CV.

### QTL analysis

In a first step, QTL were detected in the entire HvDRR population using the multi-parent population (MPP) procedure of the mppR package (Garin *et al.*, 2017). The MPP QTL analysis used a consensus map of the 45 HvDRR sub-populations. Next, the parental model was applied to estimate the multi-QTL effect with the assumption that each parent contributes a unique allele and the effect of a parent is constant across all crosses involving the parent in question. In short, the QTL detection procedure involved a simple interval mapping (SIM) in identifying the associated markers. Then, markers detected from the SIM scan were used as cofactors to run a first composite interval mapping. Finally, a second composite interval mapping scan was run by using a backward elimination procedure to obtain a final list of QTL for the grain size characters.

In the second step, we also performed a single population QTL analysis (SPA) for grain size and TGW in 45 individual HvDRR populations using the R package qtl (Broman and Sen, 2009). First, genotype probabilities of every 1cM were estimated, followed by a genome-wide scan for marker-trait association using Haley-Knott regression approximation. A forward search algorithm was then applied to perform multiple QTL mapping. A permutation test of 4000 runs was performed and the LOD score at 0.05 significance level was set as a threshold for QTL detection. A 1.5 LOD drop sets the confidence interval on either side of the detected QTL (Dupuis and Siegmund, 1999). The SPA QTLs detected across the HvDRR sub-populations were summarized as consensus QTLs. For the consensus QTLs, the shortest possible confidence interval of co-located QTLs (across sub-populations and phenotypes) was used to define the interval of consensus QTL. The confidence interval remained the same in the case of QTL detected in only one sub-population for a single seed size or weight character.

Finally, a two-dimensional genome-wide scan was performed to detect epistatic loci. The LOD score for interaction at 0.05 significance level obtained from the permutation test of 1000 runs was set as a significance threshold.

### Allelic series estimate

To understand the parental inbreds contribution to the genetic makeup of grain size and weight, we considered the effect sizes from the above described MPP QTL analysis. A multiple comparison test was performed on the allelic effect of each parental inbreds across all MPP QTLs. Then we calculated the cumulative allele effect for each inbred as the sum of the negative or positive standardized allele effects. The standardized allele effect for an inbred was the difference between the mean of the estimated allele effect for the 23 parental inbreds and the estimated allele effect of the corresponding inbred.

### QTL validation

We developed a high-resolution mapping population that segregates for a GL-associated QTL on chromosome 1H by crossing two RILs from the HvDRR33 sub-population, namely Hv-2018S-7-03050 and Hv-2018S-7-03225 harboring Lakhan and Georgie allele, respectively. Cleaved amplified polymorphic sequence markers were developed for genotyping the parent-specific SNP allele at the left border (JHI-Hv50k-2016-43007) and right border (JHI-Hv50k-2016-43714) of the QTL interval for 924 F2 progenies (Table S2). For genotyping, DNA was extracted from one-week-old seedlings and the SNP loci at left and right borders were PCR amplified. Then, the PCR product on the left and right border were digested using XbaI and BspEI, respectively. Digested products were separated on a 1.5 % agarose gel and visualized under UV-transilluminator. Recombinants were selected and transferred to 1.5 L pots. The grains were harvested, and GL was evaluated using MARViN seed analyzer. The GL was also evaluated from three non-recombinant F2 plants, each carrying either Lakhan or Georgie allele at both borders of the QTL interval.

### Candidate gene analysis

A detailed candidate gene analysis was performed for the consensus QTLs smaller than 10 Mb that explained more than 10 % of the phenotypic variance. First, the list of high confidence genes present in the consensus QTLs were retrieved from the plant genomics & phenomics research data repository, IPK Gatersleben, Germany. The annotation and coordinates were based on the Morex v2 gene models (Monat *et al.*, 2019). In the next step, we used the variant calling data obtained from the whole genome sequencing project of the 23 inbreds (Weisweiler *et al.*, 2022) to select the genes that showed polymorphisms between the parental inbreds. We examined non-synonymous (amino acid substitutions) or deleterious synonymous mutations, indels in the coding region, as well as indels and structural variants (SVs) in the 5-end regulatory region (5 kb upstream of the start codon). The genes with polymorphisms co-segregating between the parental groups with distinct allele effects on grain size and TGW at consensus QTLs were considered candidate genes. Finally, Sanger sequencing was used to validate the mutations detected in the variant calling data for two candidate genes.

## Results

### Phenotypic variation for grain size and weight

Grain size and weight phenotypes were collected across four different environments for around 4500 RILs from the 45 sub-populations. The parental inbreds were replicated between 16 to 20 times across the experiment in each environment as repeated checks to account for field heterogeneity. We observed significant (p < 0.0001) genotype, environment, and genotype*environment effects for all four characters GW, GL, GA and TGW (Table 1). On an entry mean basis, broad-sense heritability was 84 % or above for all characters (Table 1). Although a significant positive correlation was observed between all grain size characters, the degree of association between TGW and GW (r = 0.84) was about 50% higher than with GL (r = 0.4) (Fig. 2). In addition, we observed contrasting relations among the grain size characters for some sub-populations compared to the whole HvDRR population. For instance, GL showed no or weak correlation with GW and TGW for HvDRR18 and HvDRR32. In general, GA showed significant positive correlations with all characters at the whole HvDRR population as well as sub-populations level (Fig. S2 and Fig. 2).

**Fig. 2:**
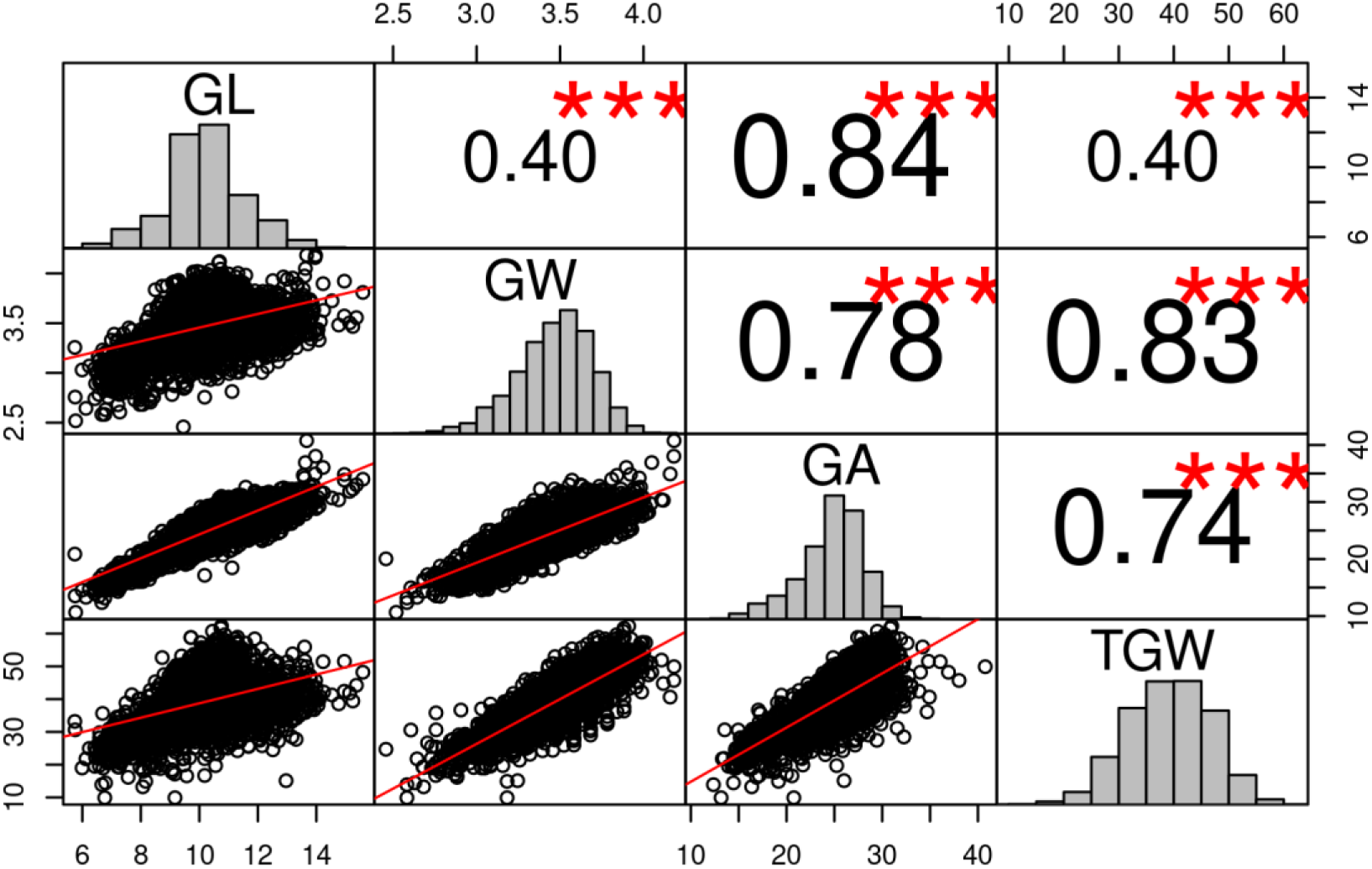
Pairwise correlation coefficients between four grain size characters grain area (GA), grain length (GL), grain width (GW), and thousand-grain weight (TGW). Adjusted entry means of the recombinant inbred lines from 45 individual HvDRR sub-populations and the parental inbreds were used for estimating the correlation. Asterisks indicate that the observed correlation coefficient estimates were statistically different from zero (***, p ≤ 0.001).

**Table 1:** Variance components and broad-sense heritability (*H*^2^) of grain size-related traits and thousand-grain weight (TGW). Asterisks indicate the significance of a likelihood ratio test (***, p ≤ 0.001).

| Character | Genotype | Environment | Genotype:Environment | Residual | $H^2$ |
| --- | --- | --- | --- | --- | --- |
| Grain length | 1.53*** | 0.57*** | 0.24*** | 0.56*** | 0.84 |
| Grain width | 0.04*** | 0.02*** | 0.01*** | 0.01*** | 0.85 |
| Grain area | 9.95*** | 0.16*** | 0.92*** | 2.43*** | 0.89 |
| TGW | 46.34*** | 20.92*** | 7.42*** | 17.72*** | 0.84 |

The adjusted entry means for GL and GW ranged from 5.7 to 15.6 mm and 2.5 to 4.2 mm, respectively, among the RILs of HvDRR sub-population (Fig. S3 and S4). Likewise, the lowest and highest observed adjusted entry means for GA were 10.7 mm^2^ and 40.2 mm^2^, respectively (Fig. S5). The adjusted entry means for TGW ranged between 10 and 62.3 g (Fig. S6). We also compared the average progeny performance with the parental average for 45 HvDRR sub-populations across all environments, as this indicates the importance of additive*additive epistatic variance. Mean progeny character values for GL, GW, GA and TGW were statistically different (p < 0.01) from the parental average in 16, 16, 18 and 17 HvDRR sub-populations for GL, GW, GA and TGW, respectively (Fig. S3-6). Among them, 7 sub-populations showed significantly (p < 0.01) different average parental and progeny means for all four characters. Overall, we observed a considerable genotypic variation for grain size and weight in the HvDRR population that can be utilized for breeding and genetic dissection of grain size and therewith yield-related characters in barley.

### QTL associated with grain size

First, we performed MPP analyses that detected 17, 24, 19 and 19 QTLs associated with GL, GW, GA and TGW, respectively. Nine of these QTLs were associated with all four evaluated characters (Fig. 3). The full model of additive QTLs in MPP explained 41.6 %, 46.9 %, 45.7 % and 40.4 % of the phenotypic variance for GL, GW, GA and TGW, respectively (Table 2). Likewise, the percentage of phenotypic variance (r² of prediction ability) explained by the GBLUP model based on genome-wide SNPs was 69.6 % (GW), 71.7 % (GL) and 73.4 % (GA and TGW), respectively (Table 3).

**Fig. 3:**
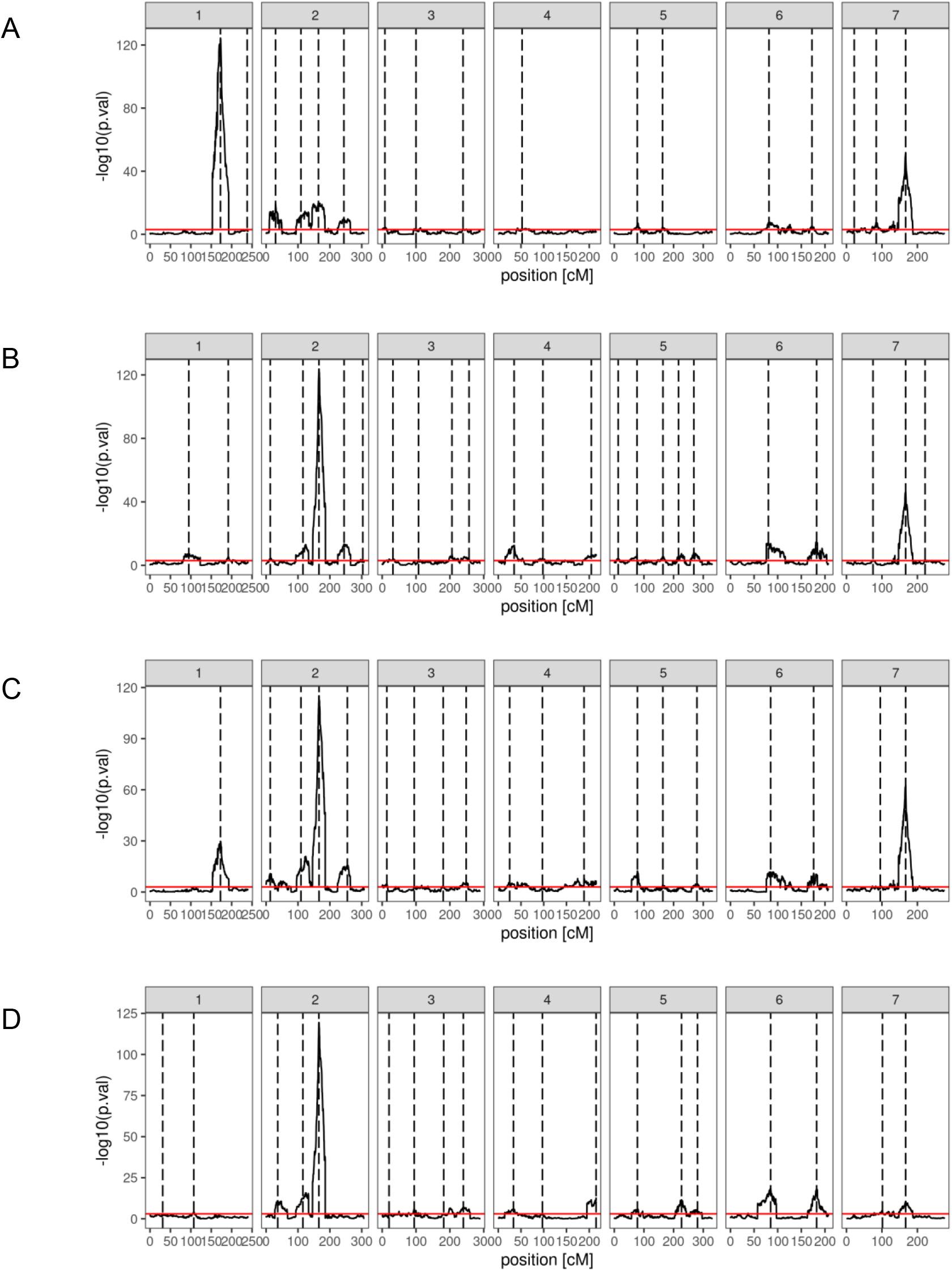
Quantitative trait loci (QTL) profile for grain size-related characters from multi-parent population analysis of 45 HvDRR sub-populations using the parental model. The QTLs were detected across all seven chromosomes using composite interval mapping [significant threshold of –log10(p) = 3]. The vertical dashed lines indicate the positions of the detected QTL. QTL profile for (A) grain length (B) grain width (C), grain area and (D) thousand-grain weight.

**Table 2:** Percent of explained phenotypic variance by the full model of additive quantitative trait loci (QTL) detected in multi-parent population analysis using a parental model.

| **Character** | **Explained phenotypic variance (%)** |
| --- | --- |
| Grain length | 41.6 |
| Grain width | 46.9 |
| Grain area | 45.7 |
| Thousand-grain weight | 40.4 |

**Table 3:** Prediction ability of the genome-wide SNP marker data for the grain size-related characters and thousand-grain weight (TGW). The predictions were performed using genomic best linear unbiased predictions. The first column indicates the prediction abilities without cross-validation (CV). The second and last columns are the median of prediction ability and standard deviation across 20 runs of 5 folds CV made with random sampling, respectively.

| **Character** | **Prediction ability overall 45 HvDRR** |  | **Standard deviation** |
| --- | --- | --- | --- |
|  | **Without CV** | **5 Fold CV 20 runs** | **5 Fold CV 20 runs** |
| Grain length | 0.919 | 0.847 | 0.012 |
| Grain width | 0.919 | 0.834 | 0.012 |
| Grain area | 0.932 | 0.856 | 0.012 |
| TGW | 0.92 | 0.857 | 0.009 |

Next, in SPA we detected QTLs in 38, 35, 39 and 33 of the HvDRR sub-populations for GL, GW, GA and TGW, respectively (Table S3-6). A total of 316 QTLs for four characters were detected in SPA of which 260 were observed at the same position as MPP QTLs (Fig. 4-6). When examining the detailed distribution of SPA QTLs, 99 and 56 were located on chromosomes 2H and 7H, respectively and the number of QTLs ranged between 29 and 36 for the other chromosomes (Table S3-6). The percent of explained variance (PEV) by individual QTLs in SPA ranged between 3 % and 70 % (Table S3-6). More than two-thirds of the QTLs detected in our study explained 10 % or more of the phenotypic variance for the associated character (Fig. S7). Finally, we summarized the SPA QTLs into 62 consensus QTLs, the shortest possible interval of co-located QTLs in SPA (Table 4). Only six consensus QTLs were present in the pericentromeric region.

**Fig. 4:**
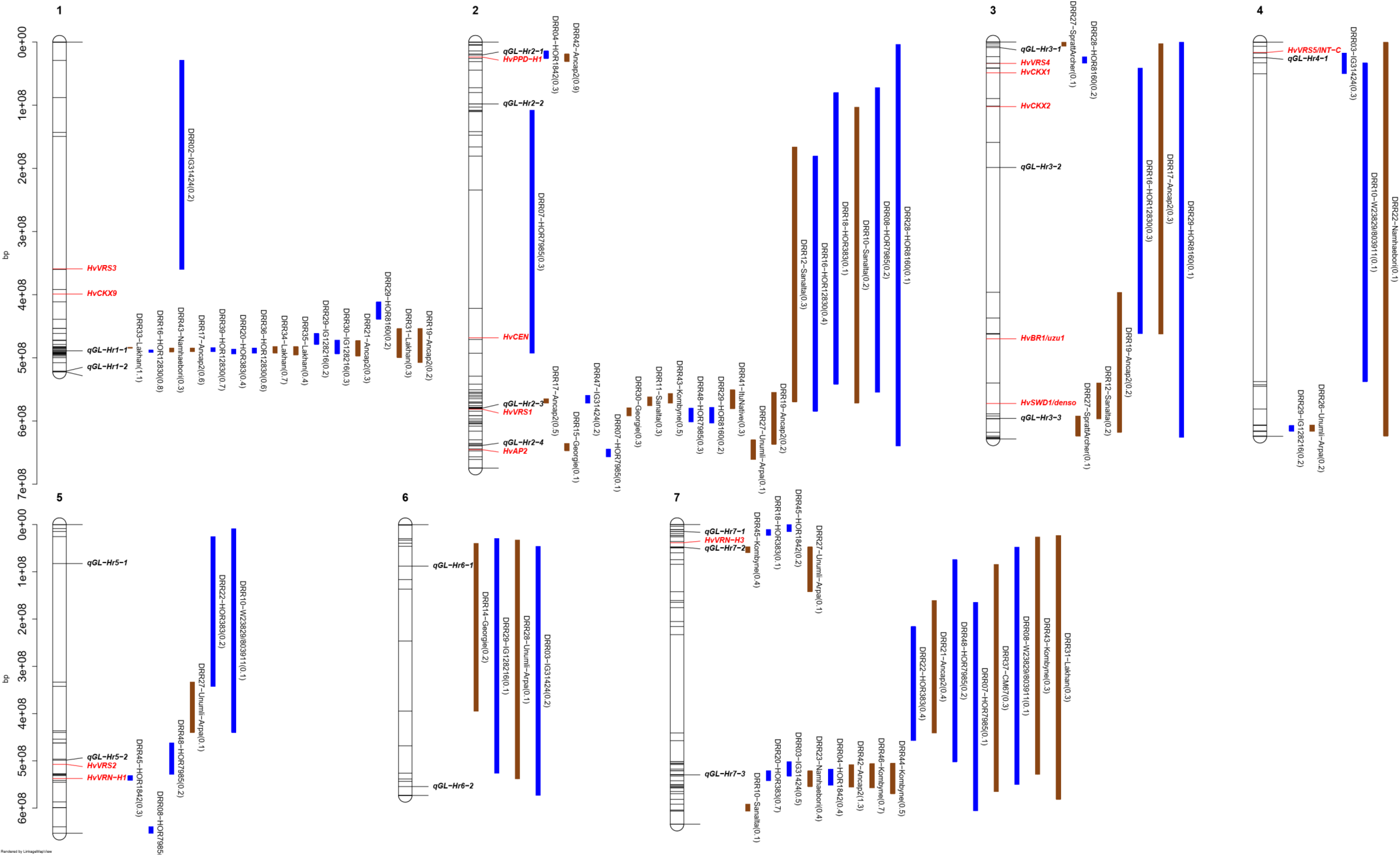
Confidence intervals of quantitative trait loci (QTLs) detected for grain length (GL) in single population analyses. QTL mapping was performed in 45 HvDRR sub-populations. The parental allele with a positive additive effect inside the parenthesis is connoted in the QTL name. The color of the confidence interval implies if the parent contributing a positive additive effect is a landrace (blue) or a cultivar (brown). The position of known genes controlling developmental phenotypes such as flowering time, plant height and spikelet branching are indicated in red letters. The loci in bold italics letters are the GL-associated QTLs detected in multi-parent population analysis using the parental model. A consensus map was used to order the marker for plotting the confidence intervals. The genetic positions of the top and bottom five markers of the genetic map and the markers flanking the QTL interval are indicated in the plot.

**Fig. 5:**
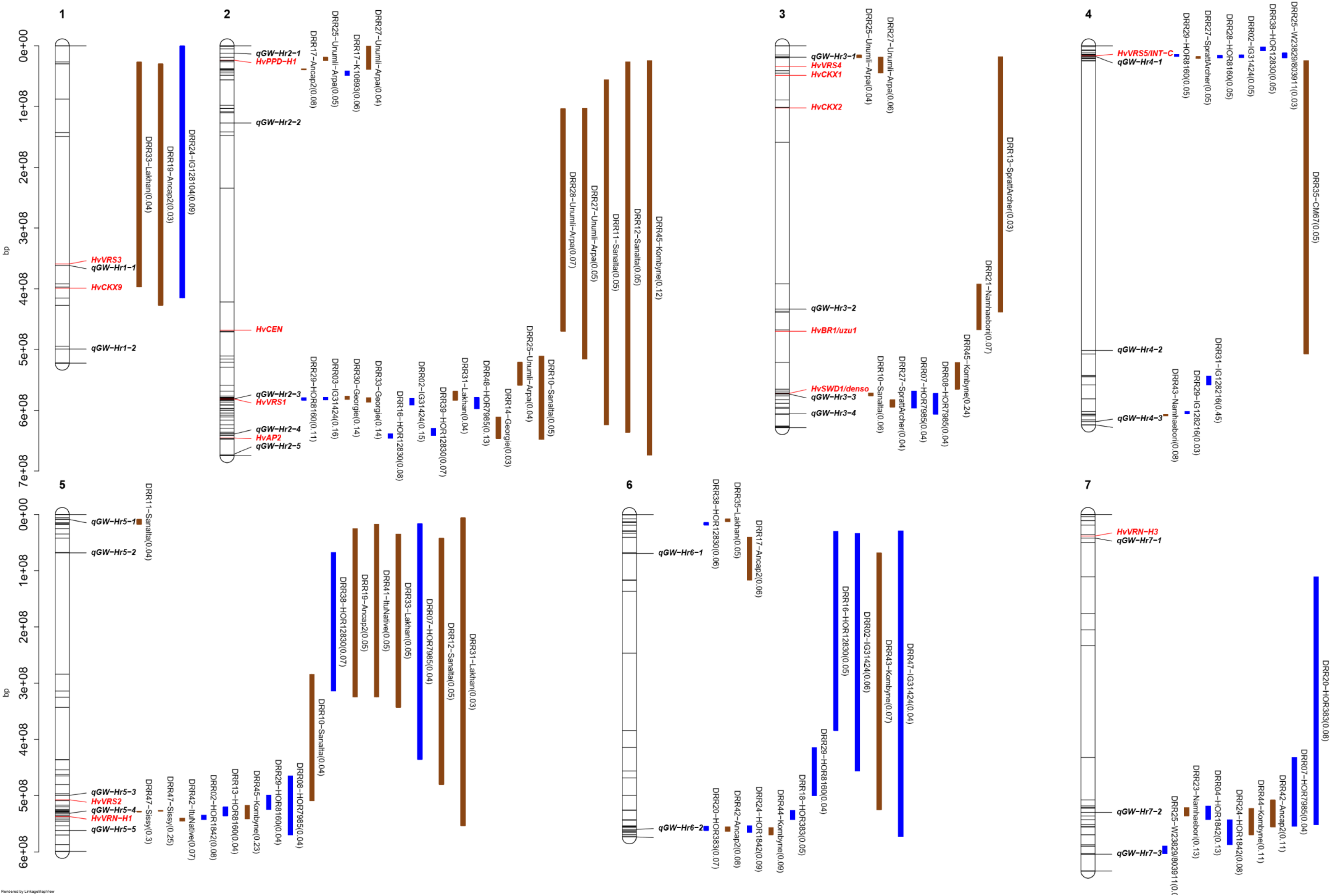
Confidence intervals of quantitative trait loci (QTLs) detected for grain width (GW) in single population analyses. QTL mapping was performed in 45 HvDRR sub-populations. The parental allele with a positive additive effect inside the parenthesis is connoted in the QTL name. The color of the confidence interval implies if the parent contributing a positive additive effect is a landrace (blue) or a cultivar (brown). The position of known genes controlling developmental phenotypes such as flowering time, plant height and spikelet branching are indicated in red letters. The loci in bold italics letters are the GW-associated QTLs detected in multi-parent population analysis using the parental model. A consensus map was used to order the marker for plotting the confidence intervals. The genetic positions of the top and bottom five markers of the genetic map and the markers flanking the QTL interval are indicated in the plot.

**Fig. 6:**
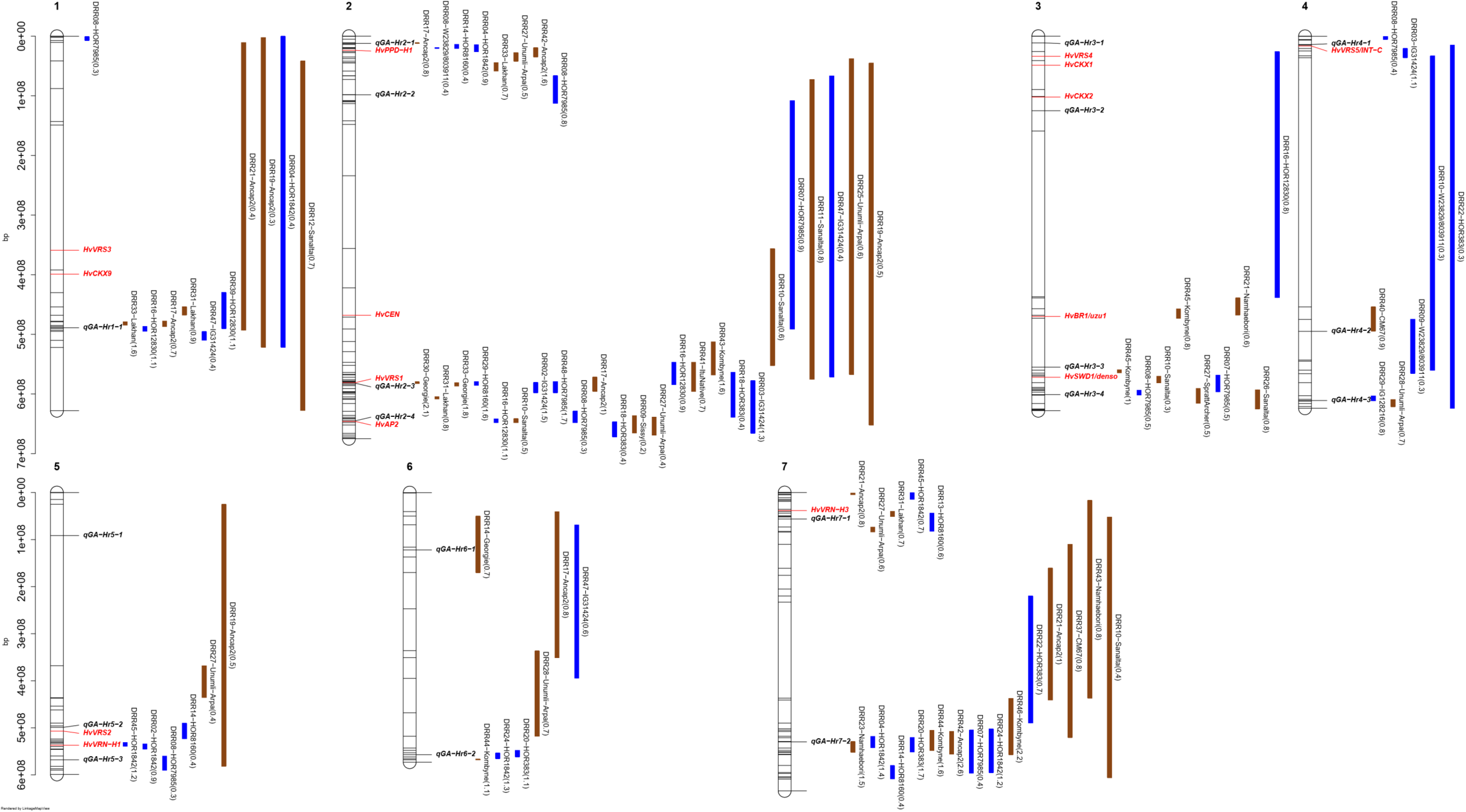
Confidence intervals of quantitative trait loci (QTLs) detected for grain area (GA) in single population analyses. QTL mapping was performed in 45 HvDRR sub-populations. The parental allele with a positive additive effect inside the parenthesis is connoted in the QTL name. The color of the confidence interval implies if the parent contributing a positive additive effect is a landrace (blue) or a cultivar (brown). The position of known genes controlling developmental phenotypes such as flowering time, plant height and spikelet branching are indicated in red letters. The loci in bold italics letters are the GA-associated QTLs detected in multi-parent population analysis using the parental model. A consensus map was used to order the marker for plotting the confidence intervals. The genetic positions of the top and bottom five markers of the genetic map and the markers flanking the QTL interval are indicated in the plot.

**Fig. 7:**
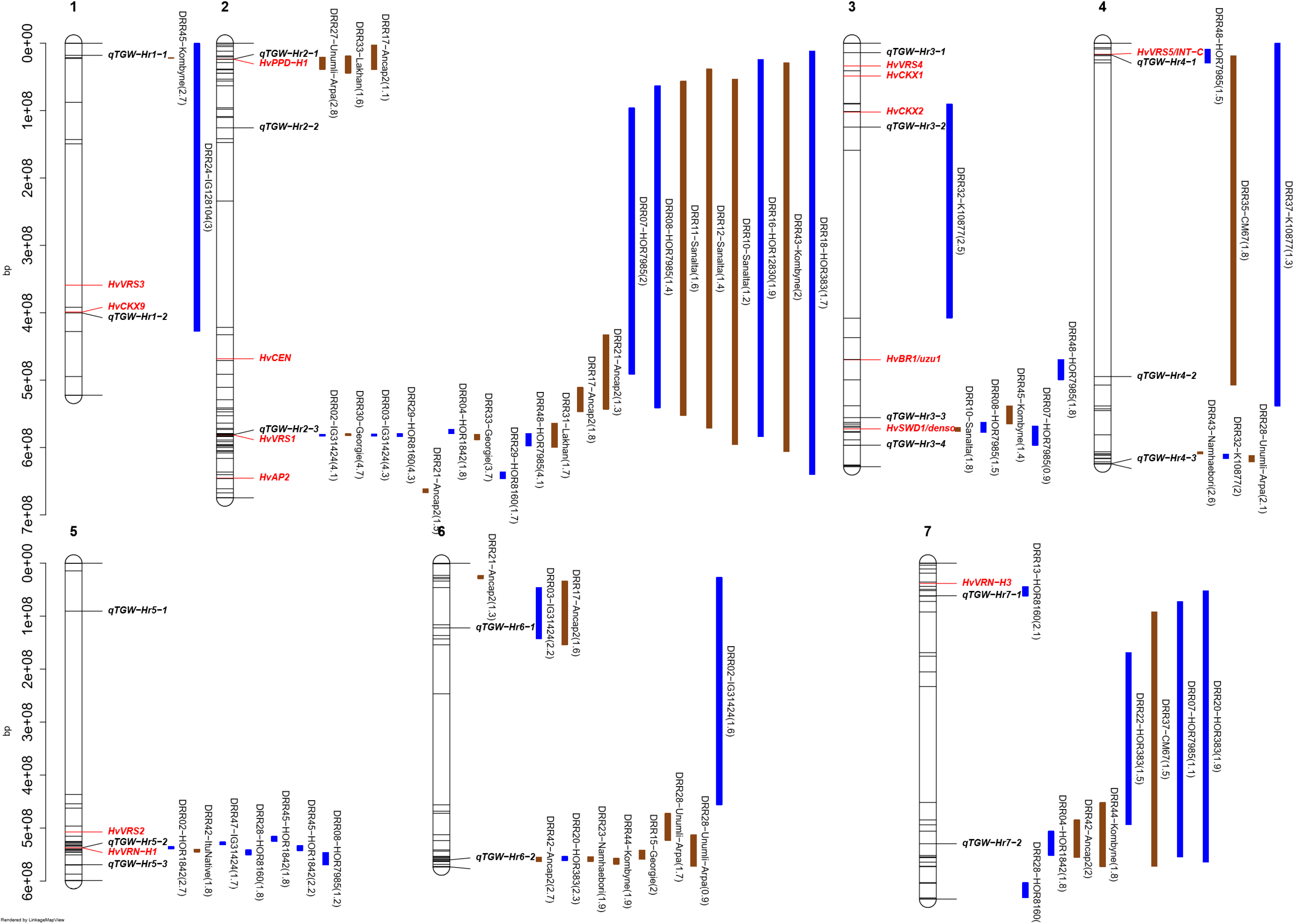
Confidence intervals of quantitative trait loci (QTLs) detected for thousand-grain weight (TGW) in single population analyses. QTL mapping was performed in 45 HvDRR sub-populations. The parental allele with a positive additive effect inside the parenthesis is connoted in the QTL name. The color of the confidence interval implies if the parent contributing a positive additive effect is a landrace (blue) or a cultivar (brown). The position of known genes controlling developmental phenotypes such as flowering time, plant height and spikelet branching are indicated in red letters. The loci in bold italics letters are the TGW associated QTLs detected in multi-parent population analysis using the parental model. A consensus map was used to order the marker for plotting the confidence intervals. The genetic positions of the top and bottom five markers of the genetic map and the markers flanking the QTL interval are indicated in the plot.

**Table 4:**
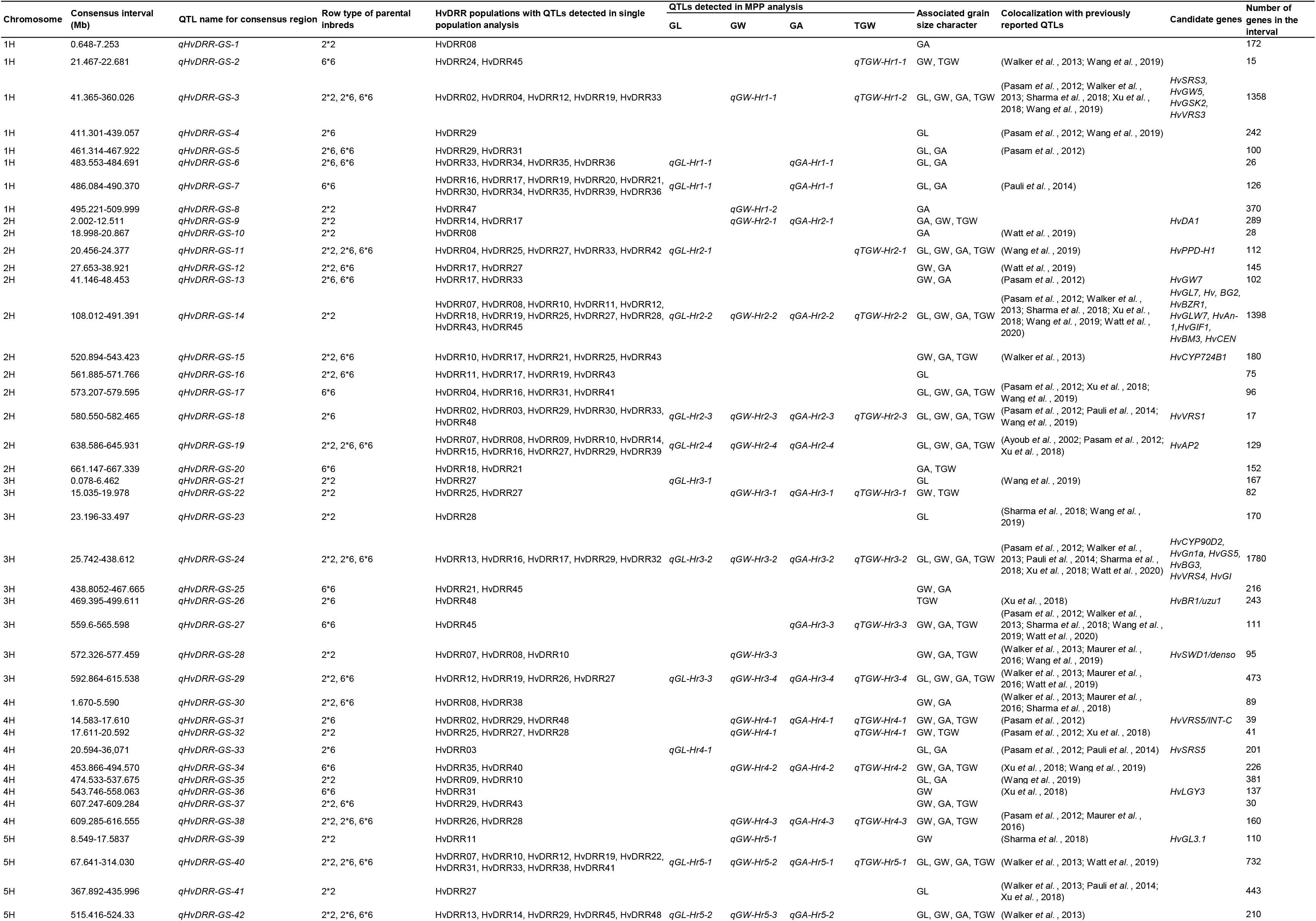
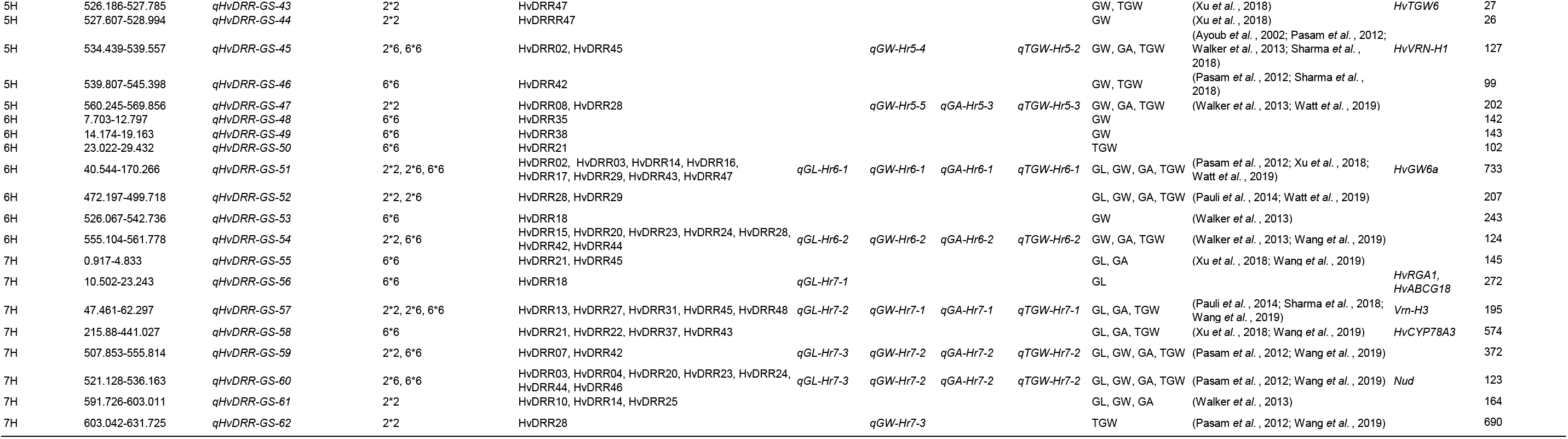
Consensus quantitative trait loci (QTLs) associated with grain length (GL), grain width (GW), grain area (GA) and thousand-grain weight (TGW) in HvDRR population. The consensus interval of QTL/ cluster of QTLs detected in single population analysis were summarized as consensus QTLs and the corresponding loci detected in multi-parent population analysis are indicated. The consensus QTLs that coincided with grain size and weight associated loci detected in previous genetic mapping studies are reported.

As explained above, we observed a significant difference between progeny and parental mean for a few sub-populations and, thus, evaluated the presence of epistatic QTL (Table S7). We detected 14 significant (p < 0.05) epistatic interaction QTLs. The epistatic interaction QTLs were frequently detected (10 times) in sub-populations with significant differences (p < 0.01) for the grain size and weight characters between parent and population mean compared to sub-populations without that feature (Table S7).

### QTL effect estimate

To understand the parental inbreds’ contribution to the genetic makeup of sink size and grain weight, we estimated the effect size of the parental inbreds across all QTLs detected in MPP QTL analysis using a parental model. That means one MPP QTL for a character will have 23 alleles corresponding to 23 parental inbreds, resulting in total QTL counts of 391, 552, 437 and 437 for GL, GW, GA and TGW, respectively (Fig. 8). The distribution of the QTL allele effect indicated that the differences in grain size and weight of 23 parental inbreds were caused by the cumulative effect of minor-effect alleles and a few large-effect alleles (Fig. 8). Only 10 % of the QTL alleles caused TGW to increase or decrease by 1 g or more (Fig. 8D). Similarly, 4 % of the QTLs affected GL, GW and GA by more than 2.5 mm, 0.5 mm and 1 mm^2^, respectively (Fig. 8A-C).

**Fig. 8:**
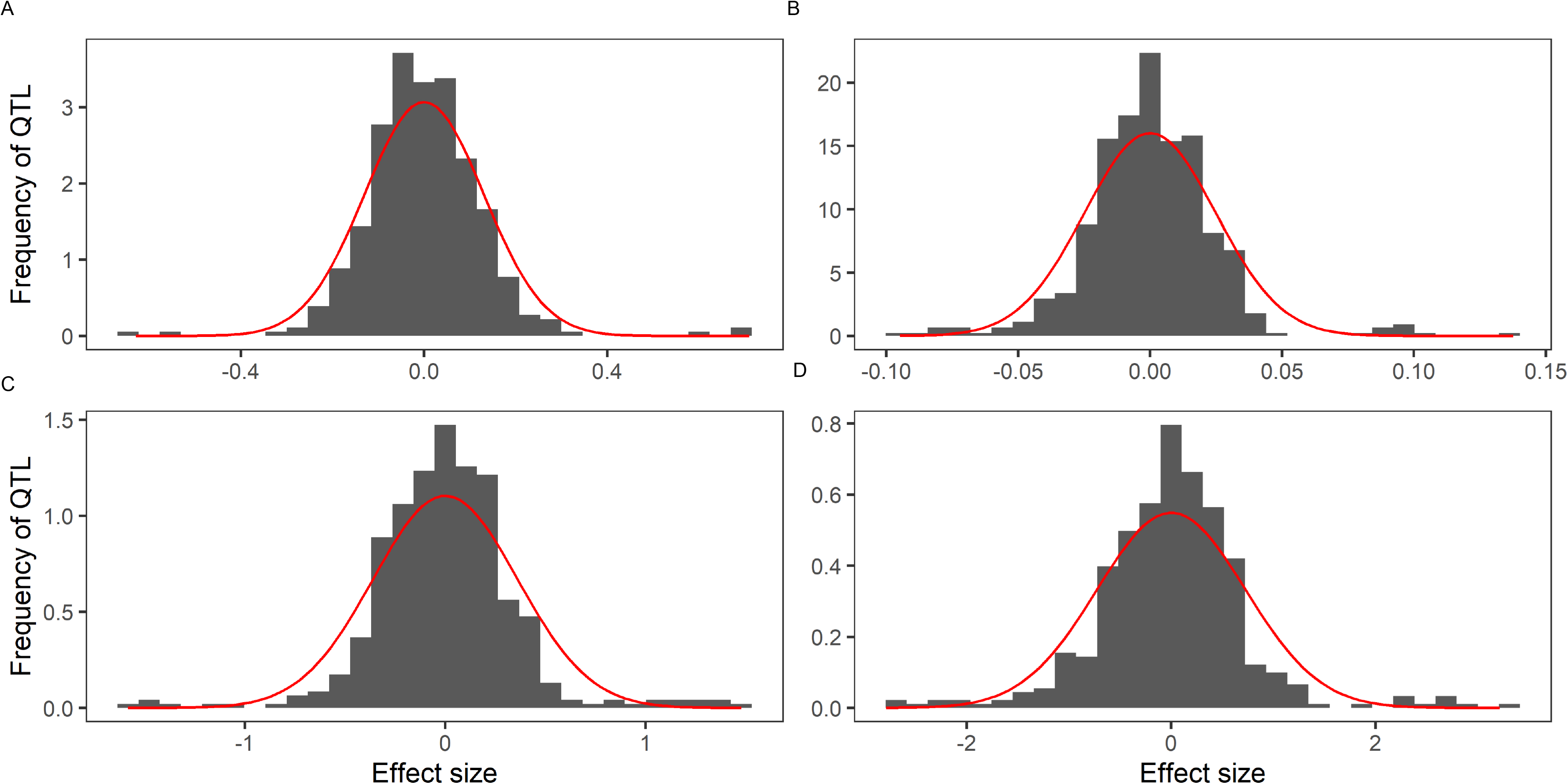
Histogram of standardized allele effect estimates for quantitative trait loci (QTLs) detected in multi-parent population analysis of 45 HvDRR sub-populations using a parental model for (A) grain length, (B) grain width, (C) grain area and (D) thousand-grain weight. The standardized allele effect for an inbred was the difference between the mean of the estimated allele effects for the 23 parental inbreds and the estimated allele effect of the considered inbred. A normal distribution curve (red color) is overlaid onto the histograms.

Next, allelic series across the MPP QTLs revealed that parental inbred contribute to both positive and negative allele effects (increase or decrease in grain size) (Fig. S8-S11). In addition, the allele effect of parental inbreds was statistically different within a single QTL locus (Fig. S8-11). Because we observed the allelic series for parental inbreds, we estimated the cumulative positive and negative allele effect across the MPP QTLs and counted the number of QTL for each inbred parental carries a positive or negative allele (Fig. 9). The net cumulative allele effect (cumulative decrease subtracted from cumulative increase) and the number of favorable alleles were lowest for a landrace IG128104 across the MPP QTLs for all four characters, followed by another landrace Kharsila. On the other hand, the HOR12830 (1.76 mm and 2.73 mm^2^) and Ancap2 (1.68 mm and 2.88 mm^2^) allele contributed to a higher net cumulative effect across GL and GA QTLs than the other parental inbreds. Sanalta, Unumli-Arpa and K10877 were the most significant contributor to the number of favorable alleles and the cumulative positive allele effect across GW QTLs detected in MPP analysis (Fig. 9).

**Fig. 9:**
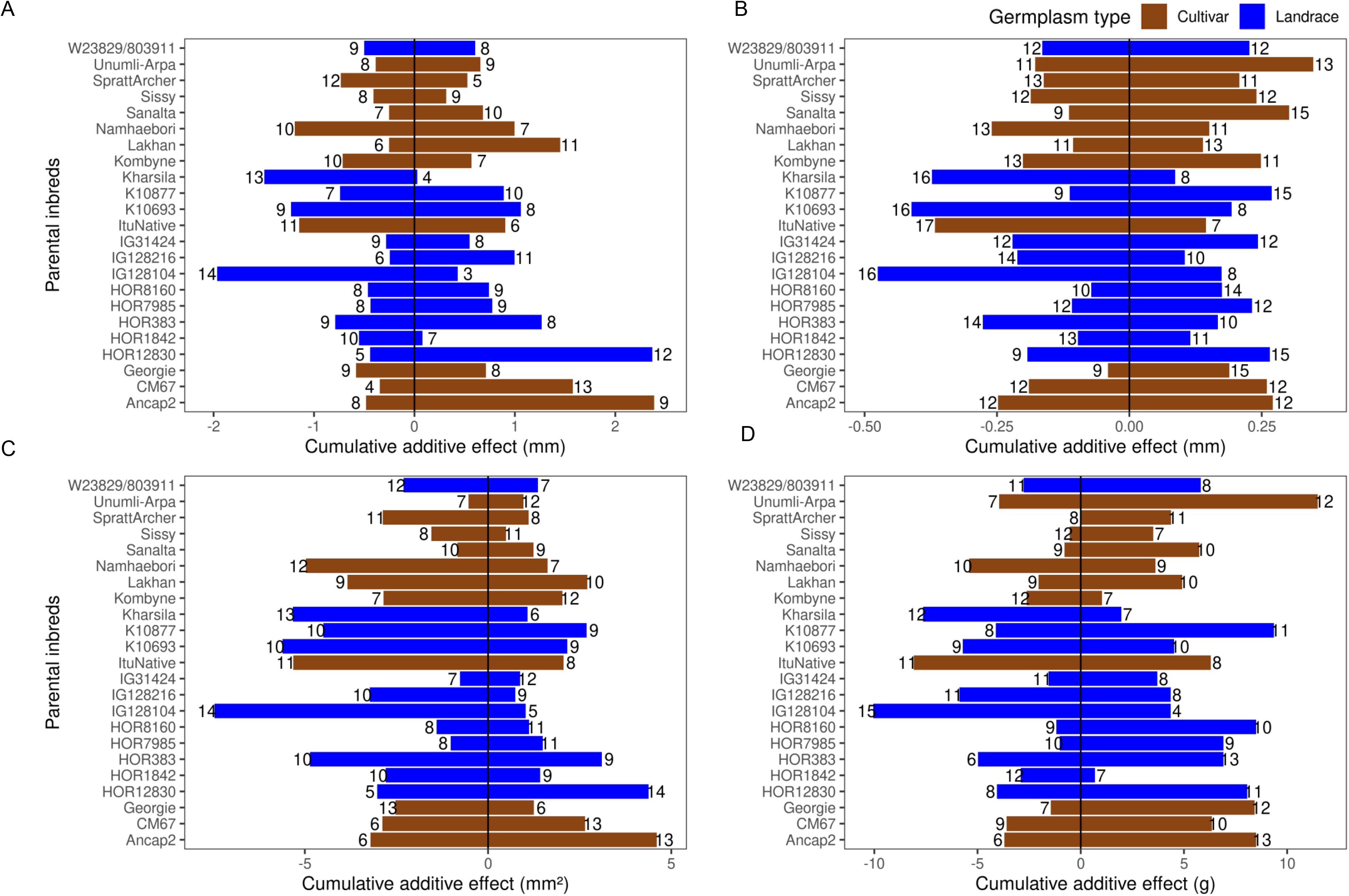
Cumulative allele effect estimate for quantitative trait loci (QTLs) detected in multi-parent population analysis of 45 HvDRR sub-populations using a parental model. Sum of positive (right box) and negative (left box) standardized allele effect size for 23 inbreds stacked across (A) grain length, (B) grain width, (C) grain area and (D) thousand-grain weight QTLs. The cumulative allele effect was the sum of the standardized allele effect for an inbred, where the standardized allele effect for an inbred was the difference between the mean of the estimated allele effects for 23 parental inbreds and the estimated allele effect of the considered inbred. The number next to the bars indicated the count of QTLs with negative or positive standardized allele effect.

The average allele count of positive and negative effects across MPP QTLs was almost the same for landraces and cultivars across all four characters. However, landraces showed higher cumulative negative effects and lower cumulative positive effects than cultivars (Fig. 9). For example, the average allele count for negative allele effect for TGW was 9.1 and 10.2 for landraces and cultivars, respectively. Nevertheless, the average cumulative negative effect for landraces (−4.3 g) was 1.5 fold greater than for cultivars (−2.9 g). However, it was noteworthy that alleles from landraces (HOR12830, K10877, HOR8160 and HOR7985) also contributed to a high genome-wide count of positive allele effects as well as cumulative allele effects (Fig. 9).

### Candidate genes associated with grain size

We obtained SNP annotation, indels and predicted SVs information from the whole-genome sequencing of the 23 parental inbreds (Weisweiler *et al.*, 2022). This information was used to compare the sequence polymorphism between the parental inbreds of HvDRR populations in the coding and regulatory region of the genes present in the confidence intervals of those 19 consensus QTLs that have an interval length < 10 Mbp. The genes present in the consensus QTLs that showed sequence polymorphisms in the coding region and the 5’-end regulatory region between the parental inbreds with distinct allele effects on grain size characters were selected as candidate genes. As a result, we identified 869 candidate genes for these 19 consensus QTLs. The number of candidate genes for the consensus QTLs varied between 6 and 142 (Table S8).

Fourteen consensus intervals harbored the closest barley orthologs of genes controlling grain size in other crops (Table 4). Among them, we identified allelic variations in the regulatory or coding region for eight orthologs. Furthermore, these polymorphisms corresponded to the parental allele effect on QTLs detected across the sub-populations (Additional File 1).

Likewise, we also detected QTLs for grain size in the region associated with lateral spikelet fertility and hull adherence. For example, the consensus QTLs *qHvDRR-GS-14* and *qHvDRR-GS-31* were detected for 2-row by 6-row crosses and harbored *Vrs1 and Vrs5/Int-c*, respectively (Table 4). In addition, we observed allelic variation in the coding sequence or regulatory region between 2-row and 6-row inbreds for the loci mentioned above (Fig. S12 and Additional File 1). Similarly, QTL detected at *Nud* locus (*qHvDRR-GS-61*) was exclusive to populations involving cross between hulled and naked (IG128104 and Kharsila) barley inbreds. Predicted SV indicated the characteristic 17 kb deletion on chromosome 7H (chr7H:529047042-529063730) in naked barley accessions.

*qHvDRR-GS-6* was one of the major effect QTLs for GL that explained with 25-62% a very high proportion of the phenotypic variance in SPA. Further, we validated *qHvDRR-GS-6* in a fine-mapping population derived from the cross of two RILs from the HvDRR33 sub-population carrying Lakhan (long grains) and Georgie (short grains) alleles at this QTL. We found 35 recombinants (bearing different parental alleles at the left and right border of the QTL interval) among 924 F2 progenies. The phenotyping of GL in the F2 population distinguished the two classes of non-recombinants with Lakhan and Georgie allele-bearing progenies producing long and short grains, respectively. The average GL for different groups of recombinants ranged from 8.6 to 9.2 mm (Fig. S13). Although only the initial steps of the positional cloning of the underlying gene for *qHvDRR-GS-6* were done in the current study, the consensus interval harbored genes annotated as carboxypeptidase (HORVU.MOREX.r2.1HG0063410) and TCP transcription factor (HORVU.MOREX.r2.1HG0063230). The variant calling data indicated amino-acid substitution mutations in the above-mentioned candidate genes between the parental inbreds contributing to short and long grains. The protein sequence of both candidate genes was highly conserved among other monocots (Fig. 10, Fig. S14).

**Fig. 10:**
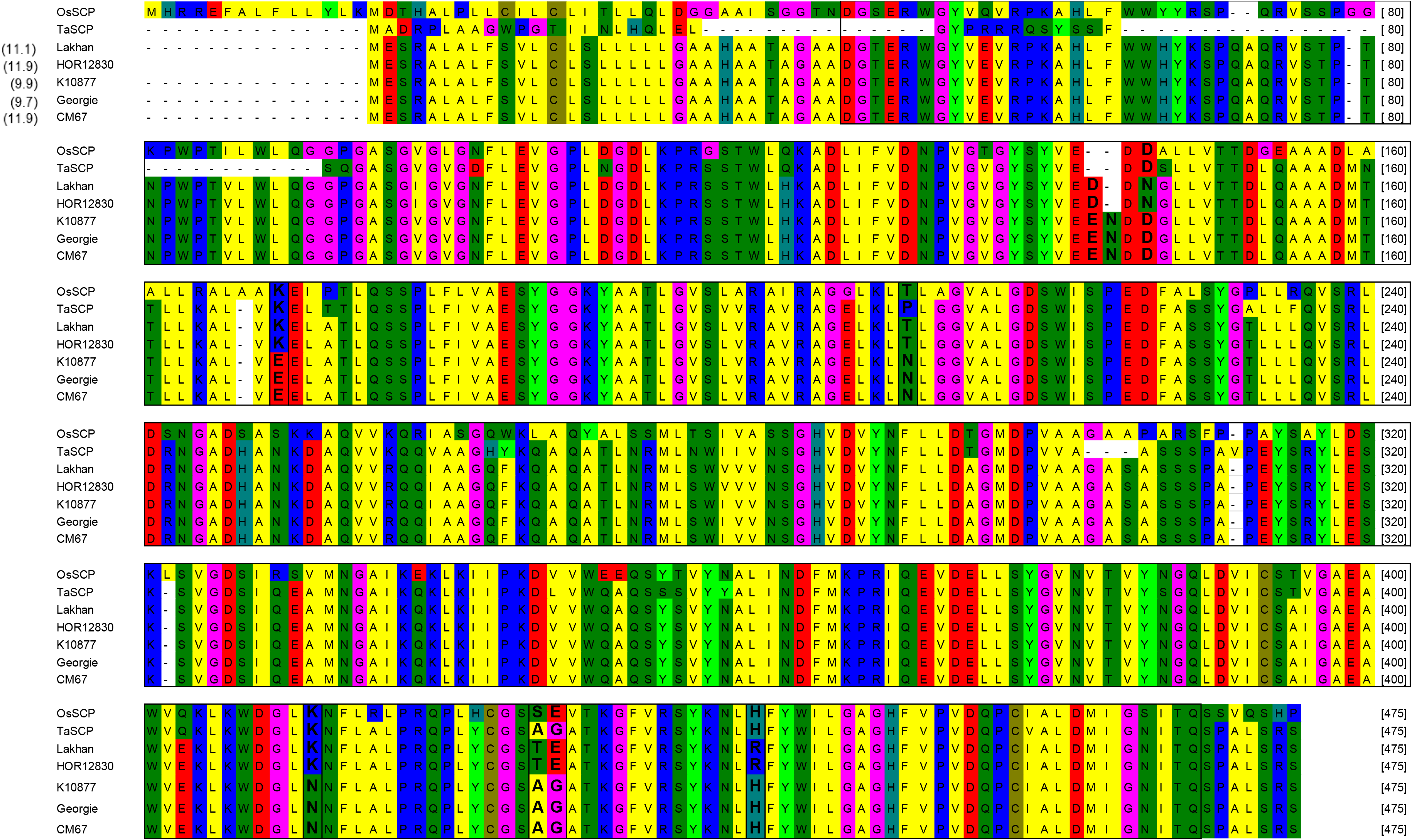
Candidate gene underlying qHvDRR-GS-6 on chromosome 1H. Protein alignment of a candidate gene HORVU.MOREX.r2.1HG0063410. SNP and 3 bp indel caused amino acid substitutions in the conserved carboxypeptidase domain. The protein sequence of the barley gene was highly conserved in rice and wheat. Lakhan and HOR12830 allele at the QTL locus contributed to longer grains. Sites in bold letters indicate amino acid substitutions between parental groups with contrasting allele effects at the qHvDRR-GS-6.

## Discussion

In the present study, we demonstrated the application of the HvDRR population of barley to identify the genetic factors controlling the important yield component characters related to grain size. First, the grain size phenotypes (GL, GW, GA and TGW) were collected in four environments from the RILs of 45 HvDRR sub-populations derived from the genetic crosses between 23 diverse barley inbreds in a double round-robin design. We observed that the broad-sense heritability (*H^2^*) on an entry mean basis was more than 84 % for all characters (Table 1), indicating a substantial genetic variability for grain size in the HvDRR population. In addition to a significant (p < 0.0001) genotypic variance, we also observed significant (p < 0.0001) environment and genotype*environment interaction variances (Table 1). The high genotypic contribution to grain size variation observed in our study was in accordance with earlier studies. With 95% a slightly higher *H^2^* was reported by Wang *et al.* (2019) for grain size characters in a double haploid population of barley. The *H^2^* for GW, GA and TGW observed for the HvDRR population was comparable to that observed for the HEB-25 NAM population grown under adequate nitrogen fertilization. At the same time, that for GL was more than twofold higher in our study (Sharma *et al.,* 2018). The latter observation might be due to that the HEB-25 NAM population was derived from a cross between Barke and wild barley genotypes. In such populations, GL might be overestimated by awn retention due to threshability problems (Schmalenbach *et al.*, 2011), leading to lower heritability.

Although the maximum (4.2 mm) and minimum (2.5 mm) GW observed in RILs of HvDRR population was similar to that of previous reports, we observed a considerably broader range of adjusted entry means for GL, GA and TGW. For instance, the minimum and maximum GL on the adjusted entry mean basis among all RILs was 5.7 and 15.6 mm, respectively. On the other hand, the range for mean GL reported earlier for barley segregating populations was 7.4 to 11 mm (Walker *et al.*, 2013; Zhou *et al.*, 2016; Sharma *et al.*, 2018; Wang *et al.*, 2019; Watt *et al.*, 2019, 2020). The above observation can be explained by the fact that most studies comprised a single bi-parental population and the genetic variance is restricted to two parents. In addition, the parental inbreds of the HvDRR population were selected to maximize genotypic and phenotypic diversity (Weisweiler *et al.*, 2019). Therefore, TGW in the HvDRR population showed about the same range as worldwide spring barley (Pasam *et al.*, 2012). These observations support the results of earlier simulation studies (Stich, 2009; Klasen *et al.*, 2012), that partial diallel designs as implemented in our study capture more allelic diversity than NAM type populations, thus, providing high power for unraveling the genetics of natural phenotypic variation.

### Genetic complexity of grain size variation

We performed MPP and SPA analyses and detected 79 and 316 grain size and weight QTLs, respectively (Fig. 3-7). In general, the LOD scores of QTLs detected in MPP analysis were higher than SPA, which suggested an increased QTL detection power in MPP analysis (Garin *et al.*, 2021). However, 56 SPA QTLs were not detected at the same positions as MPP QTLs (Fig. 4-7) because the rare allele with a small effect detected in specific sub-populations might not pass the significant threshold in MPP analysis. Therefore, as well as due to the potential negative effect of imprecisions in the consensus map, we considered the results of both MPP and SPA in our study.

In order to do so, we summarized the SPA QTLs into 62 consensus QTLs, the shortest possible confidence interval of co-located QTLs across sub-populations or characters (Table 4). The number of consensus QTLs associated with grain size and weight in the HvDRR population was considerably higher than in previous reports (Table 4). For instance, in our study, 42 and 36 consensus QTLs were linked to GA and TGW (Table 4). In contrast, only 7 and 11 GA QTLs were detected in the study of Wang *et al.* (2019) and Sharma *et al.* (2018), respectively. Xu *et al.* (2018) identified 21 GA-associated QTLs, which was half of the number detected in the HvDRR population. The same trend was observed for GL, GW and TGW. When comparing the physical interval of the consensus QTLs with the QTLs detected in the previous linkage and association mapping studies concerning grain size and weight (Ayoub *et al.*, 2002; Pasam *et al.*, 2012; Walker *et al.*, 2013; Pauli *et al.*, 2014; Maurer *et al.*, 2016; Sharma *et al.*, 2018; Xu *et al.*, 2018; Wang *et al.*, 2019; Watt *et al.*, 2019, 2020), 13 consensus QTLs for grain size characters detected in our study have not been reported earlier. These findings suggest that grain characters in barley are more complexly inherited than reported previously.

In addition, we observed that the prediction ability of the genomic selection model was about 0.92 (Table 3), which was 20% higher than the comparable figure for a QTL based prediction, i.e., the square root of the proportion of the explained phenotypic variance of the full additive model of MPP QTLs (Table 2). This finding suggests that even with a big mapping population like ours, the effect of many more minor effect loci did not pass the significant threshold in MPP analyses. Thus, the real genetic complexity of grain size characters is even more complex than discussed above.

However, despite the high genetic complexity of the studied characters across all HvDRR populations, we have observed that the majority (approx. 0.6 frequency) of the SPA QTLs explained 10 to 20 % of the phenotypic variance also after cross-validation (data not shown) (Fig. S7). This finding indicates that QTL fine mapping and ultimately map-based cloning of the underlying genes will be possible for a considerable proportion of the detected QTL when the most relevant sub-populations are studied, which identification was one of the objectives of this study.

### Previously described QTLs for grain parameters were detected in the HvDRR population

We detected major effect genes associated with spikelet architecture and hulled caryopsis in the HvDRR population as a proof of concept (Table 4, Fig. S13, Table S8). For example, the consensus QTL *qHvDRR-GS-18* was detected in six 2 vs. 6-row sub-populations and harbored *Vrs1* (Table 4). Sequence comparisons of the parental inbred lines revealed that six-rowed and two-rowed barley carried *vrs1* and *Vrs1*, respectively (Fig. S12). *Vrs1* (homeobox domain-containing protein) is the major gene determining the lateral spikelet fertility in barley and has a pleiotropic effect on grain size (Komatsuda *et al.*, 2007). Therefore, the bigger and heavier grain in 2-row genotypes may compensate for fewer seeds compared to 6-row genotypes (Ayoub *et al.*, 2002).

Likewise, *qHvDRR-GS-61* was detected at the *Nud* locus in HvDRR populations involving naked and hulled barley (Table 4). This observation indicated a pleiotropic effect of naked caryopsis on grain size and weight which was also reported by Wang *et al.* (2019) in a mapping population developed from naked and hulled barley. Parental inbreds of the HvDRR population with the non-adhering hull (Kharsila, IG128104) revealed a 17 Kb deletion (chr7H:529047042-529063730), resulting in the null allele for an ethylene response factor gene which is characteristic of naked barley genotypes (Taketa *et al.*, 2008).

Furthermore, we confirmed the QTL allele effect in a high-resolution population segregating for one of the novel GL QTLs (*qHvDRR-GS-6*) located on chromosome 1H (Fig. S13). These examples illustrated the accuracy and relevance of the detected loci in our study and suggested that the detailed consideration of the other detected QTL provides biologically relevant information.

### The function of some grain size-related genes might be conserved in cereals

We utilized the single nucleotide variant calling data, indels and predicted SVs of the 23 parental inbreds in coding and potentially regulatory regions (Weisweiler *et al.*, 2022), and their functional annotation to describe the underlying candidate genes (Table S8, Additional File 1). We detected polymorphisms in the coding region among the contrasting parental inbreds for the genes present in the consensus QTLs. For instance, the barley ortholog of *GS5* (*HvGS5*) was the candidate for *qHvDRR-GS-24* associated with all four evaluated grain characters (Table S8). *GS5* encodes a serine carboxypeptidase protein and has been described as positive grain size and weight regulator in rice and wheat (Li *et al.*, 2011; Ma *et al.*, 2016; Wang *et al.*, 2016). Another gene (HORVU.MOREX.r2.1HG0063410) that is closely related to *HvGS5* was the most plausible candidate for *qHvDRR-GS6* (Table S8).

In addition, *qHvDRR-GS6* harbored a barley gene (HORVU.MOREX.r2.1HG0063230) which is closely related to *TCP5* like transcription factor family proteins. The latter regulates leaf and petal size in *A. thaliana* (van Es *et al.*, 2018). We detected amino acid substitutions or indels in HORVU.MOREX.r2.1HG0063410 and HORVU.MOREX.r2.1HG0063230 between parental alleles with long (Lakhan and HOR12830) and short grains (Georgie, K10877 and CM67) at *qHvDRR-GS6* (Fig. 1 and Table S8). We further validated the non-synonymous mutations and exon indels identified from the whole-genome sequencing using Sanger sequencing of both genes. These results indicated the accuracy of the variant calling file used to select the candidate gene in our study (Fig. 10, Fig. S14). The above genes should be targeted for functional validation studies.

The barley ortholog for *TaCYP78A3* (Ma *et al.*, 2015a) encoding cytochrome P450 (*HvCYP78A*) family protein was the candidate gene for *qHvDRR-GS-58*. Similarly, a gene closely related to *HvCYP78A* on chromosome 2H was the most promising candidate for *qHvDRR-GS-11* (Table S8). Cytochrome P450 is a large protein family involved in different biosynthetic processes (Li *et al.*, 2019). For example, the CYP78A, one of the cytochrome P450 families, is a positive regulator of seed size in wheat and *A. thaliana* (Nagasawa *et al.*, 2013; Ma *et al.*, 2015a). Likewise, *HvGSK2* (*q-HvDRR-GS-3*), *HvWG7* (*q-HvDRR-GS-14*), *HvSRS5* (*q-HvDRR-GS-33*), *HvLGY3* (*q-HvDRR-GS-36*) and *HvRGA1* (*q-HvDRR-GS-55*) were the other barley orthologs of grain size associated genes from rice detected for the respective QTLs (Table S8 and Additional File 1).

HORVU.MOREX.r2.5HG0421190, a gene encoding E3 ubiquitin-protein ligase might be the causal gene for *qHvDRR-GS-43*. It revealed amino acid substitutions among the parental lines (Table S8). *GW2* in rice encodes an E3 ubiquitin-protein ligase involved in the ubiquitin-proteasome signaling pathway (Song *et al.*, 2007). It is the negative regulator of grain size and the function is conserved in monocots, including rice, wheat and maize (Song *et al.*, 2007; Li *et al.*, 2010a; Simmonds *et al.*, 2016). Genes encoding strictosidine synthase were among the candidates for *qHvDRR-GS-30* (HORVU.MOREX.r2.4HG0276930) and *qHvDRR-GS-44* (HORVU.MOREX.r2.5HG0424360). Strictosidine synthase showed indole-3-acetic acid glucose hydrolase activity and negatively regulated TGW in rice (Ishimaru *et al.*, 2013). We also detected genes coding for the bi-directional sugar transporter SWEET (HORVU.MOREX.r2.6HG0521560) as the candidate gene for *qHv-DRR-54* on chromosome 6H (Table S8). The SWEET family protein is involved in seed filling in *A. thaliana*. Triple knockout mutant SWEET family protein (*sweet11;12;15*) in *A. thaliana* produced wrinkled seeds with reduced weight (Zhou *et al.*, 2015). Different classes of zinc finger proteins, receptor kinases and transcription factors (MADS-box, MYB) were the other categories of highly represented candidate genes that are involved in seed development. In conclusion, the candidate gene analysis identified genes that are known to be involved in cell proliferation, cell elongation and grain filling, which ultimately determines grain size and weight

### Sink size primarily determines grain weight in the HvDRR population

The final grain weight is determined by the sink size, i.e. the physical dimension of the grain and grain filling (Li *et al.*, 2018). We measured GL, GW and GA as a proxy of sink size in our study. Across the HvDRR population, the correlation between these characters and TGW was up to r = 0.84 (Fig. 2). This suggests that variability in grain filling can only explain a low additional proportion of the variability of grain weight and that sink size is the primary determinant of grain weight in the HvDRR population. In addition, of 62 consensus QTLs detected in our study, 14 and 11 loci were associated exclusively with GL and GW, respectively but not with TGW (Table 4). Likewise, 35 consensus QTL were linked to TGW and only three were not associated with any of the grain size parameters (Table 4). This supported our above conclusion that sink size was the major determinant of grain weight in the HvDRR population. Our observation agrees with Bingham *et al.* (2007) findings that the mean grain weight of field-grown barley was strongly determined by potential grain size. The positive correlation between grain size and grain weight observed in our study was furthermore in accordance with other crops (Li *et al.*, 2019). For example, genes and genetic loci controlling GW also enhanced grain weight in rice (Song *et al.*, 2007; Shomura *et al.*, 2008; Li *et al.*, 2011). Likewise, the GL QTLs in rice were associated with TGW and long grains were heavier than short grains. It has also been shown that grain filling and grain size are regulated by the same or closely related genes in rice (Wang *et al.*, 2008; Yan *et al.*, 2011).

Similar to previous studies (Sharma *et al.*, 2018; Xu *et al.*, 2018; Wang *et al.*, 2019), we observed a stronger correlation between TGW and GW than between TGW and GL in the HvDRR population (Fig. 2). This might be explained by the order in which GW and GL develop. It has been shown in wheat that GL reaches its maximum at the initial grain filling period, whereas GW reaches a maximum at the late grain filling period. Another explanation might be that the increase in grain volume occurs in the transverse plane during the rapid grain filling stage, which will more directly affect GW than GL (Xie *et al.*, 2015). Due to the genetic relatedness between barley and wheat, the change in grain dimension post-anthesis might be similar, explaining a better correlation between TGW and GW compared to TGW and GL, which warrants further investigation.

Across the entire HvDRR population, a significant positive correlation between GL and GW was observed (Fig. 2). However, in agreement with earlier studies (Sharma *et al.*, 2018; Watt *et al.,* 2019,2020), in seven sub-populations (HvDRR13, HvDRR14, HvDRR18, HvDRR19, HvDRR22, HvDRR26, and HvDRR32), we observed a particularly weak correlation between GL and GW (Fig. S2). This observation suggested that GL and GW are at least partially genetically independent. The QTL detected in SPA of the above-mentioned sub-populations for GL did not overlap with those for GW (Fig. 4-5, Table S3-S4), suggesting that different genes contribute to their variation. GL associated QTLs without effect on GW and vice versa were also observed in other cereals such as rice (Yin *et al.*, 2020). Across 45 sub-populations, a total of 15 consensus QTLs were detected for both GL and GW (Table 4). Among them, the allele effect was in the same direction for GL and GW for 13 QTLs (Table S3-S4). This trend was also reported for most of the grain size QTL in rice (Song *et al.*, 2007; Che *et al.*, 2015; Xiao *et al.*, 2019; Zhong *et al.*, 2020) and wheat (Bednarek *et al.*, 2012; Ma *et al.*, 2015b; Liu *et al.*, 2020a). For two of the QTL, namely *qHvDRR-GS-45* and *qHvDRR-GS-52*, the QTL effects revealed an opposite direction for GL and GW (Table 4, Table S3-S4). *qHvDRR-GS-45* and *qHvDRR-GS-52* did not harbor the orthologs of rice genes for GS9 (Zhao *et al.*, 2018) and GW5 (Li *et al.*, 2011) that regulate the development of short and wide grains. Therefore, identifying the underlying genes in barley is particularly of interest to maximize yield.

### Beneficial alleles for grain size in barley landraces

The favorable allele for increased grain size and uniform shape is accumulated in elite germplasm because of directional selection during the domestication process (Gegas *et al.*, 2010; Zhang *et al.*, 2021). However, landraces that evolved from natural and human selection might still be a significant source of positive genetic diversity (Casañas *et al.*, 2017). To this date, linkage mapping studies for grain size in barley were limited to genetic crosses between cultivars or cultivars and wild relatives (Zhou *et al.*, 2016; Sharma *et al.*, 2018; Wang *et al.*, 2019, 2021; Watt *et al.*, 2019, 2020). The parental inbreds of the HvDRR population, however, comprised 12 barley landraces and 11 cultivars. We compared the allelic count and the cumulative allele effect across QTLs detected in MPP analysis between landraces and cultivars. The net allelic count with the positive effect size for grain size and weight QTLs detected from MPP analysis was slightly higher for cultivars (54%) than for landraces (46%). In addition, the net cumulative allele effect (difference between cumulative positive and negative effects) was slightly higher for cultivars than for landraces (Fig. 9). This finding of a higher proportion of alleles from cultivars contributing to an increase in grain size and weight indicates that bigger and heavier grains were positively selected in barley in the recent breeding history. This is in agreement with the findings for rice that for most of the cloned grain size QTLs such as *GS3, GW5, GW2, GW8, GW7, TGW3* and *TGW6* the wild-type genes were the negative regulator of grain size (Li *et al.*, 2019).

However, the result of the allelic series analysis across the MPP QTL implied that all parental inbreds contributed positive and negative allele effects (Fig. S8-11). This finding was in accordance with the results of Sharma *et al.* (2018), who reported that the effect of 25 wild barley alleles segregating in cultivated barley showed both positive and negative allele effects across the QTL hotspots for different grain size characters. Nevertheless, within a single QTL locus, the effect of all wild barley alleles was primarily showing the same direction. In contrast, we identified statistically (p < 0.05) distinct groups of parental inbreds with a differing magnitude of allele effect across the respective QTL locus (Fig. S8-11) regardless of being cultivars or landraces. This is particularly advantageous because HvDRR sub-populations involving parental inbreds on the extreme of the allelic series at a QTL might be ideal for fine-mapping of the locus. The allelic series data also indicated that landraces such as K10877, HOR12830, HOR8160 and HOR7985 made significant contributions to favorable alleles across the grain size and weight QTLs (Fig. 9). For instance, the allelic series comparison at MPP QTLs revealed that landraces contributed higher positive effects at *qGL-Hr1-1, qGL-Hr7-2, qGW-Hr5-4, qGW-Hr7-3, qTGW-Hr5-1, and qTGW-Hr5-2* than did cultivars (Fig. S8, S9 and S11). This observation is in accordance to the finding that some domestication alleles of grain size genes in rice (Liu *et al.*, 2017a), maize (Li *et al.*, 2010b) and wheat (Liu *et al.*, 2020a) were detected in the primary gene pool.

## Conclusion

The results of this study indicate that the genetic architecture of grain size is more complex than reported previously. Although the grain size and weight phenotypic variation were mainly controlled by the additive effect of minor and moderate effect alleles, we also detected a few major effect alleles for all four grain characters. Allelic series across the MPP QTLs indicated that all 23 parental inbreds have positive and negative allele effects on the grain size and weight. The net cumulative allele effect was only slightly higher for cultivars than landraces illustrating the potential of the latter for breeding projects. We utilized the whole genome sequencing data of the 23 parental inbreds and identified several promising candidate genes. The future work will focus on the validation of these candidate genes and the performance of yield trials using RILs that carry the alleles with a positive allele effect. This study demonstrates the utility of the HvDRR population for dissecting the genetic architecture of complex characters and welcomes the scientific community to exploit the resources.

## Authors contribution

BS: conceptualized the study, acquired funding and supervised the work. AS, FCo, PW and BS: analyzed the data; DVI: performed genomic selection and interpreted the results; JL: created the population; FCa and DVI: prepared genetic maps as well as designed and implemented field trials; MW: prepared variant calling file; AS: created and genotyped the high-resolution population; AS and BS wrote the manuscript. DVI, MW and PW: corrected the manuscript. All authors read and approved the manuscript.

## Acknowledgment

The authors would like to thank Anja Kyriacidis, Florian Esser and George Alskief for managing the field trials. We also acknowledge the computational platform provided by the Centre of Information and Media Technology at Heinrich-Heine-University Düsseldorf. Also, thanks to Yagmur Sahbaz, Kim Schröder and Fabian Krautwald for collecting the data using MARViN seed analyzer.

## Abbreviations

DRR: double-round robin
GA: grain area
GL: grain length
GW: grain width
MPP: multi parent population
RIL: recombinant inbred line
SPA: single population analysis
SV: structural variant
QTL: quantitative trait locus
TGW: thousand-grain weight

---

**Table S1:** Broad-sense heritability of grain size-related traits. Phenotypic data were collected from the grains harvested from four environments. The phenotypic data were adjusted by correcting the field heterogeneity at each environment based on the field map of the augmented design using R package mvngGrAd. The moving plot across the field was constructed using 3, 5, 10 and 15 adjacent rows in a quadratic design for field heterogeneity correction.

| Trait | No field<br>heterogeneity<br>adjustment | Adjustment 3 | Adjustment 5 | Adjustment 10 | Adjustment 15 |
| --- | --- | --- | --- | --- | --- |
| Grain length | 0.84 | 0.71 | 0.74 | 0.79 | 0.81 |
| Grain width | 0.84 | 0.73 | 0.75 | 0.80 | 0.82 |
| Grain area | 0.89 | 0.78 | 0.81 | 0.85 | 0.87 |
| 1000 grain weight | 0.85 | 0.67 | 0.71 | 0.77 | 0.80 |

**Table S2:**
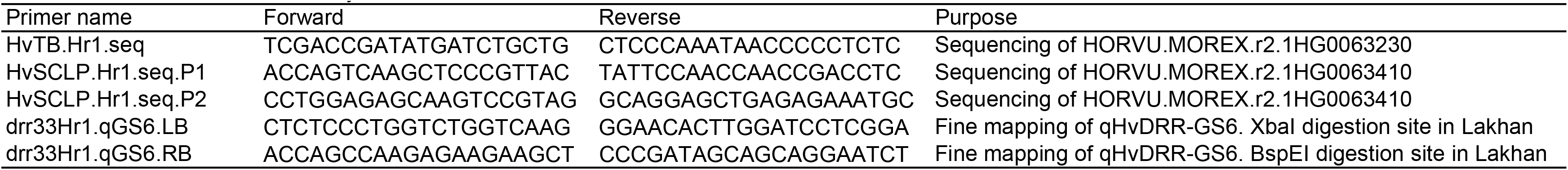
Primers used in the study

**Table S3:**
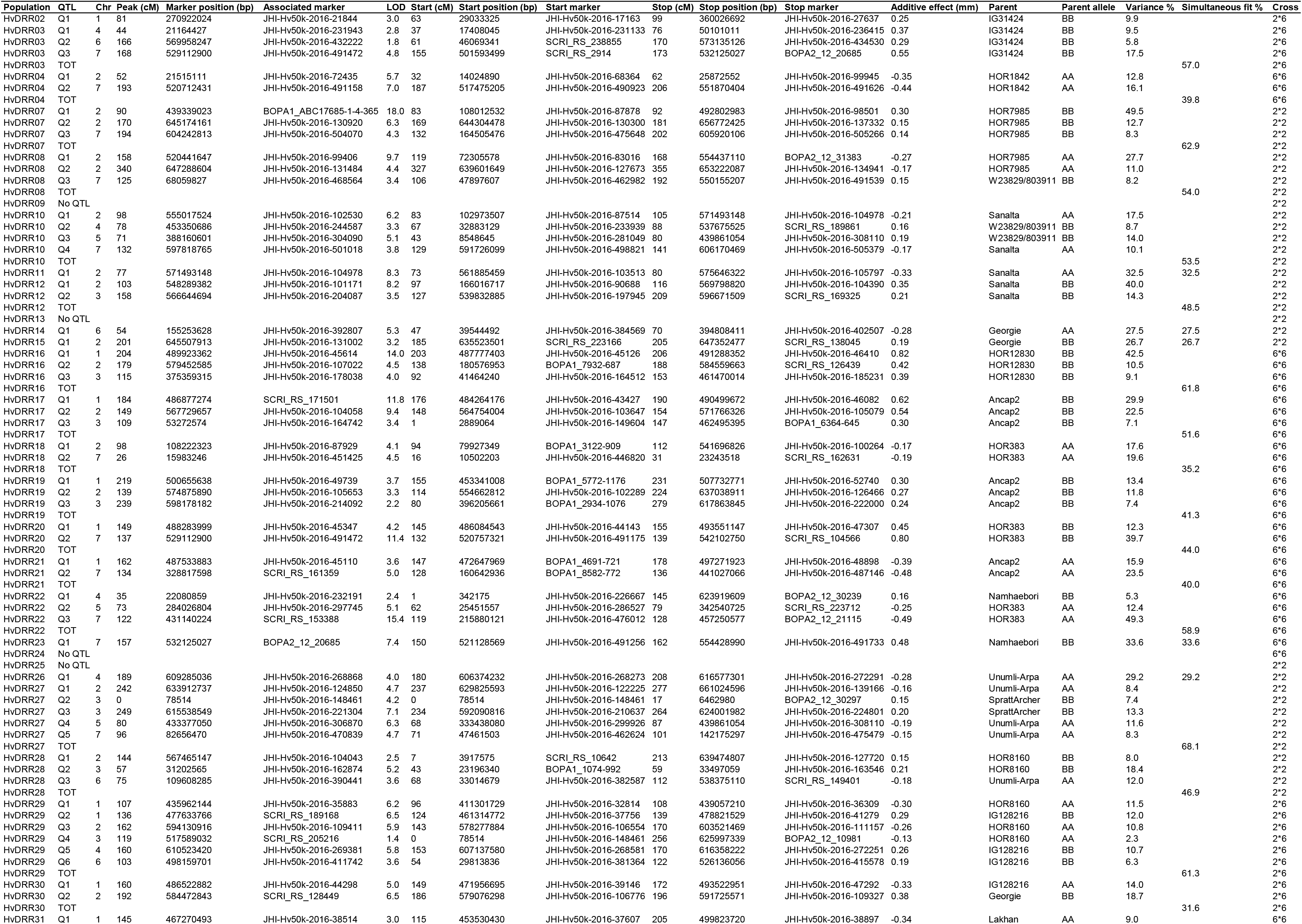
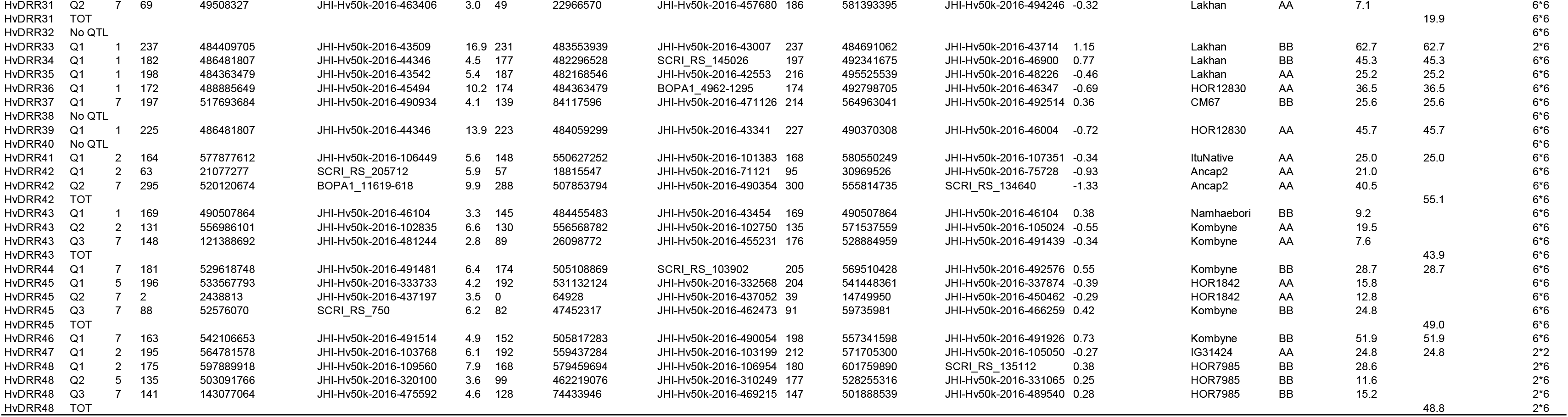
Summary of quantitative trait loci (QTLs) detected in HvDRR sub-populations for grain length. Composite interval mapping was performed using the Rqtl package. The genetic and physical position of peak markers, the markers flanking the confidence interval of the QTL and the percentage of explained phenotypic variation are reported. Parent and parental alleles contributing to positive additive effect to grain length are indicated in the table. Cross denotes the row type of parents used to develop the corresponding recombinant inbred line populations.

**Table S4:**
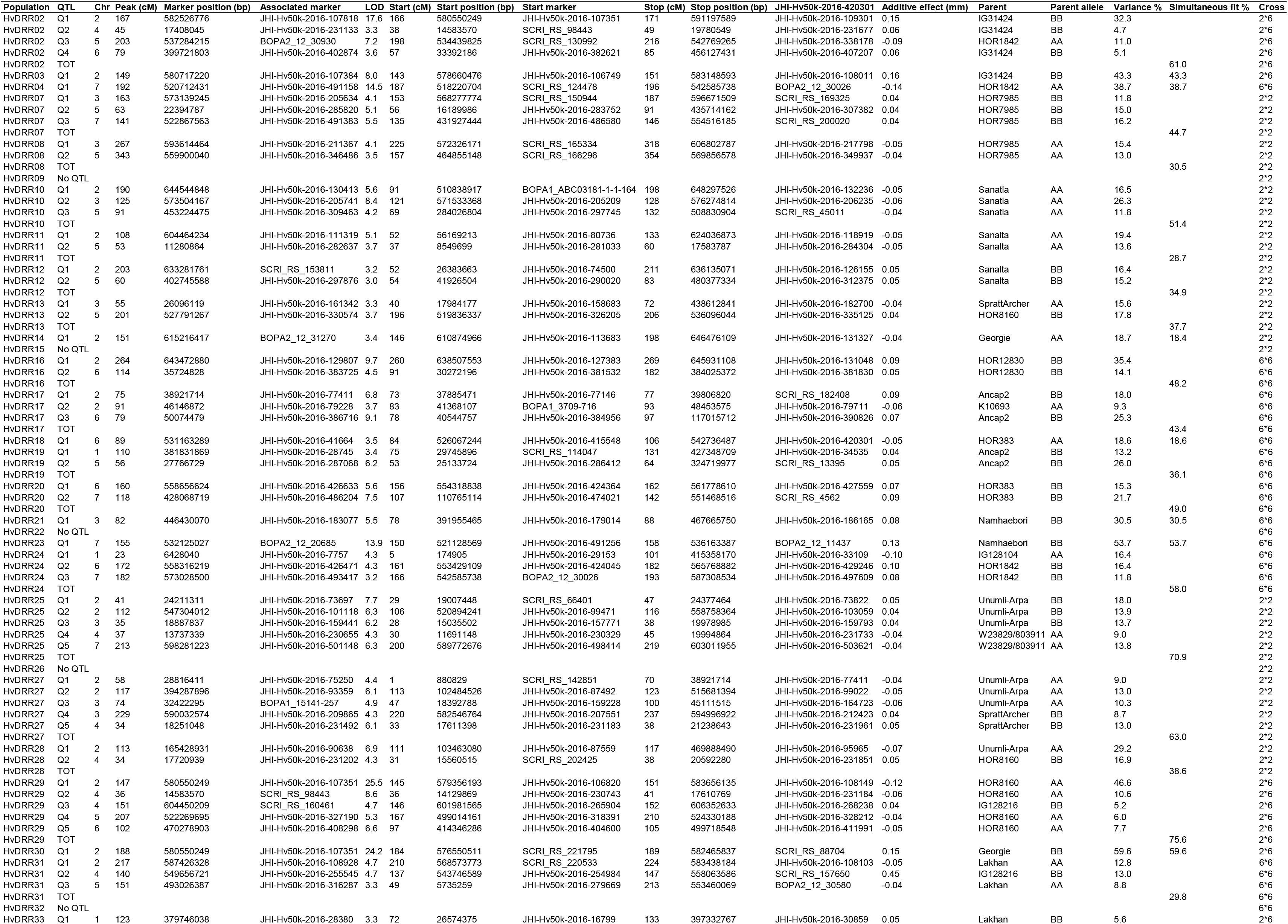
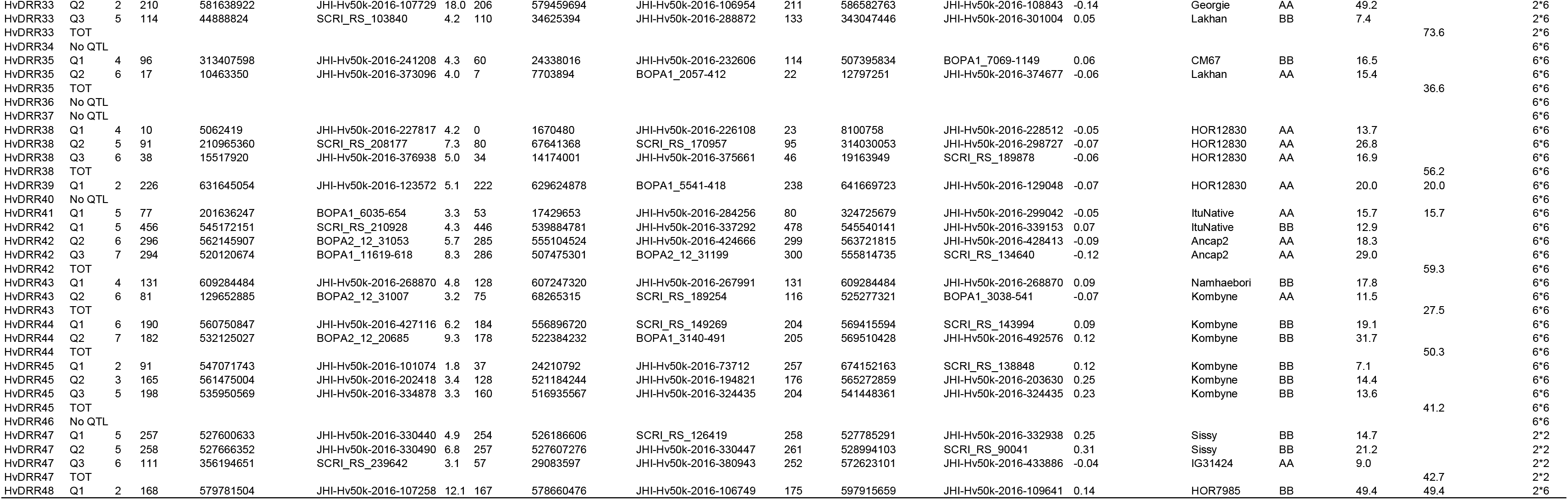
Summary of quantitative trait loci (QTLs) detected in HvDRR sub-populations for grain width. Composite interval mapping was performed using the Rqtl package. The genetic and physical position of peak markers, the markers flanking the confidence interval of the QTL and the percentage of explained phenotypic variation are reported. Parent and parental alleles contributing to positive additive effect to grain width are indicated in the table. Cross denotes the row type of parents used to develop the corresponding recombinant inbred line populations.

**Table S5:**
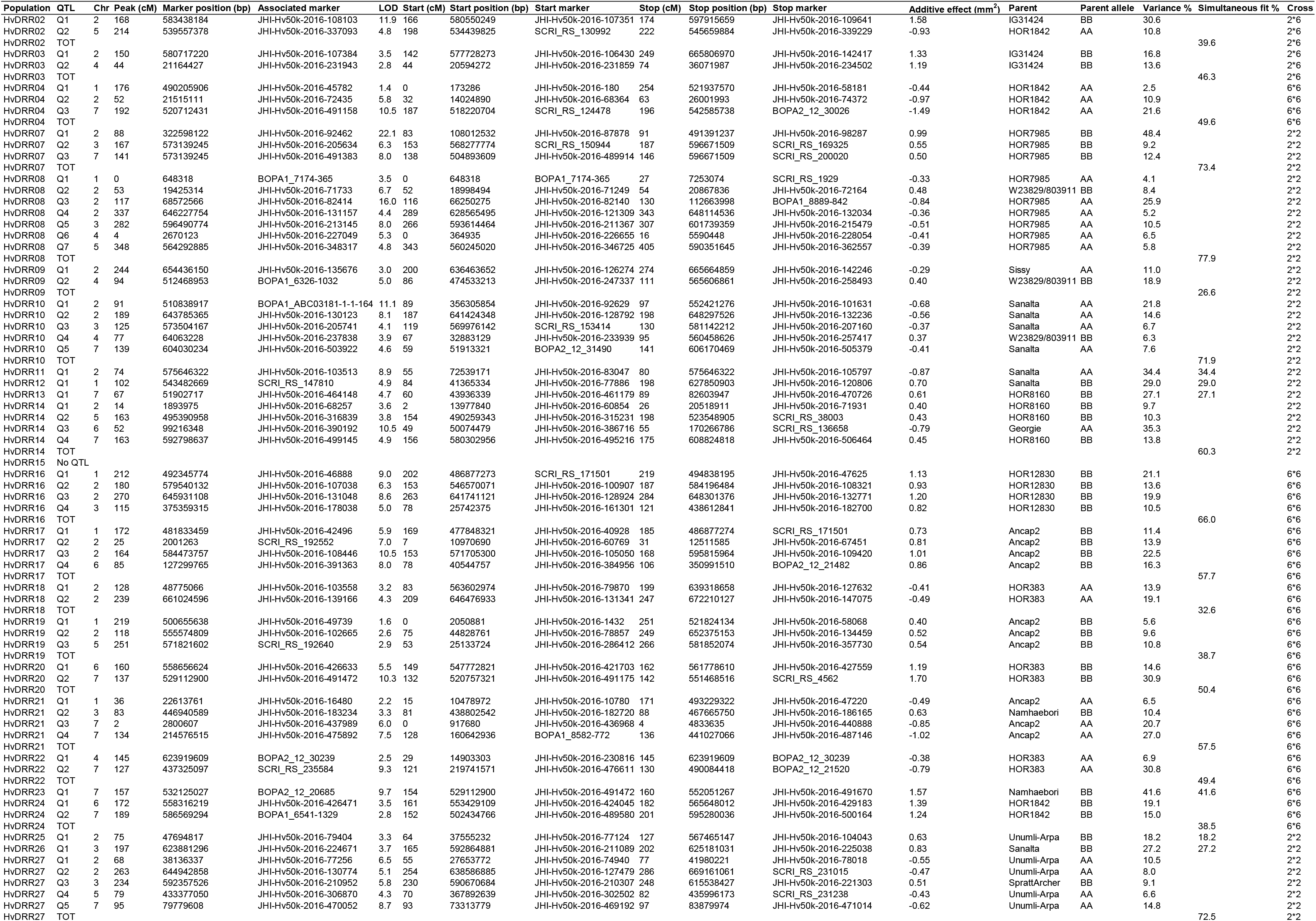
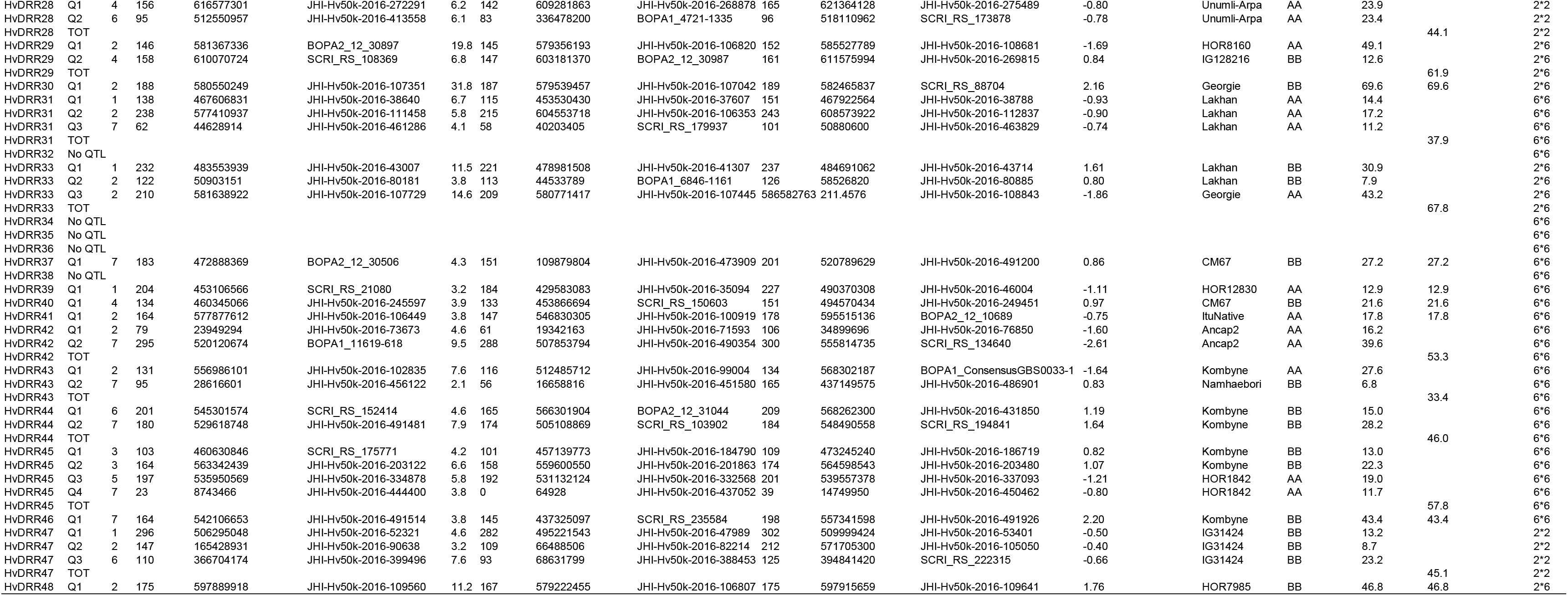
Summary of quantitative trait loci (QTLs) detected in HvDRR sub-populations for grain area. Composite interval mapping was performed using the Rqtl package. The genetic and physical position of peak markers, the markers flanking the confidence interval of the QTL and the percentage of explained phenotypic variation are reported. Parent and parental alleles contributing to positive additive effect to grain area are indicated in the table. Cross denotes the row type of parents used to develop the corresponding recombinant inbred line populations.

**Table S6:**
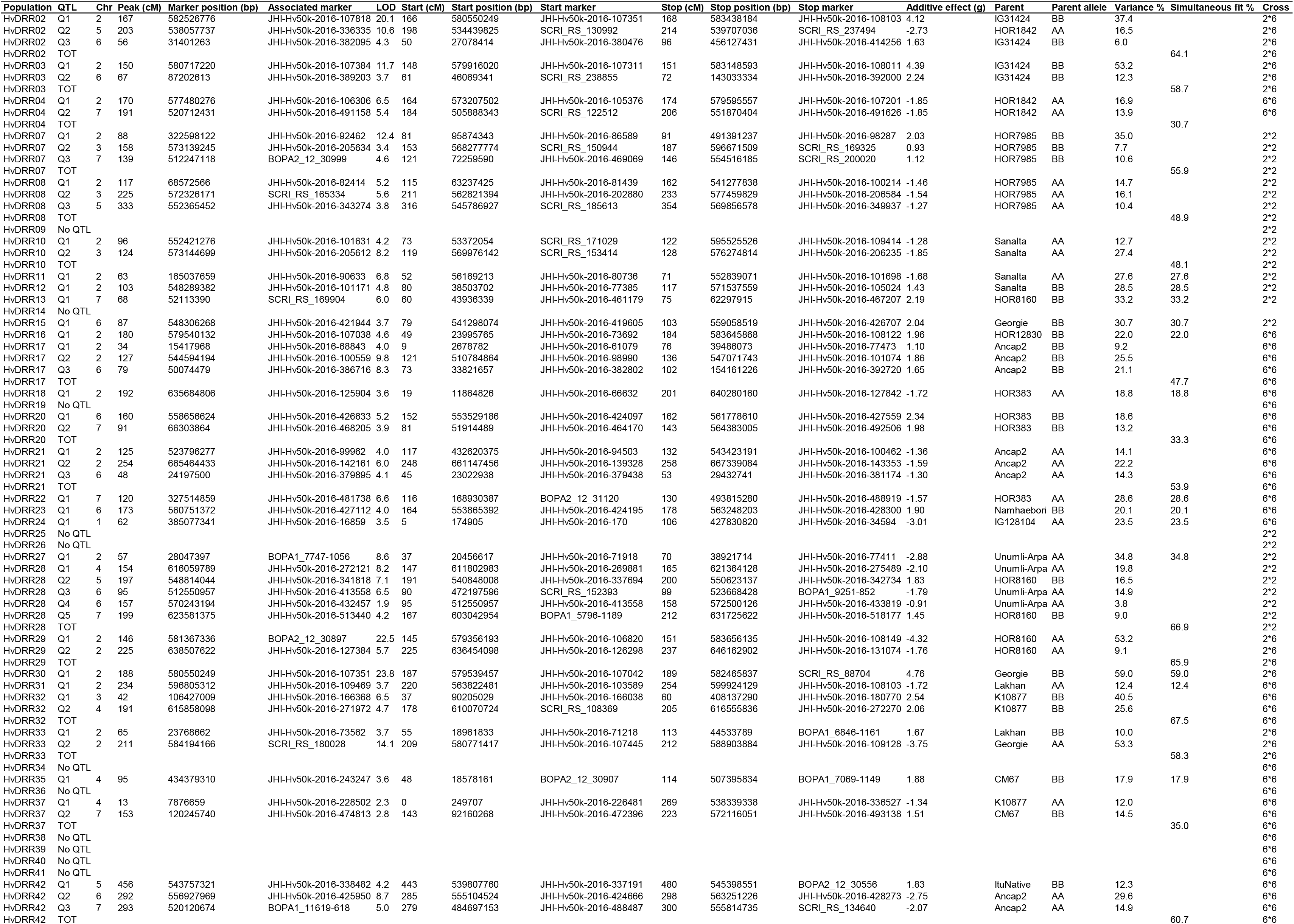
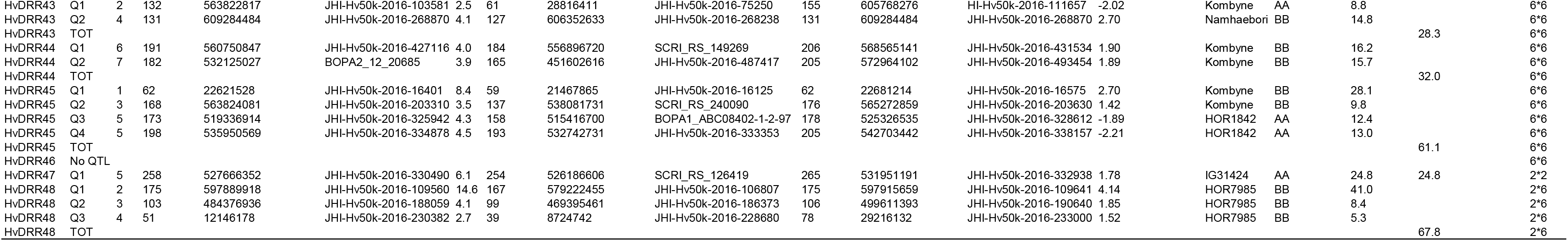
Summary of quantitative trait loci (QTLs) detected in HvDRR sub-populations for thousand grain weight. Composite interval mapping was performed using the Rqtl package. The genetic and physical position of peak markers, the markers flanking the confidence interval of the QTL and the percentage of explained phenotypic variation are reported. Parent and parental alleles contributing to positive additive effect to thousand grain weight are indicated in the table. Cross denotes the row type of parents used to develop the corresponding recombinant inbred line populations.

**Table S7:**
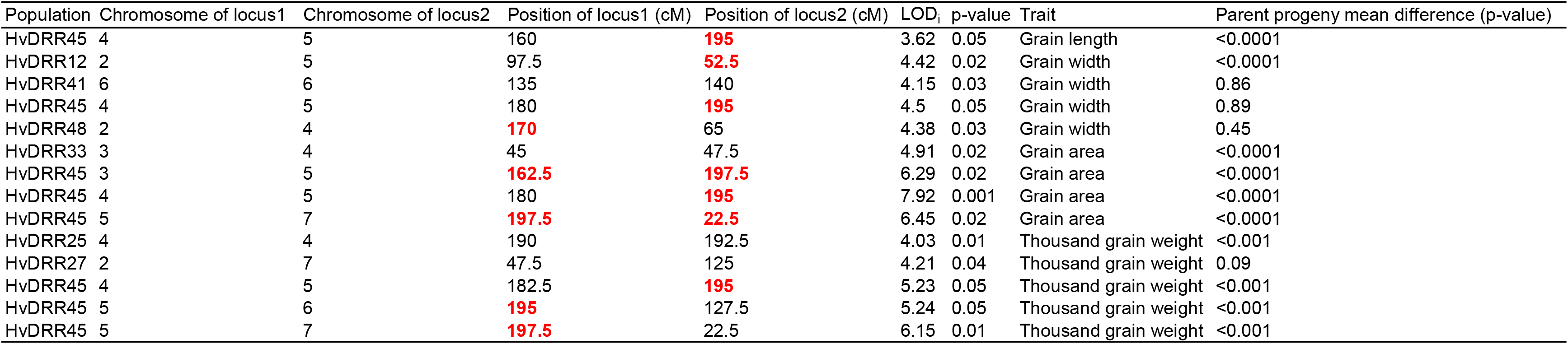
Summary of genome-wide epistatic loci detected in HvDRR population. A two-dimensional genome scan with a two-QTL model, allowing for possible interactions was performed using the Rqtl package. The genetic positions of interacting loci are reported. The loci indicated in bold red letters were also detected as the main effect quantitative trait locus (QTL) from interval mapping.

| Population | Chromosome of locus1 | Chromosome of locus2 | Position of locus1 (cM) | Position of locus2 (cM) | LOD | p-value | Trait | Parent progeny mean difference (p-value) |
| --- | --- | --- | --- | --- | --- | --- | --- | --- |
| HvDRR45 | 4 | 5 | 160 | **195** | 3.62 | 0.05 | Grain length | <0.0001 |
| HvDRR12 | 2 | 5 | 97.5 | **52.5** | 4.42 | 0.02 | Grain width | <0.0001 |
| HvDRR41 | 6 | 6 | 135 | 140 | 4.15 | 0.03 | Grain width | 0.86 |
| HvDRR45 | 4 | 5 | 180 | **195** | 4.5 | 0.05 | Grain width | 0.89 |
| HvDRR48 | 2 | 4 | **170** | 65 | 4.38 | 0.03 | Grain width | 0.45 |
| HvDRR33 | 3 | 4 | 45 | 47.5 | 4.91 | 0.02 | Grain area | <0.0001 |
| HvDRR45 | 3 | 5 | **162.5** | **197.5** | 6.29 | 0.02 | Grain area | <0.0001 |
| HvDRR45 | 4 | 5 | 180 | **195** | 7.92 | 0.001 | Grain area | <0.0001 |
| HvDRR45 | 5 | 7 | **197.5** | **22.5** | 6.45 | 0.02 | Grain area | <0.0001 |
| HvDRR25 | 4 | 4 | 190 | 192.5 | 4.03 | 0.01 | Thousand grain weight | <0.001 |
| HvDRR27 | 2 | 7 | 47.5 | 125 | 4.21 | 0.04 | Thousand grain weight | 0.09 |
| HvDRR45 | 4 | 5 | 182.5 | **195** | 5.23 | 0.05 | Thousand grain weight | <0.001 |
| HvDRR45 | 5 | 6 | **195** | 127.5 | 5.24 | 0.05 | Thousand grain weight | <0.001 |
| HvDRR45 | 5 | 7 | **197.5** | 22.5 | 6.15 | 0.01 | Thousand grain weight | <0.001 |

**Table S8:**
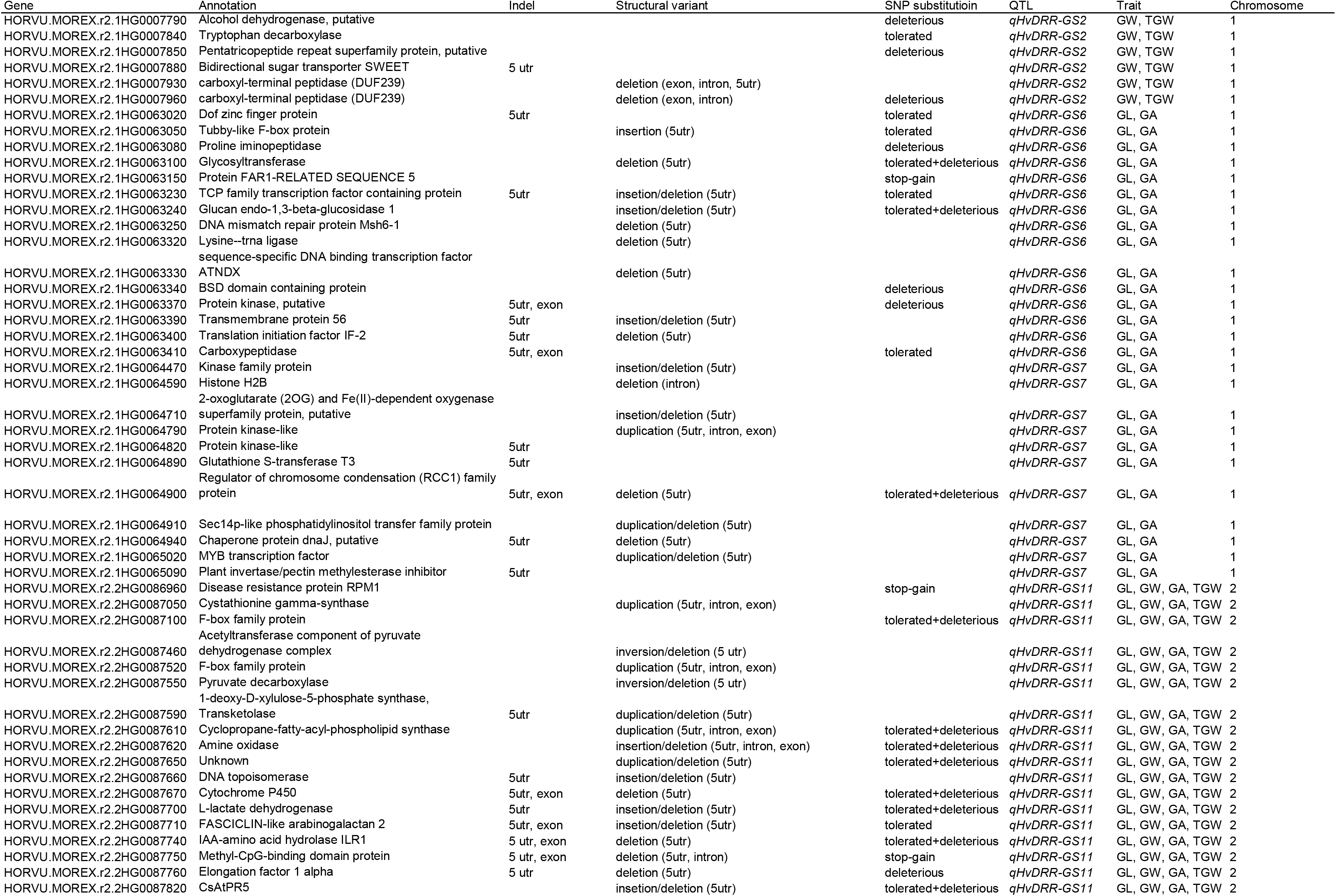
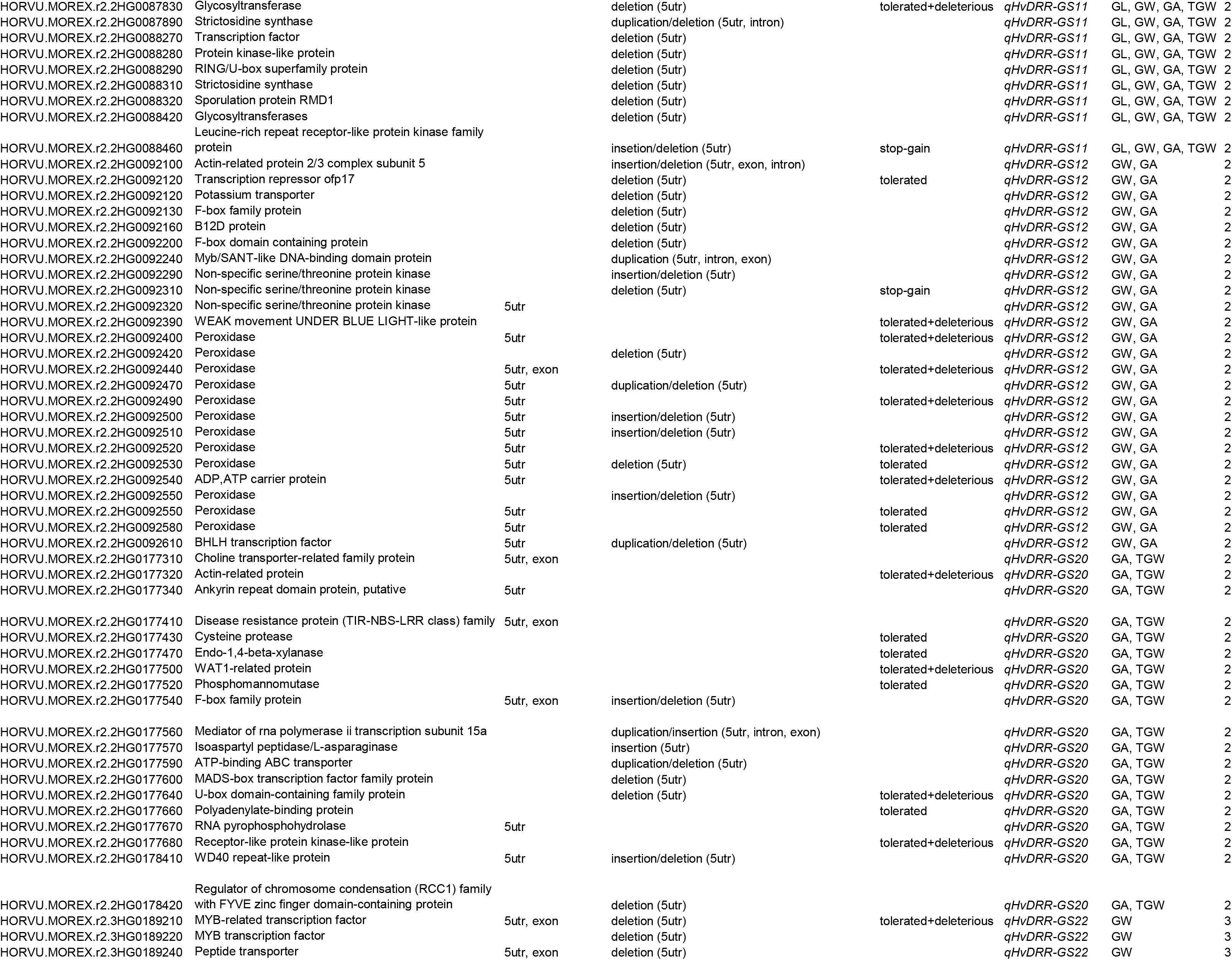
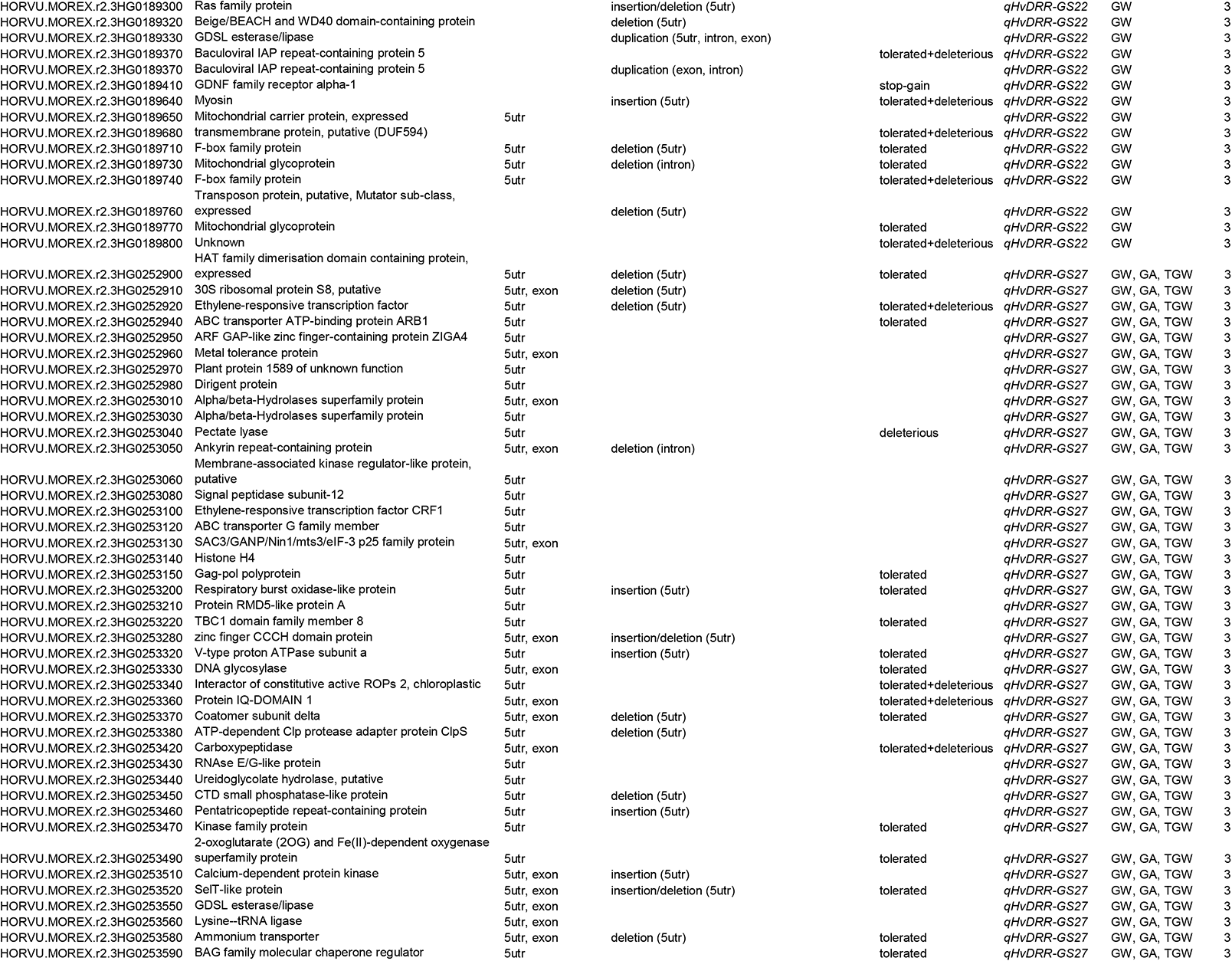
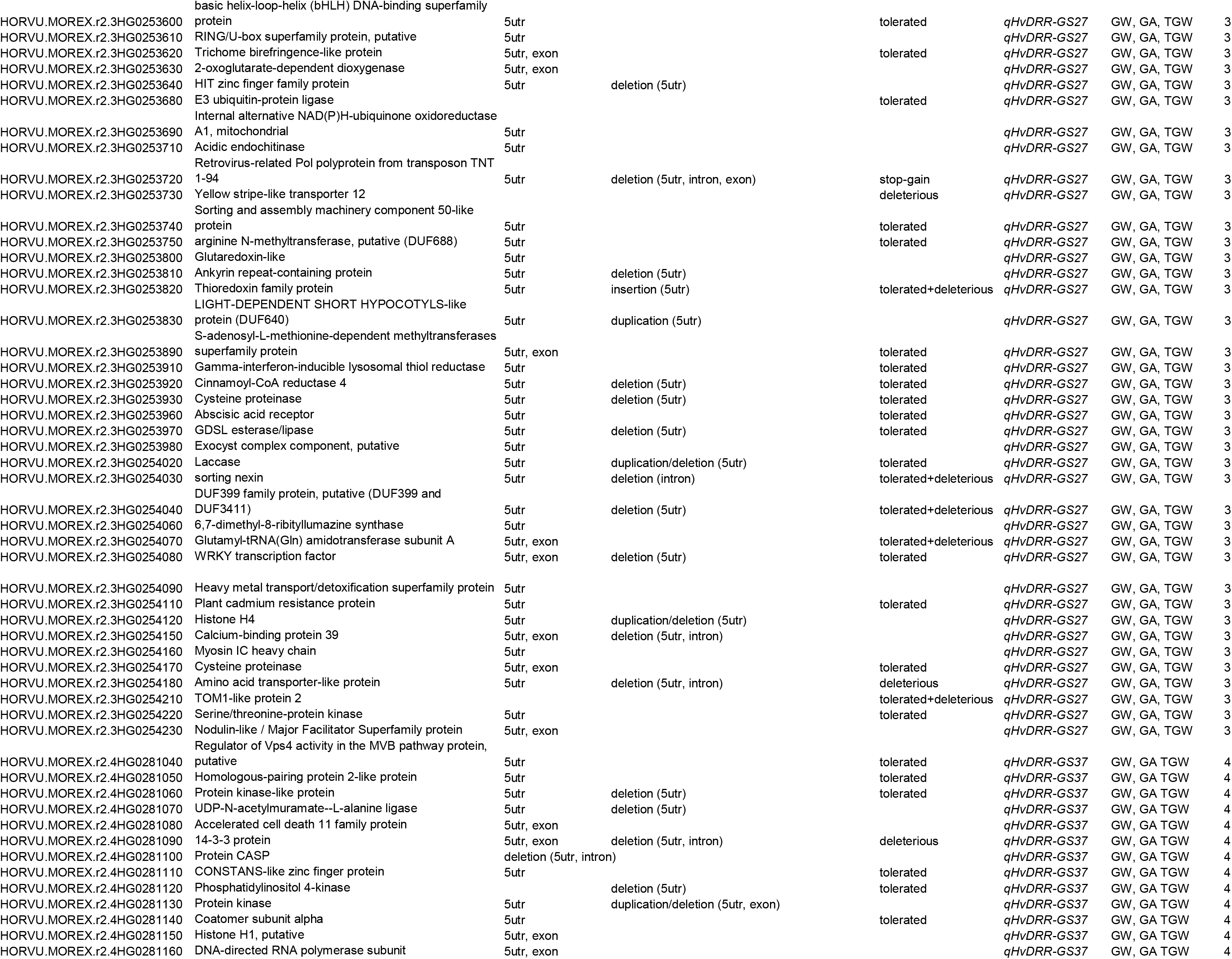
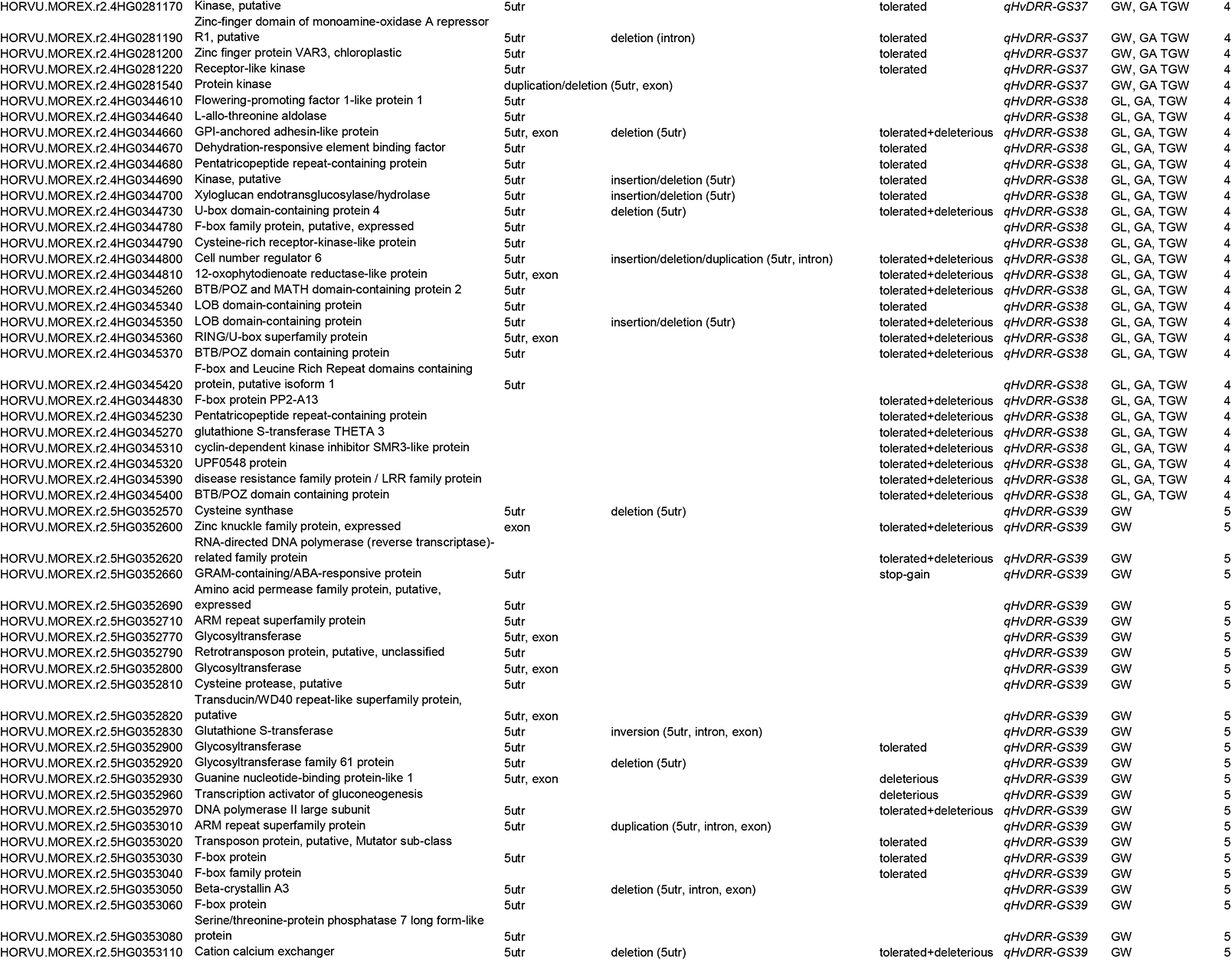
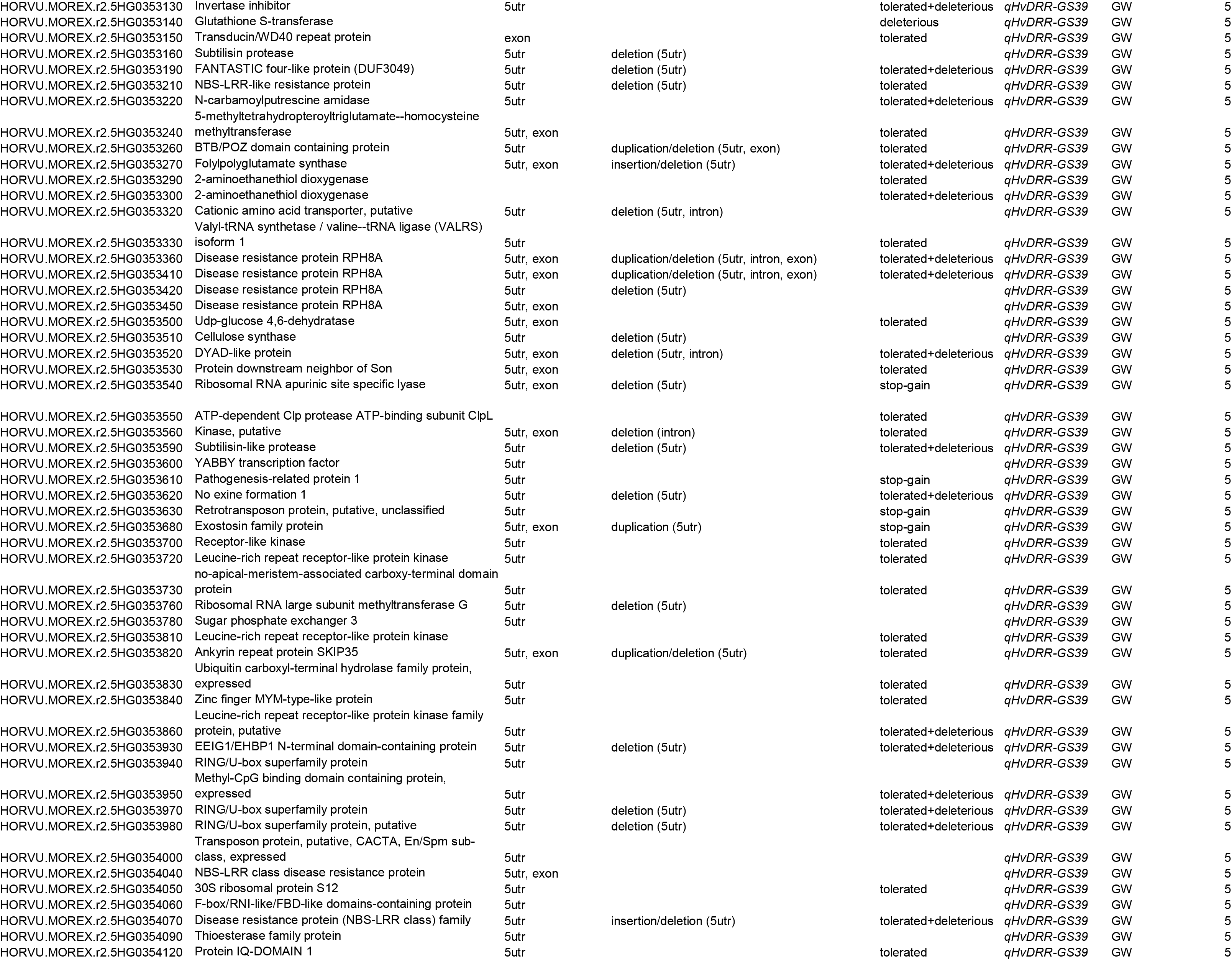
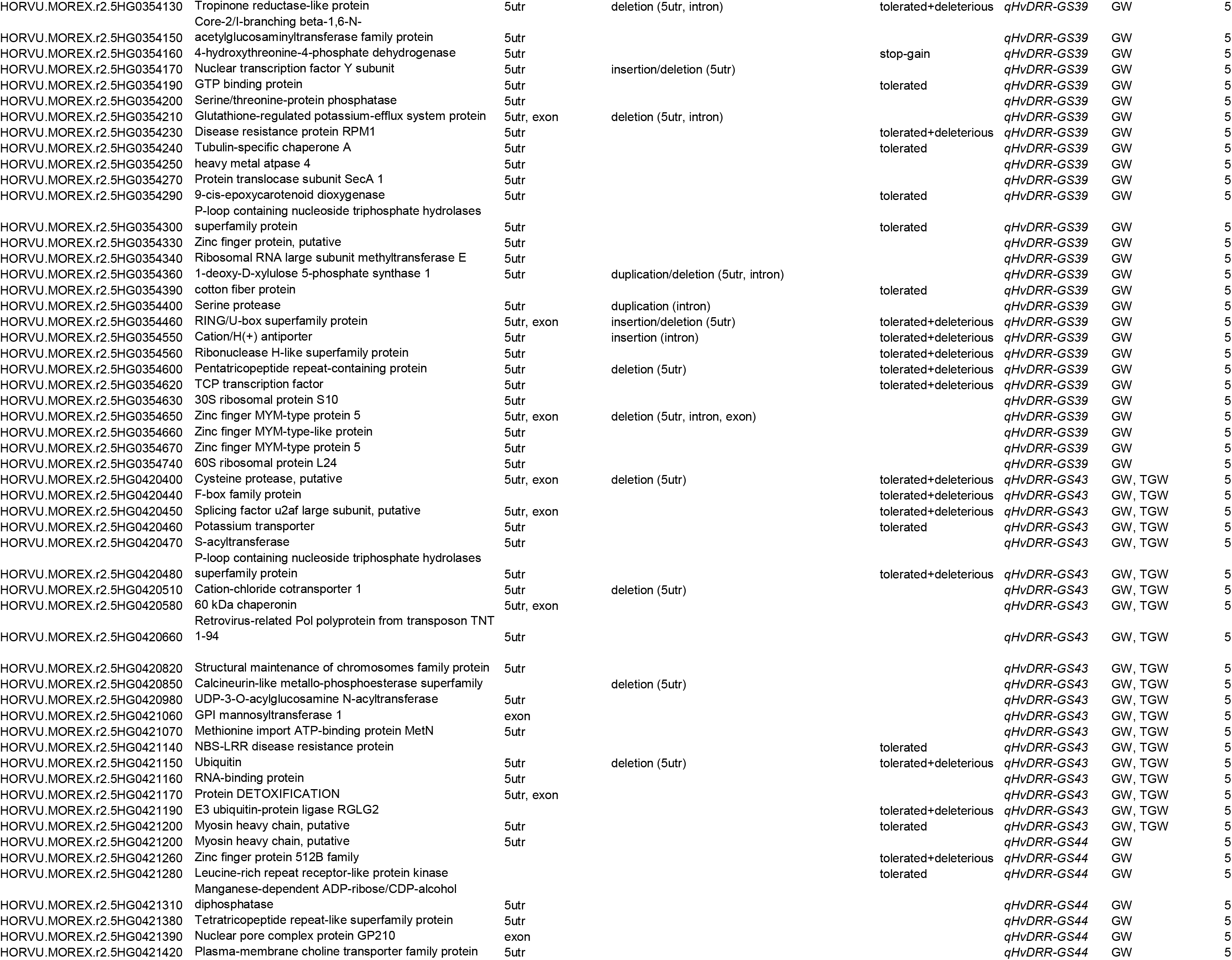
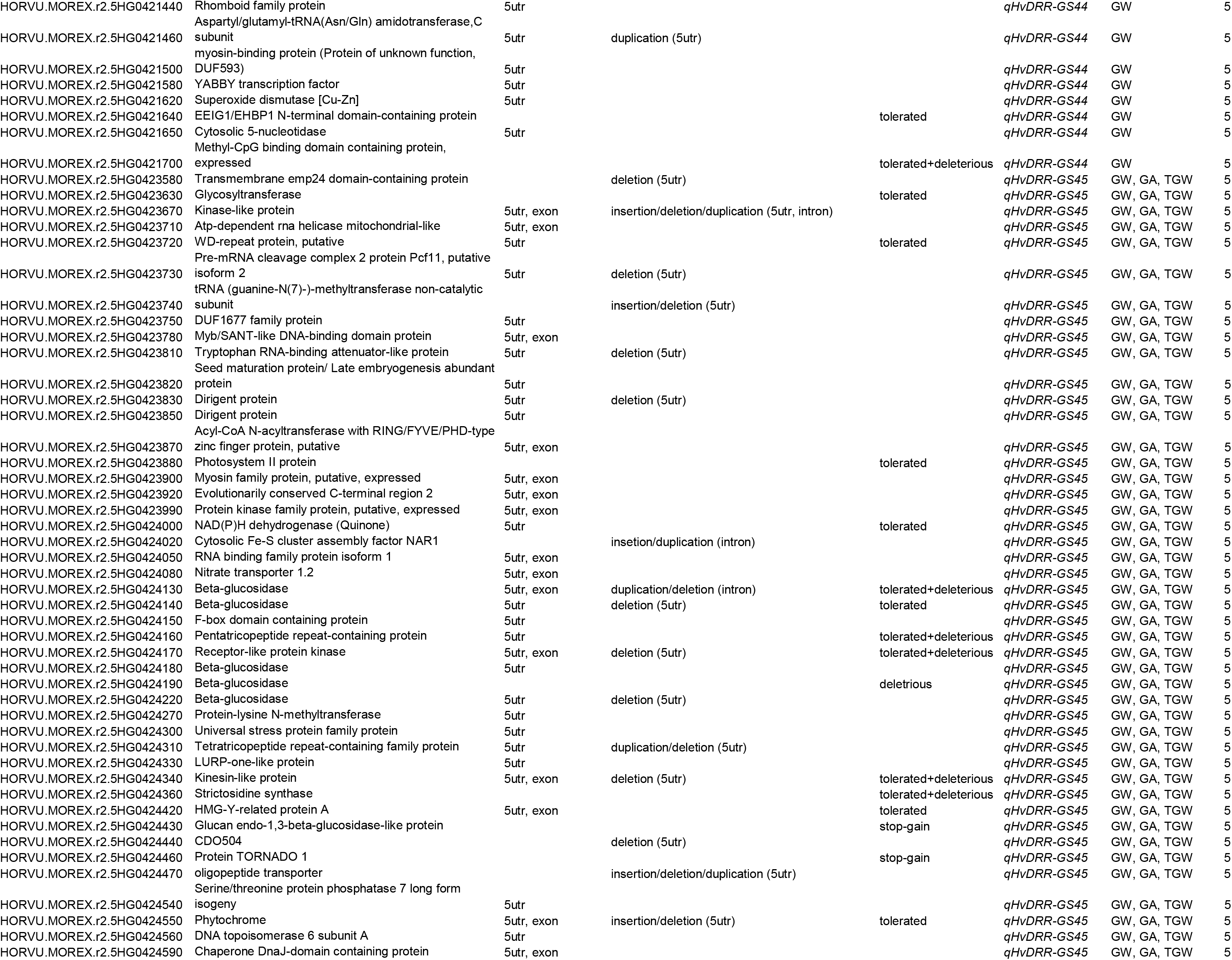
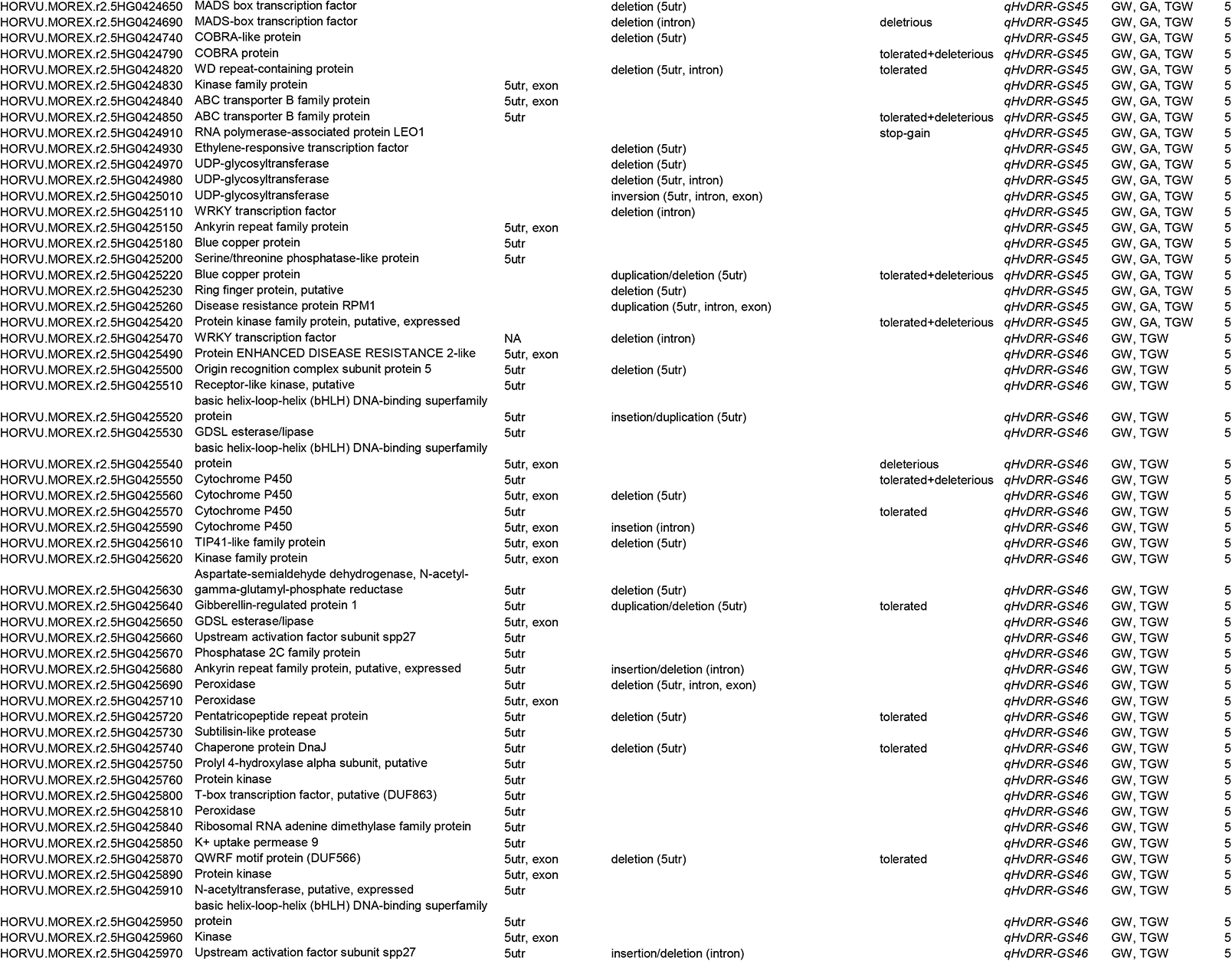
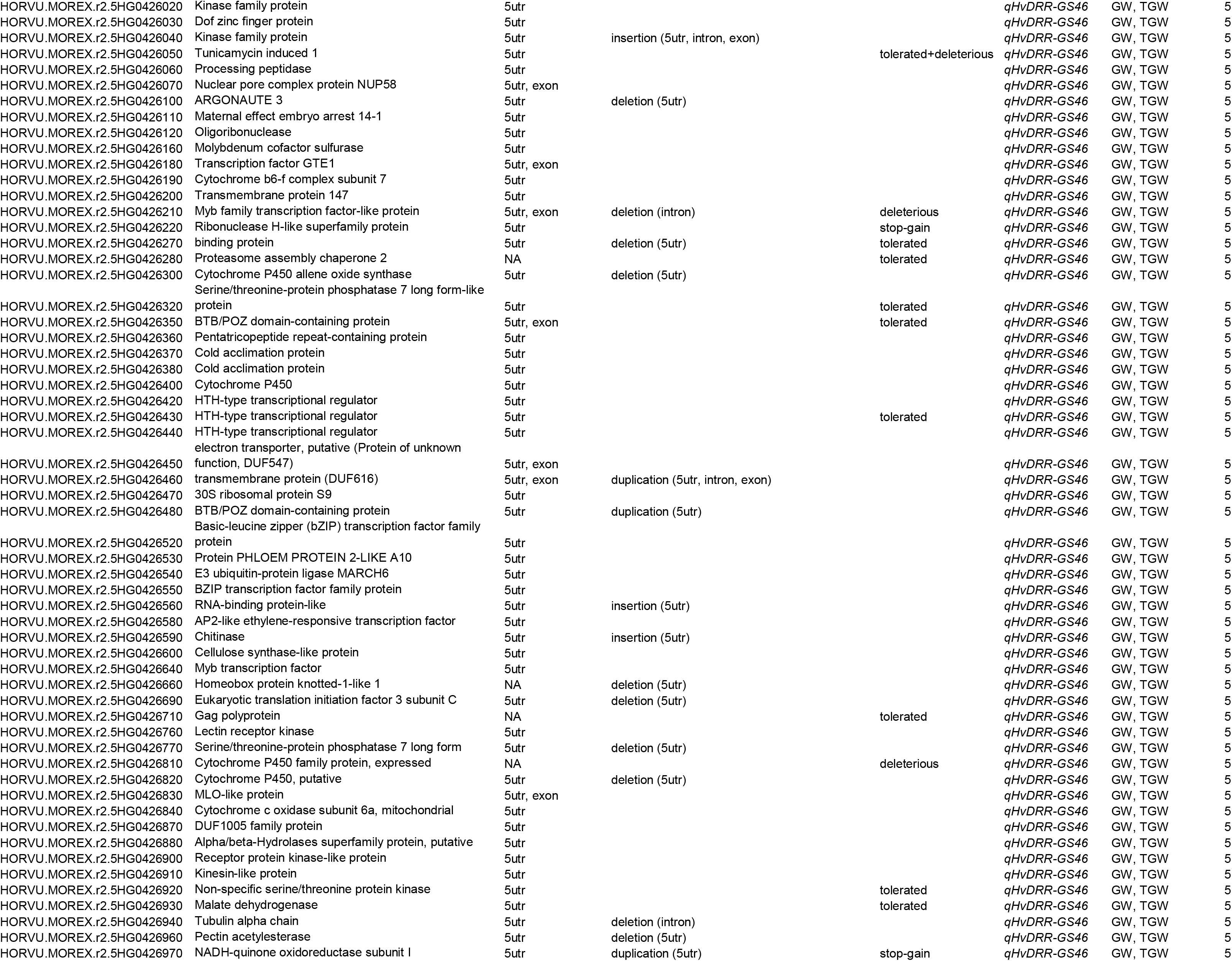
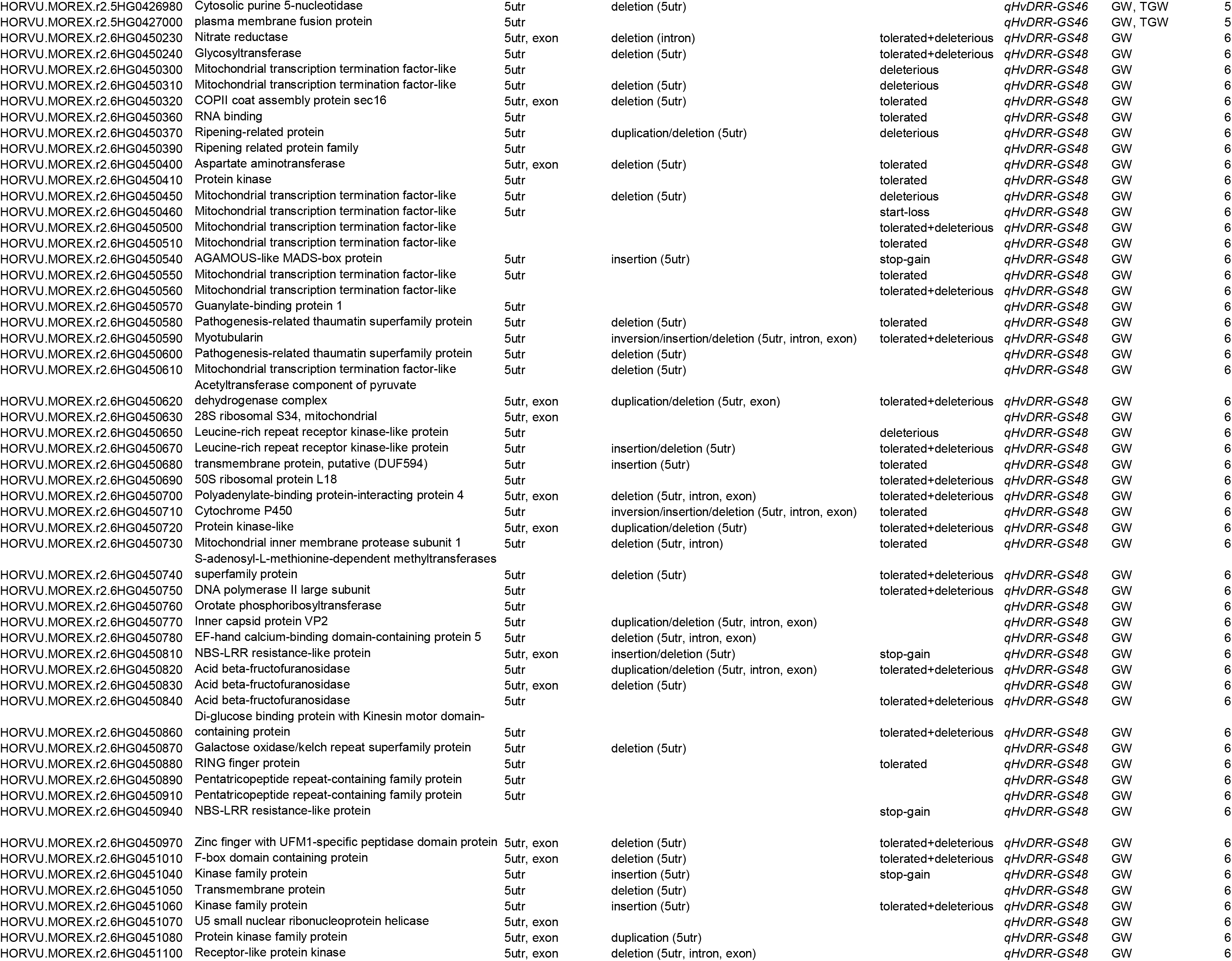
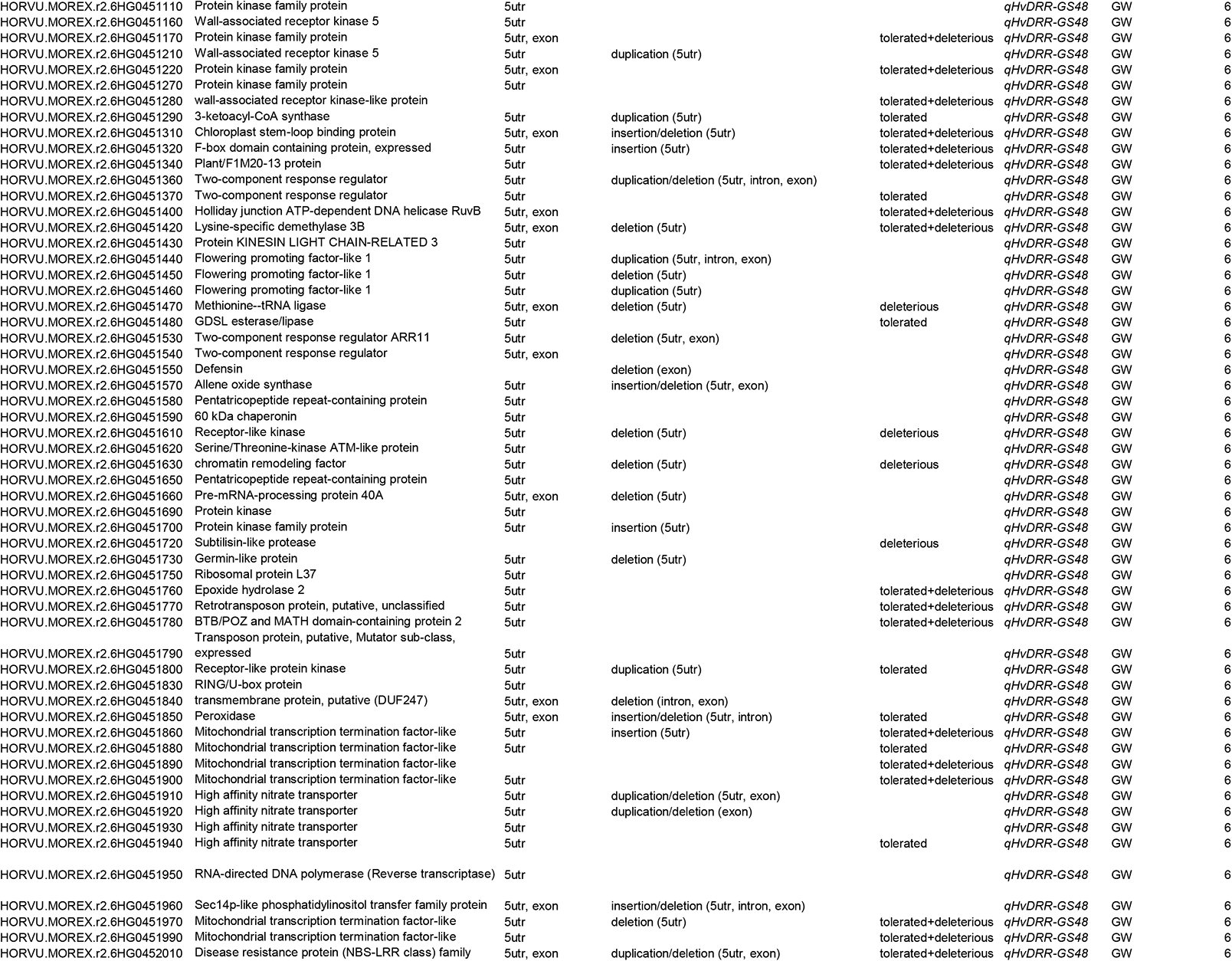
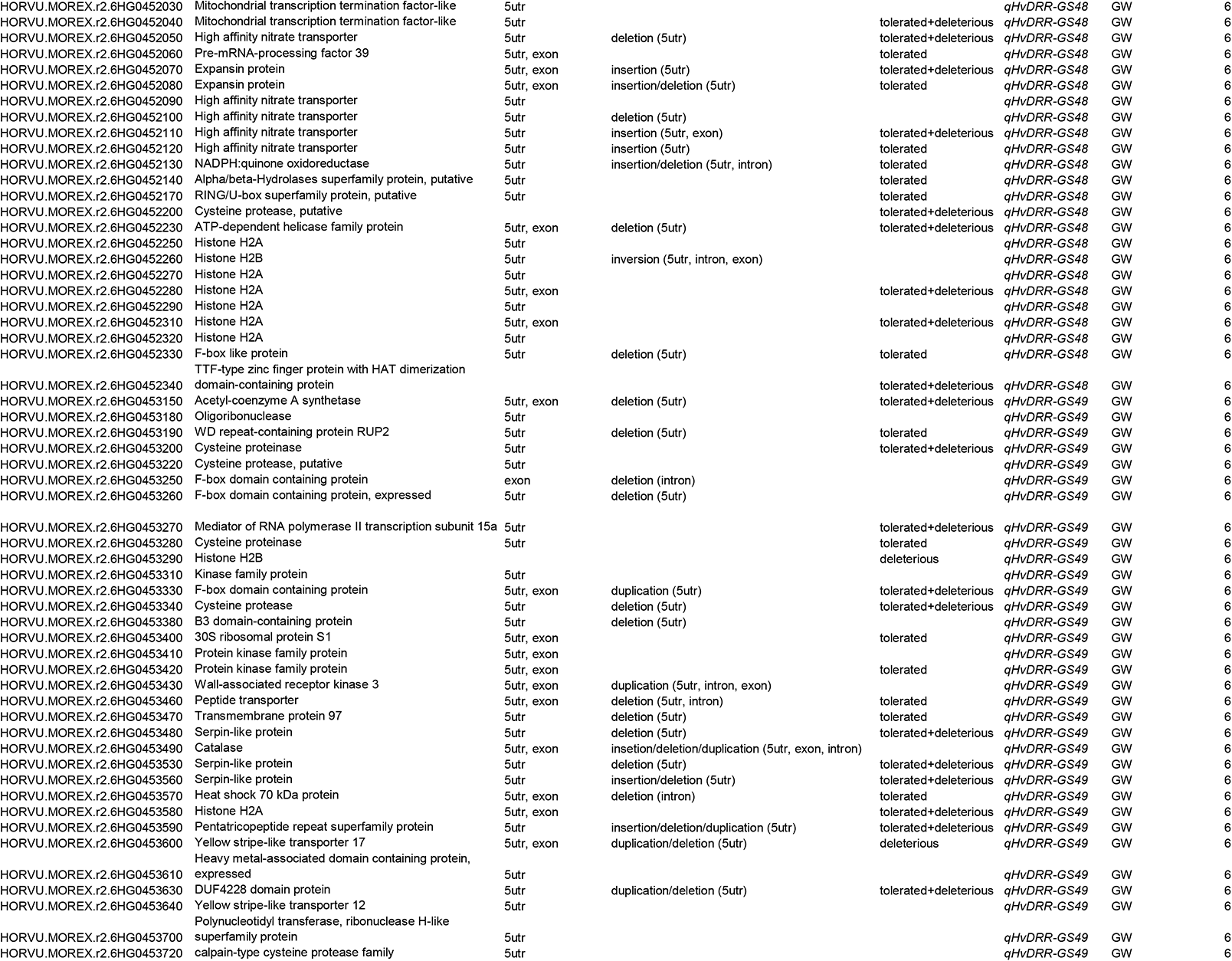
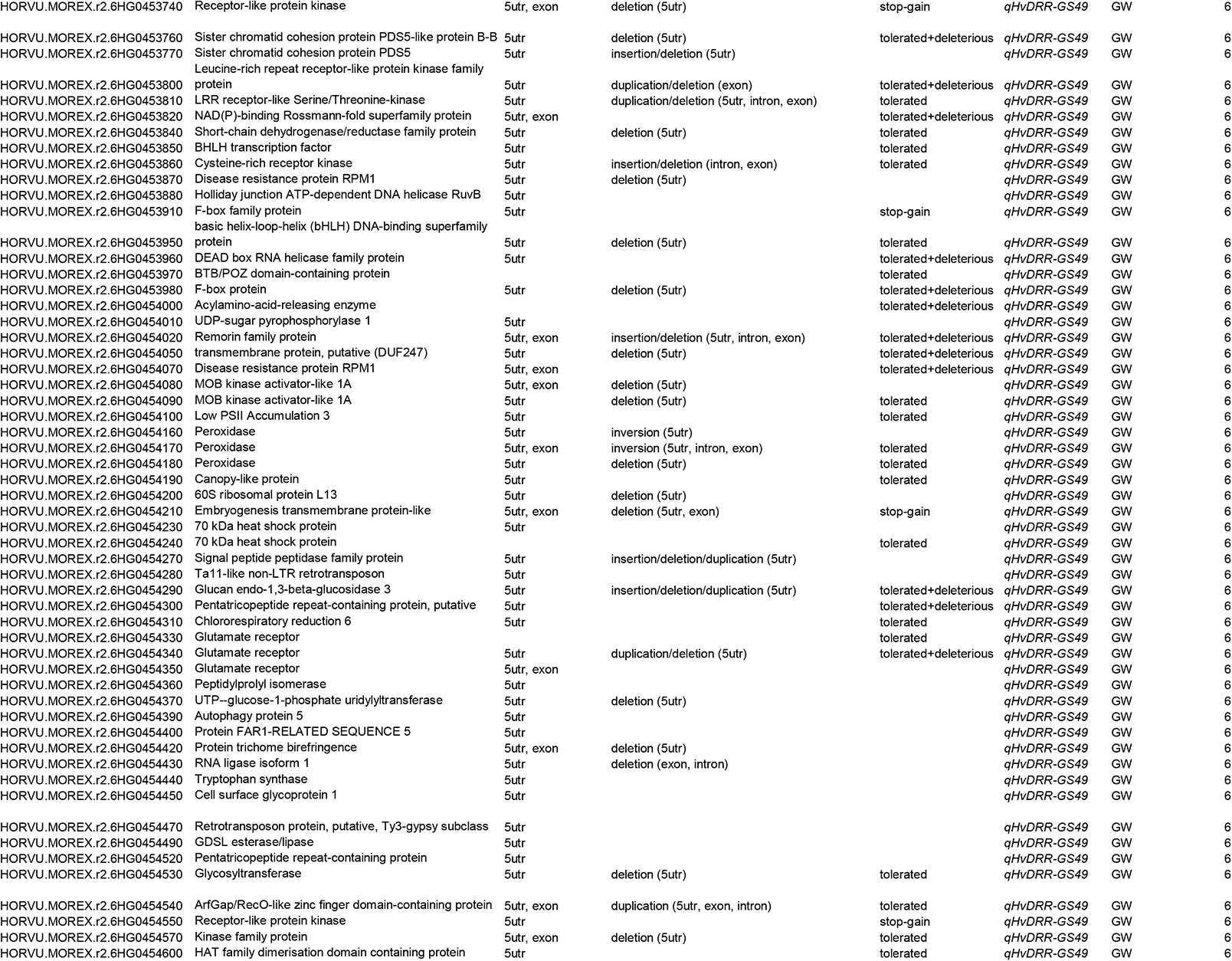
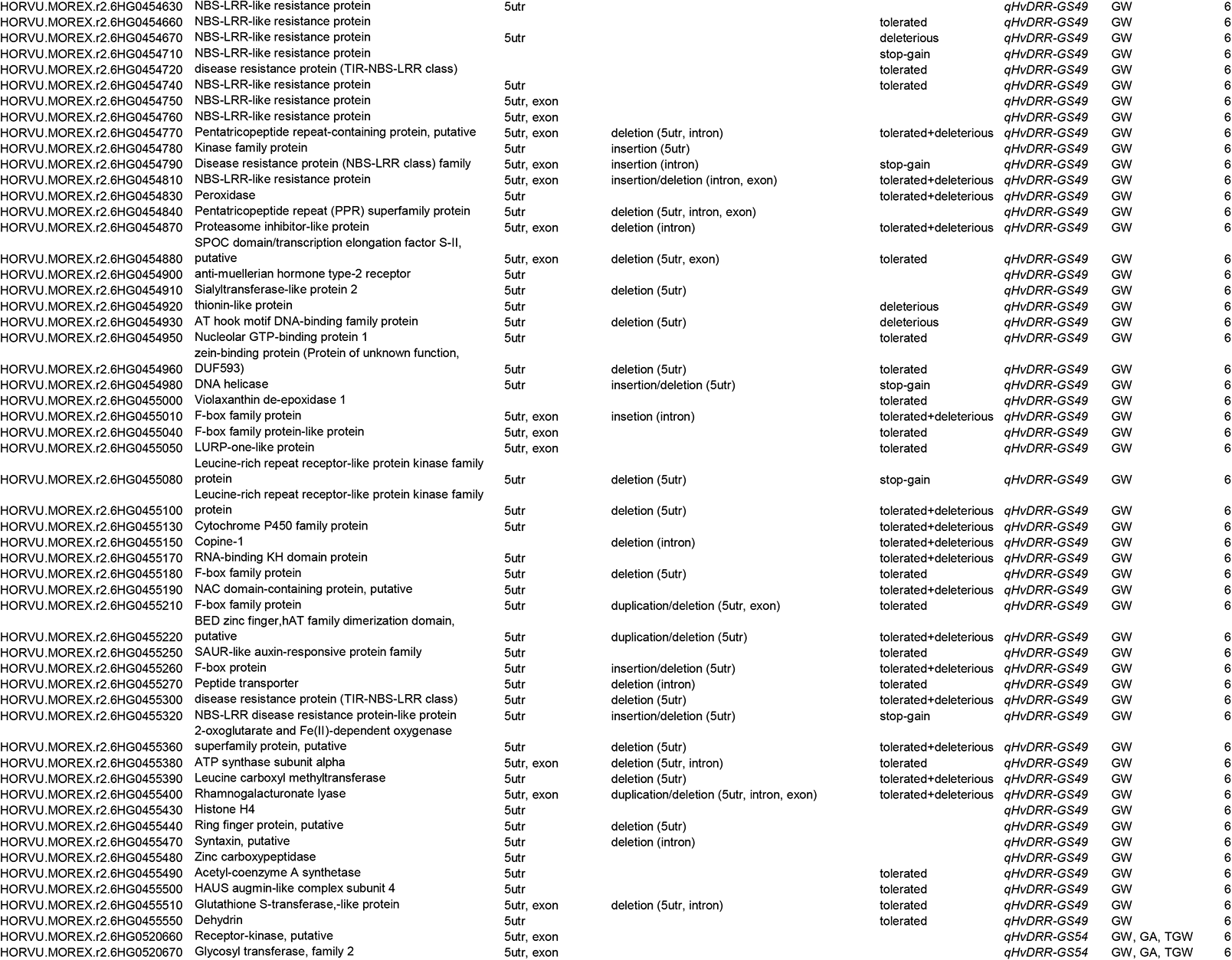
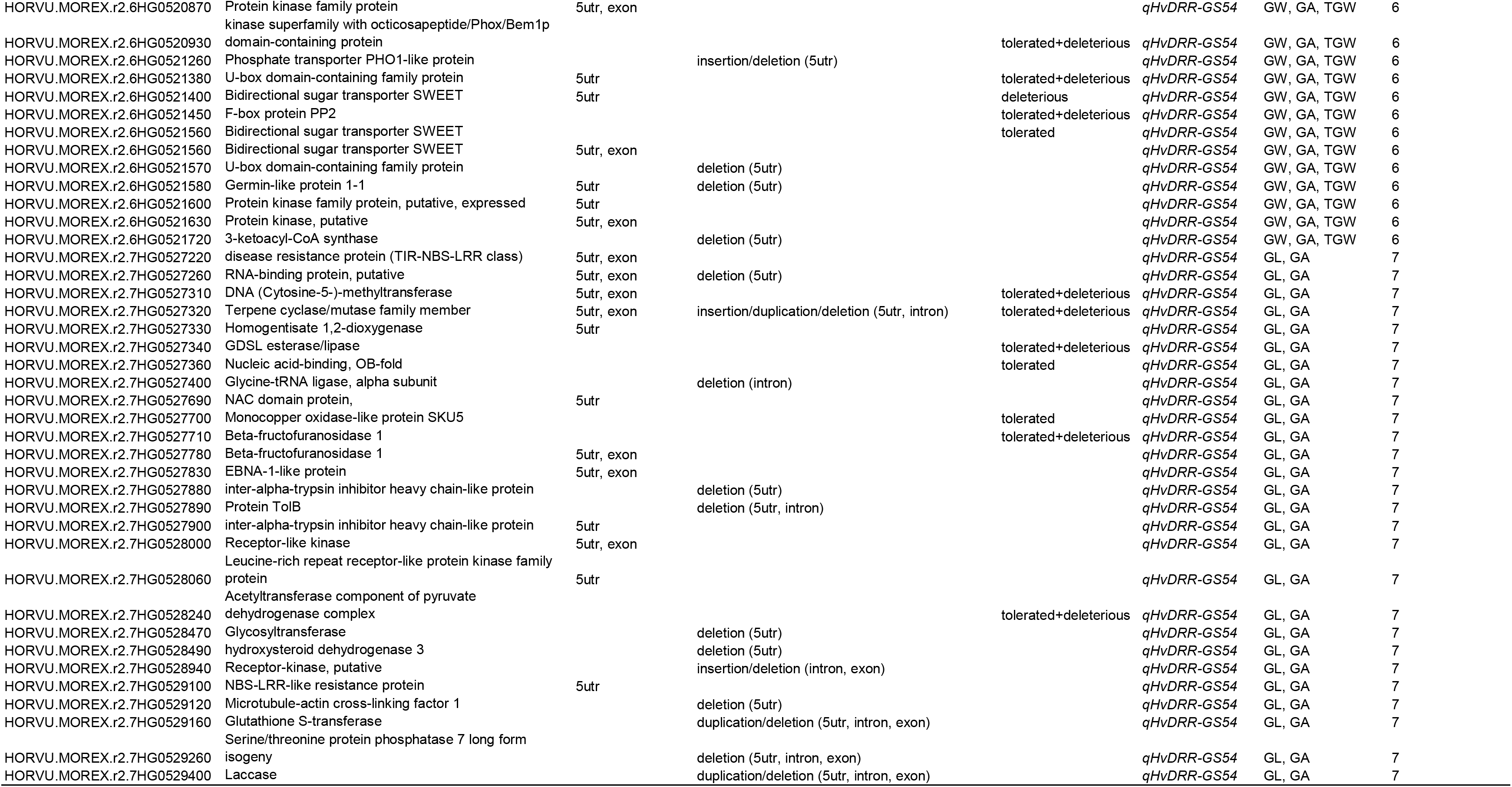
Candidate genes present in the consensus intervals. The detailed candidate gene mining was only performed for consensus interval with a physical size of less than 10 Mb and a percent of the explained phenotypic variance of more than 10 %. Sequence polymorphism data obtained from the whole-genome sequencing project of 23 parental inbreds was used to create indels (indel below 50 bp) and predicted structural variants (indels above 50 bp, duplication and inversions) in the coding and regulatory region and SNP annotation data (Weisweiler et al., 2022). The genes present in the consensus interval that revealed allelic differences between the contrasting groups of parental genotypes at a given QTL locus were retained as candidate genes. GL: grain length; GW: grain width; GA: grain area; TGW: thousand-grain weight

**Table S9:**
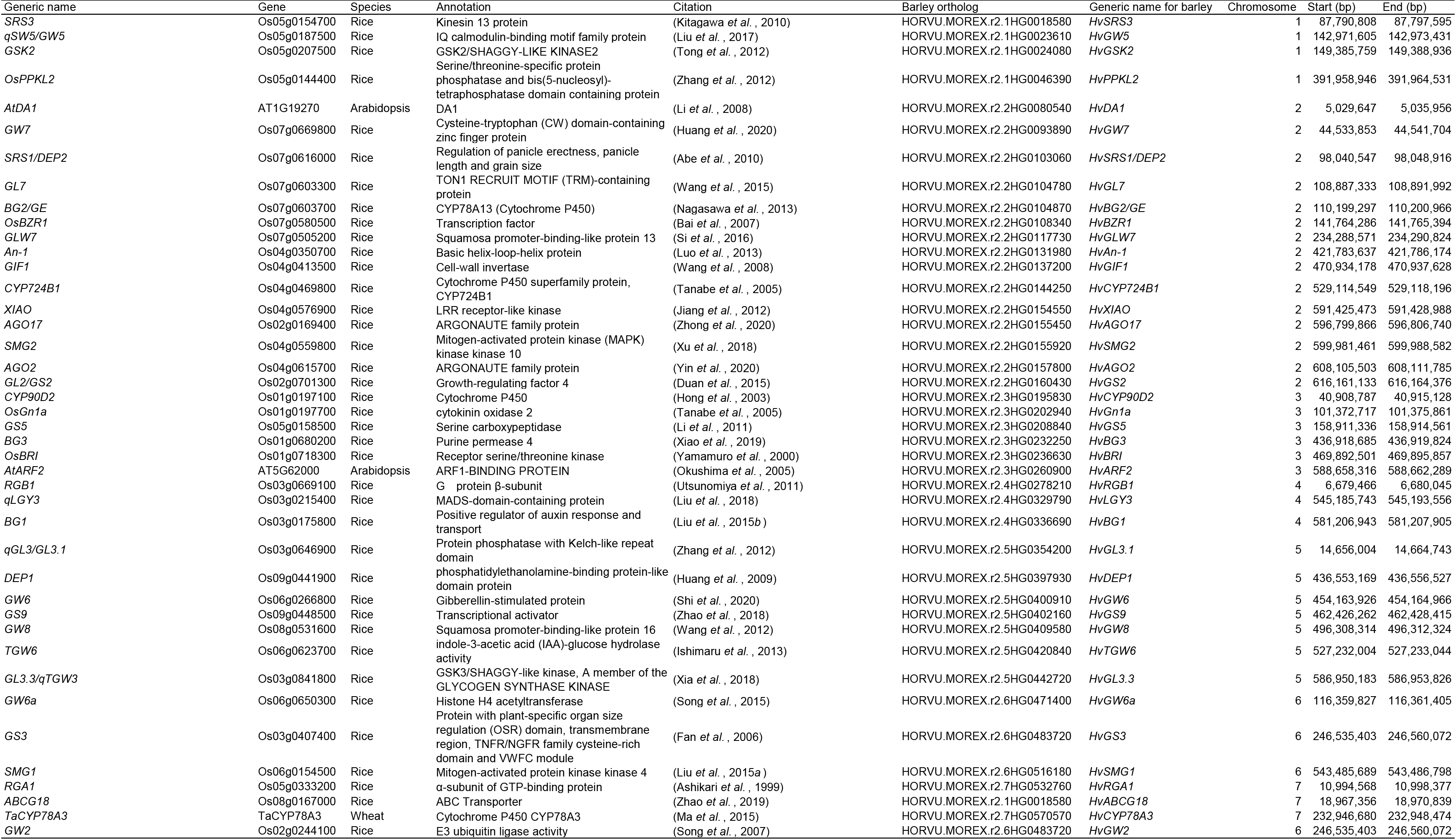
The barley orthologs of genes associated with seed size in other plant species.

**Fig. S1:**
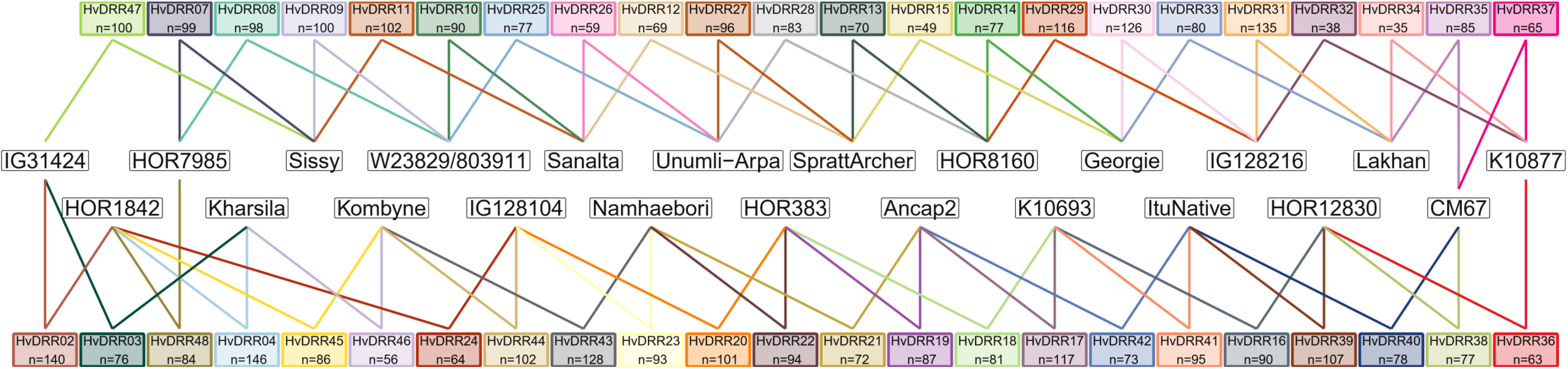
The double round-robin (DRR) crossing scheme used to establish the HvDRR population. Parental inbred lines for each individual HvDRR sub-population are connected by the lines and the ‘n’ indicates the number of recombinant inbred lines (Casale *et al.,* 2021).

**Fig. S2:**
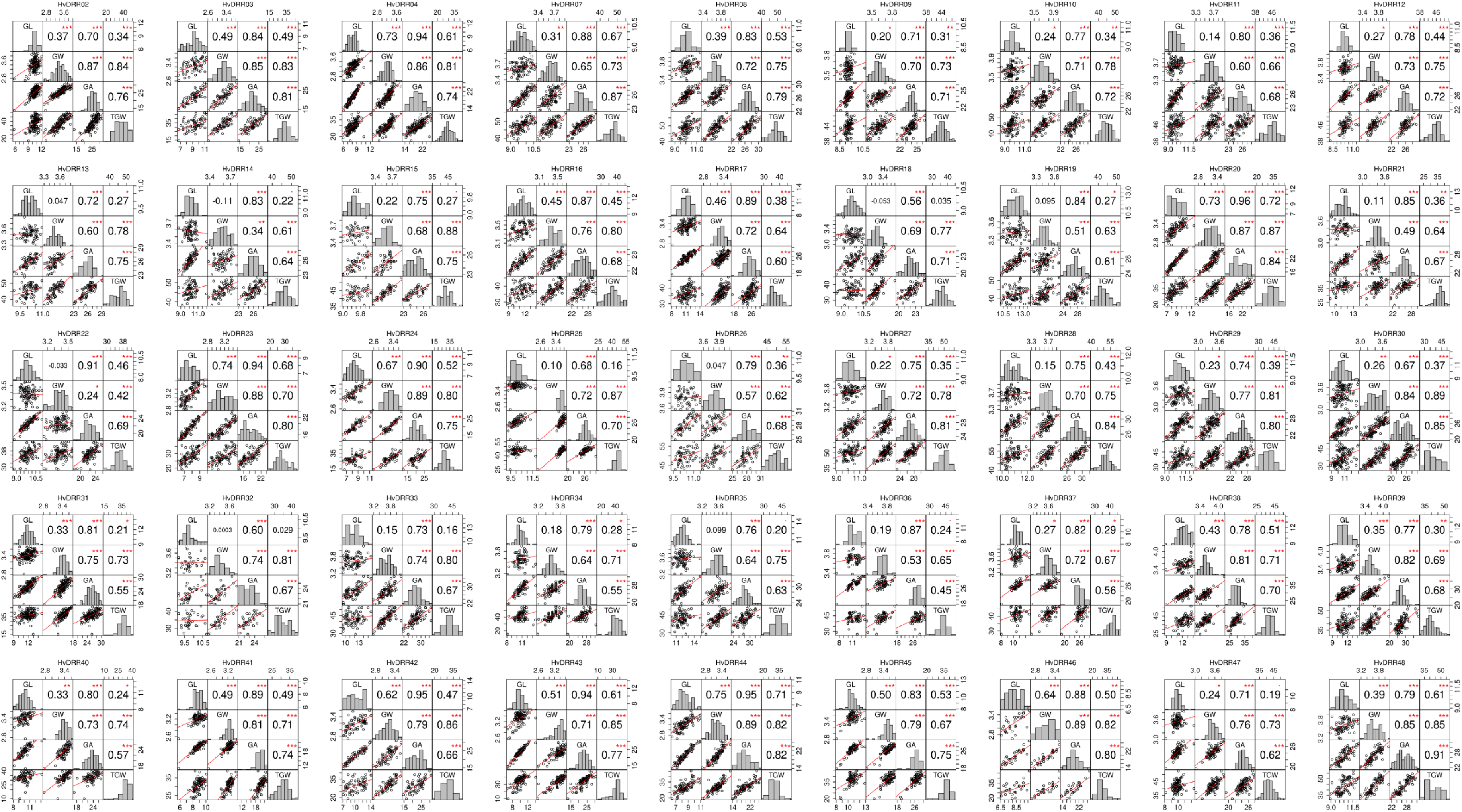
Pair-wise correlation analysis between the four evaluated traits namely grain length (GL) in mm, grain width (GW) in mm, grain area (GA) in mm^2^ and thousand grain weight (TGW) in g across 45 HvDRR sub-populations.

**Fig. S3:**
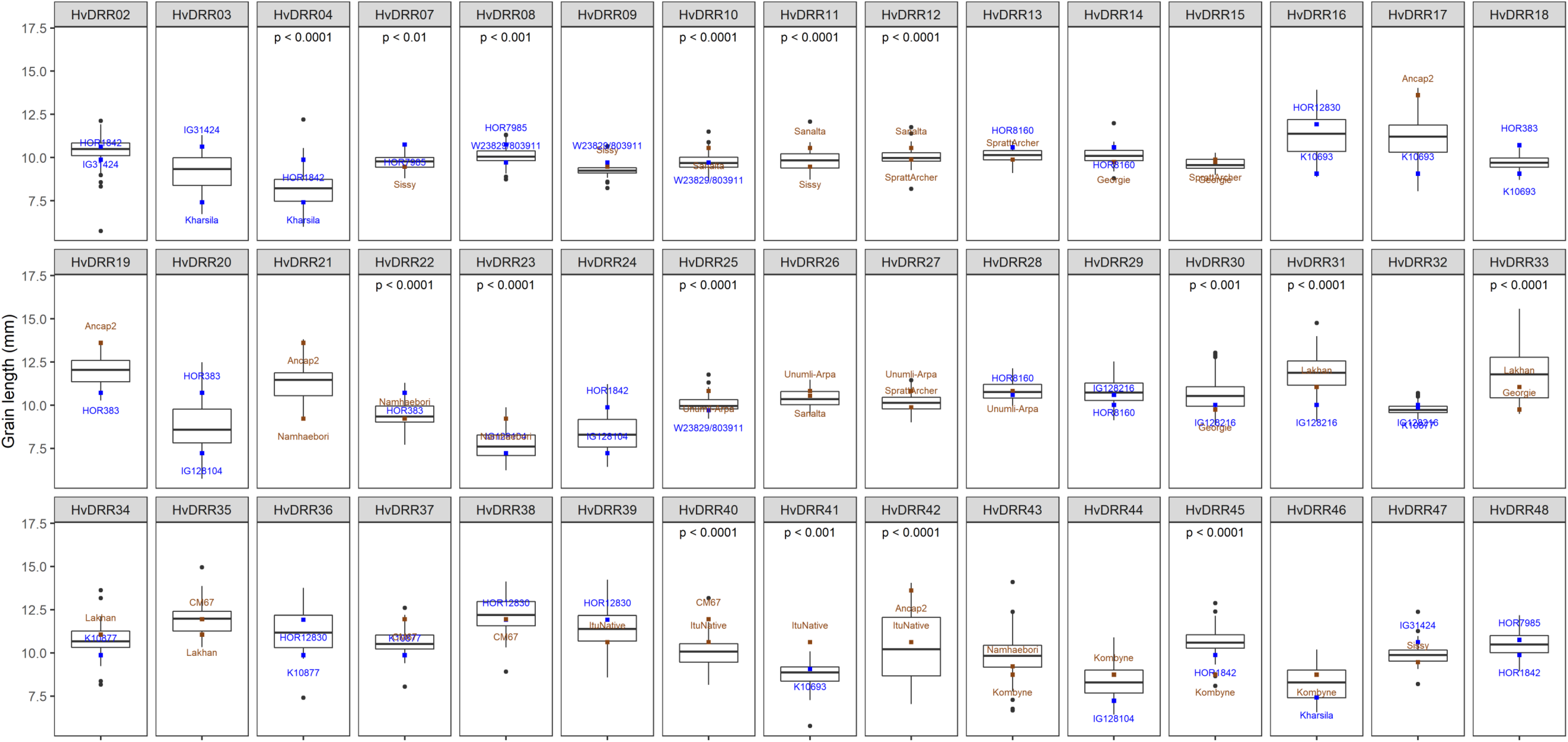
Boxplot of adjusted entry means of the recombinant inbred lines of the HvDRR populations for grain length in mm. The square dots indicate the adjusted entry mean of the parental inbreds of the respective recombinant inbred line populations. Brown and blue colors designate germplasm-type cultivars and landraces, respectively. The p-value above the boxplot indicates a significant mean differences between the segregating populations and the respective parental inbreds.

**Fig. S4:**
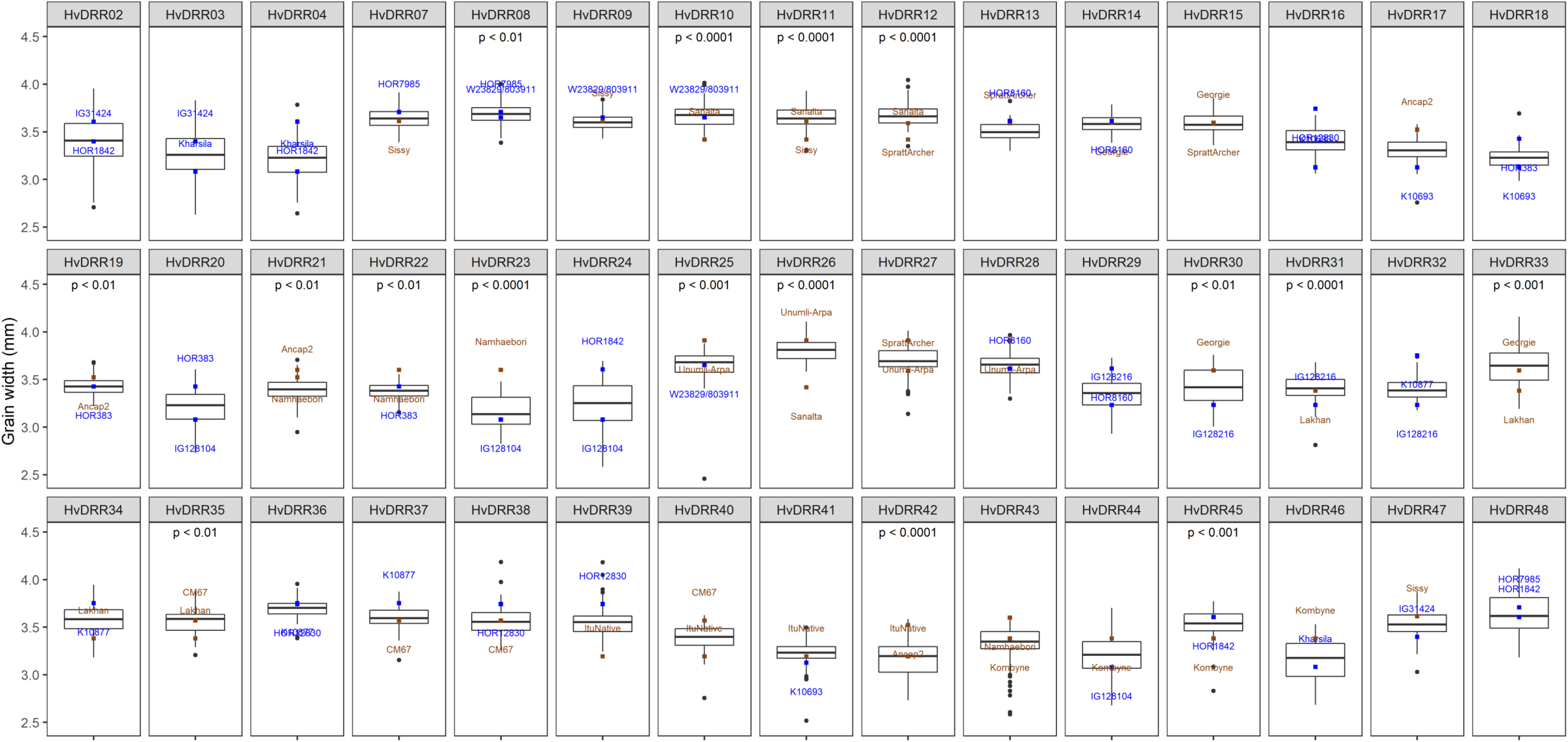
Boxplot of adjusted entry means of the recombinant inbred lines of the HvDRR populations for grain width in mm. The square dots indicate the adjusted entry mean of the parental inbreds of the respective recombinant inbred line populations. Brown and blue colors designate germplasm-type cultivars and landraces, respectively. The p-value above the boxplot indicates a significant mean differences between the segregating populations and the respective parental inbreds.

**Fig. S5:**
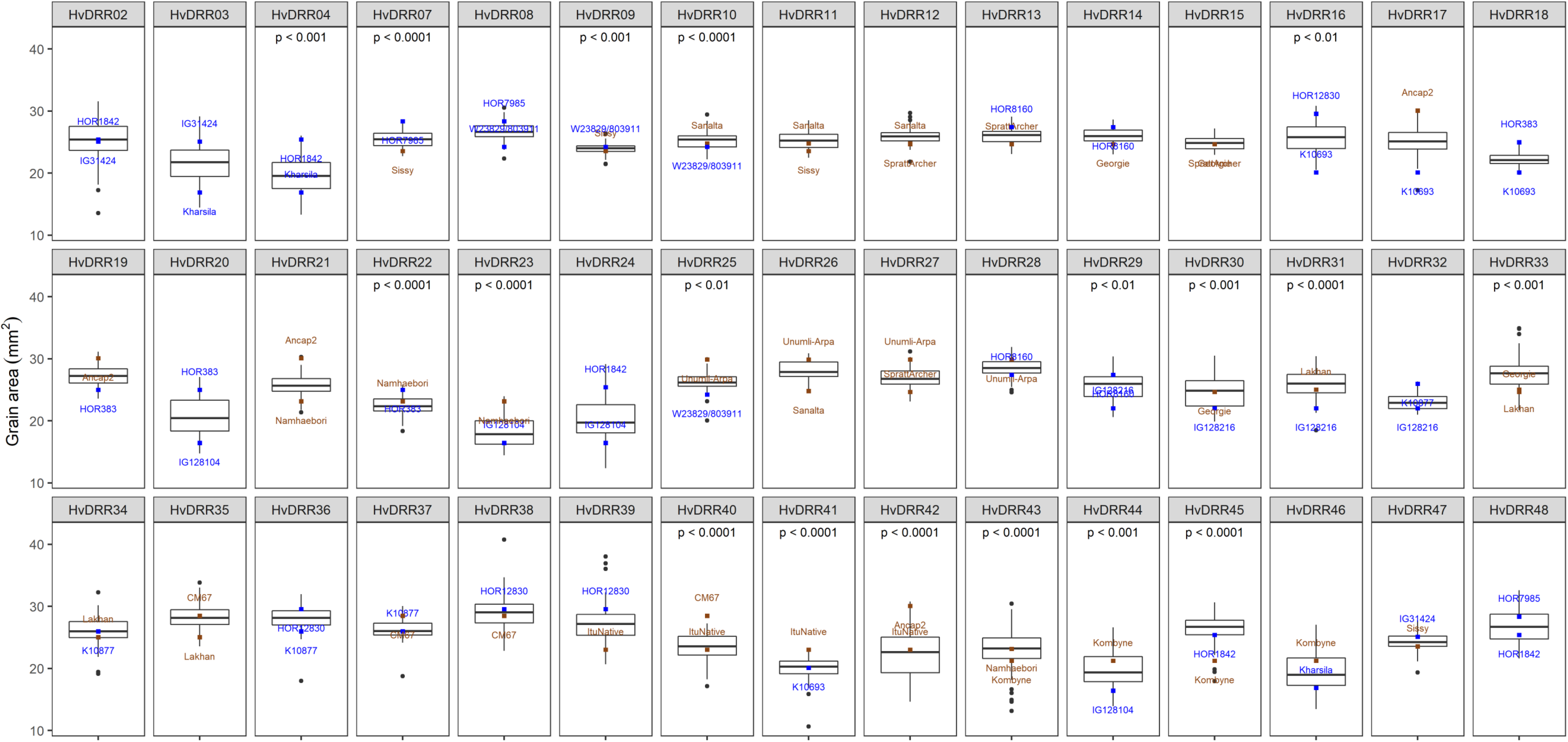
Boxplot of adjusted entry means of the recombinant inbred lines of the HvDRR populations for grain area in mm^2^. The square dots indicate the adjusted entry mean of the parental inbreds of the respective recombinant inbred line populations. Brown and blue colors designate germplasm-type cultivars and landraces, respectively. The p-value above the boxplot indicates a significant mean differences between the segregating populations and the respective parental inbreds.

**Fig. S6:**
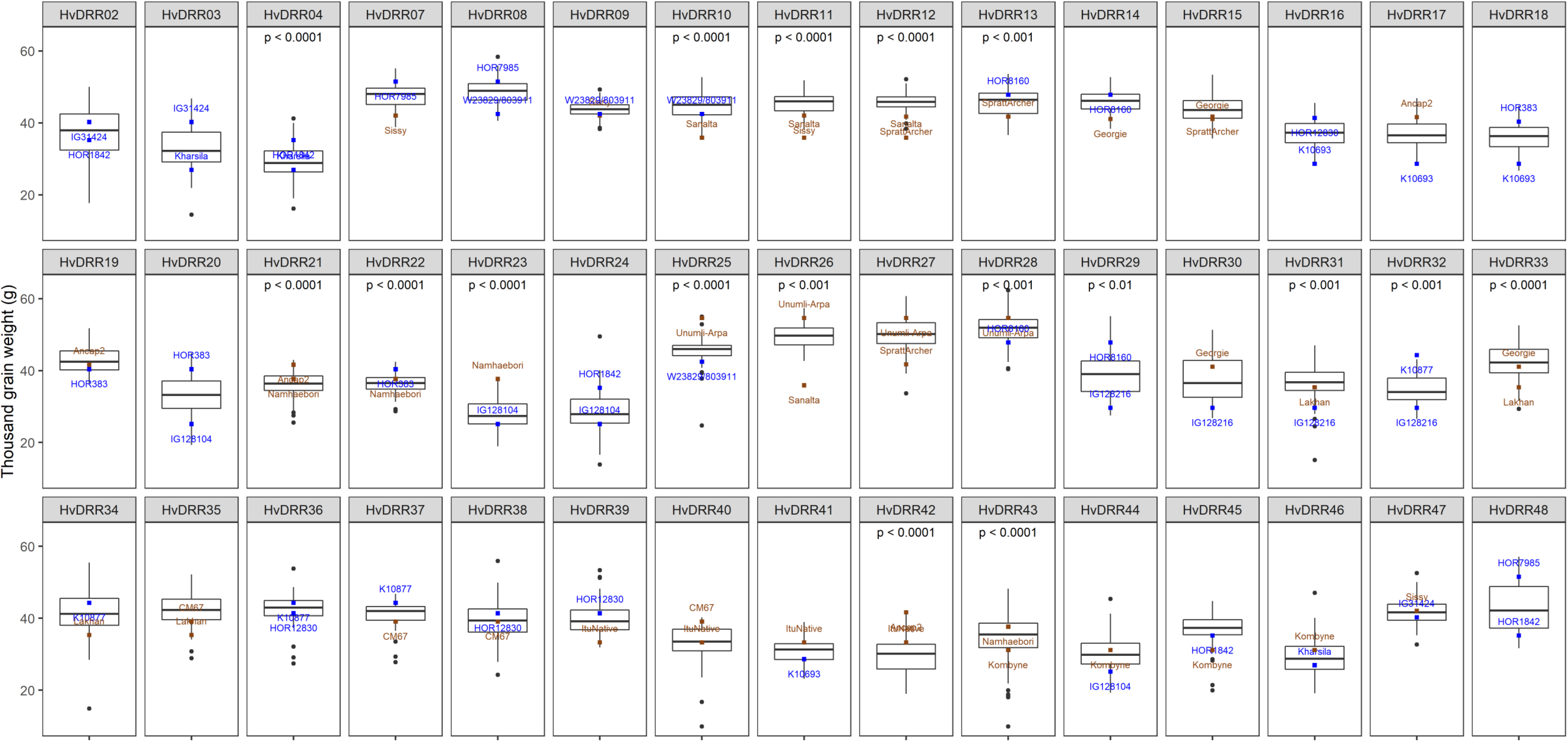
Boxplot of adjusted entry means of the recombinant inbred lines of the HvDRR populations for thousand-grain weight in grams. The square dots indicate the adjusted entry mean of the parental inbreds of the respective recombinant inbred line populations. Brown and blue colors designate germplasm-type cultivars and landraces, respectively. The p-value above the boxplot indicates a significant mean differences between the segregating populations and the respective parental inbreds.

**Fig. S7:**
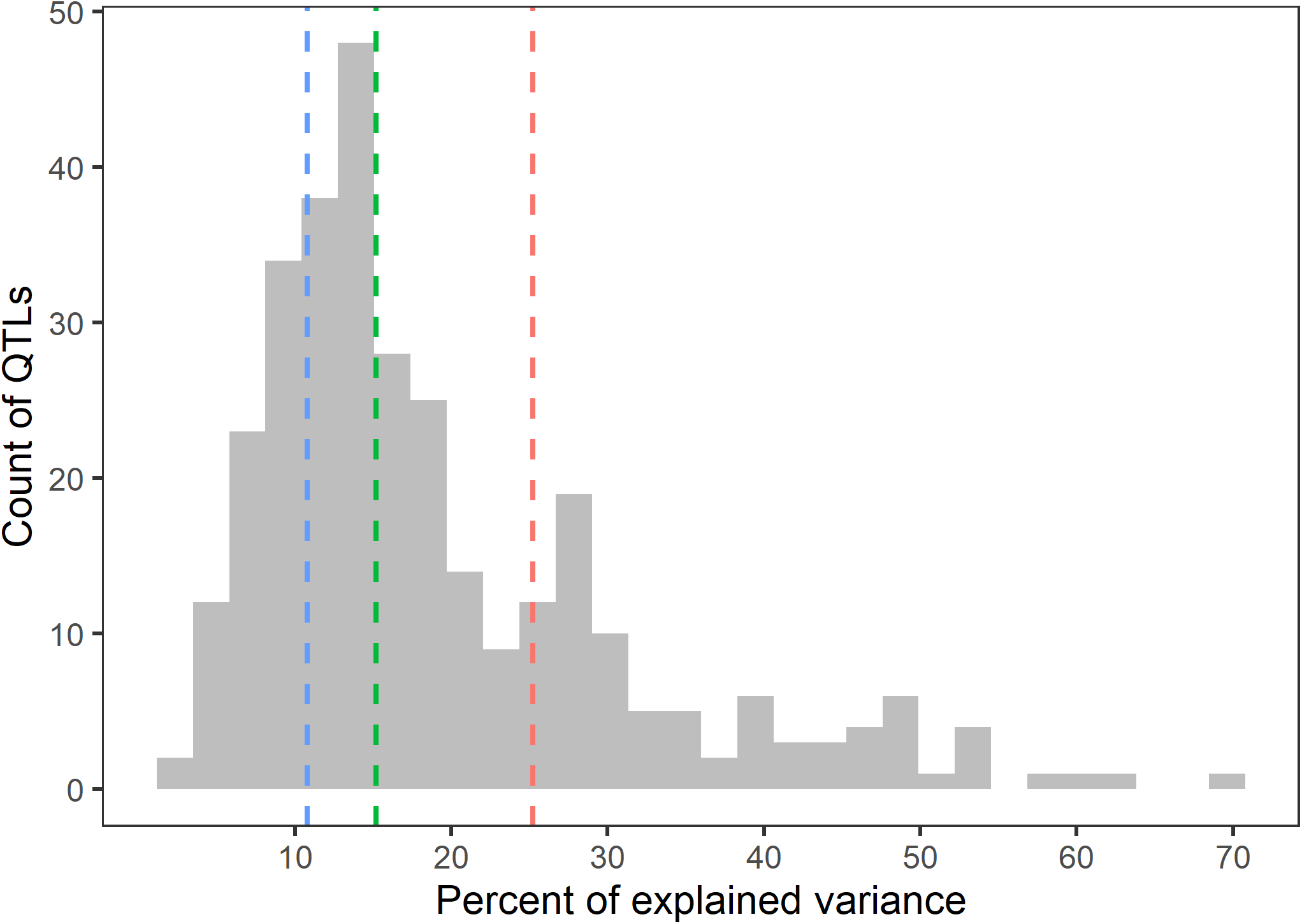
Distribution of the percentage of variance explained by quantitative trait loci detected in single population analyses for grain size and thousand-grain weight in 45 HvDRR sub-populations. The blue, green, and red dotted lines indicate the first quartile, median and third quartile of the distribution.

**Fig. S8:**
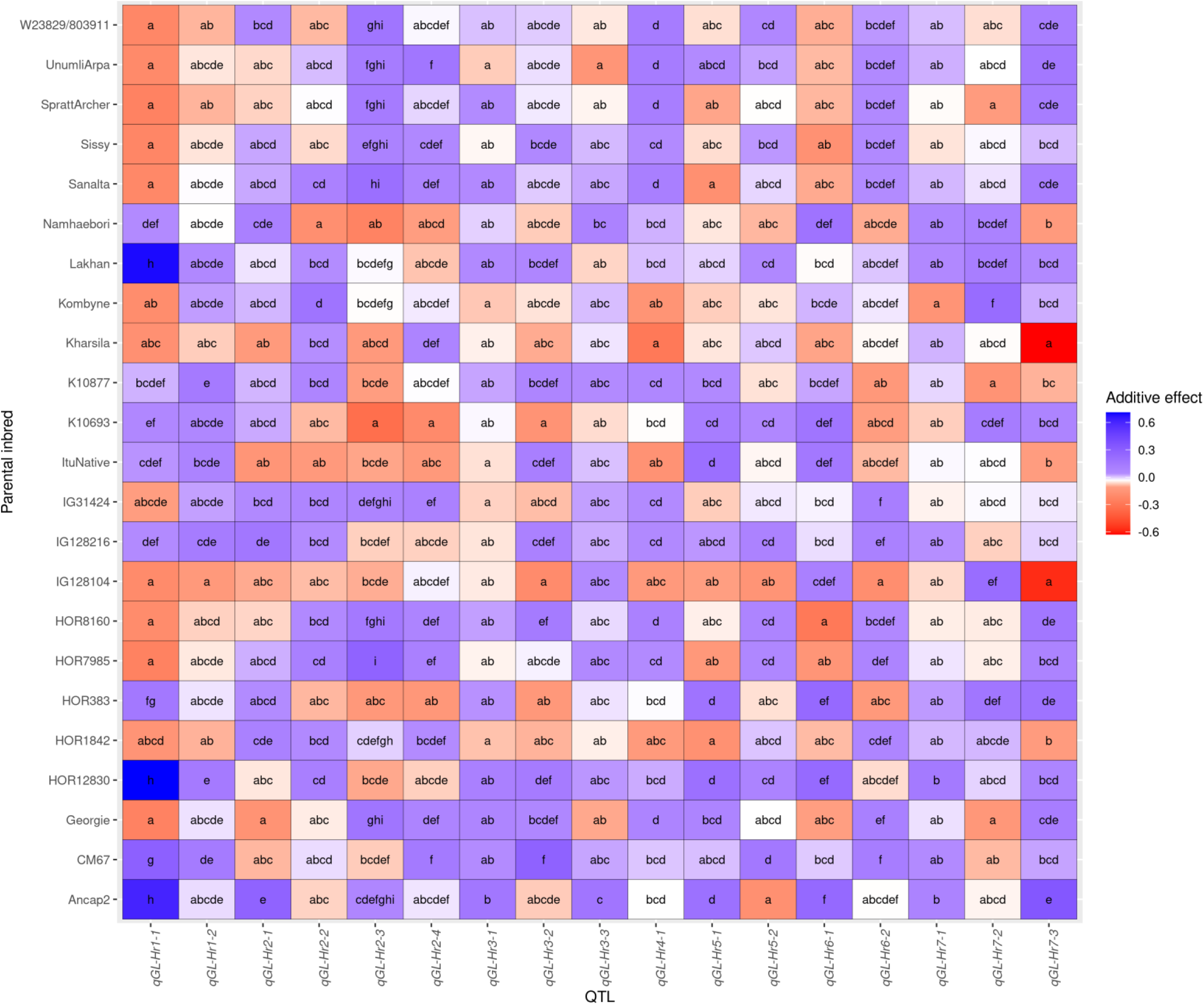
Multiple comparisons of the standardized allele effect for grain length quantitative trait loci (QTLs) detected in a multi-parent population analysis using a parental model. The standardized allele effect for an inbred is the difference between the mean of the estimated additive effect for 23 inbreds and the estimated allele effect of the corresponding inbred. The color code indicates the magnitude of the standardized allele effect. Indexed letters indicate the significant difference (p ≤ 0.05) between the genotypes not sharing the same letter by Tukey’s HSD test.

**Fig. S9:**
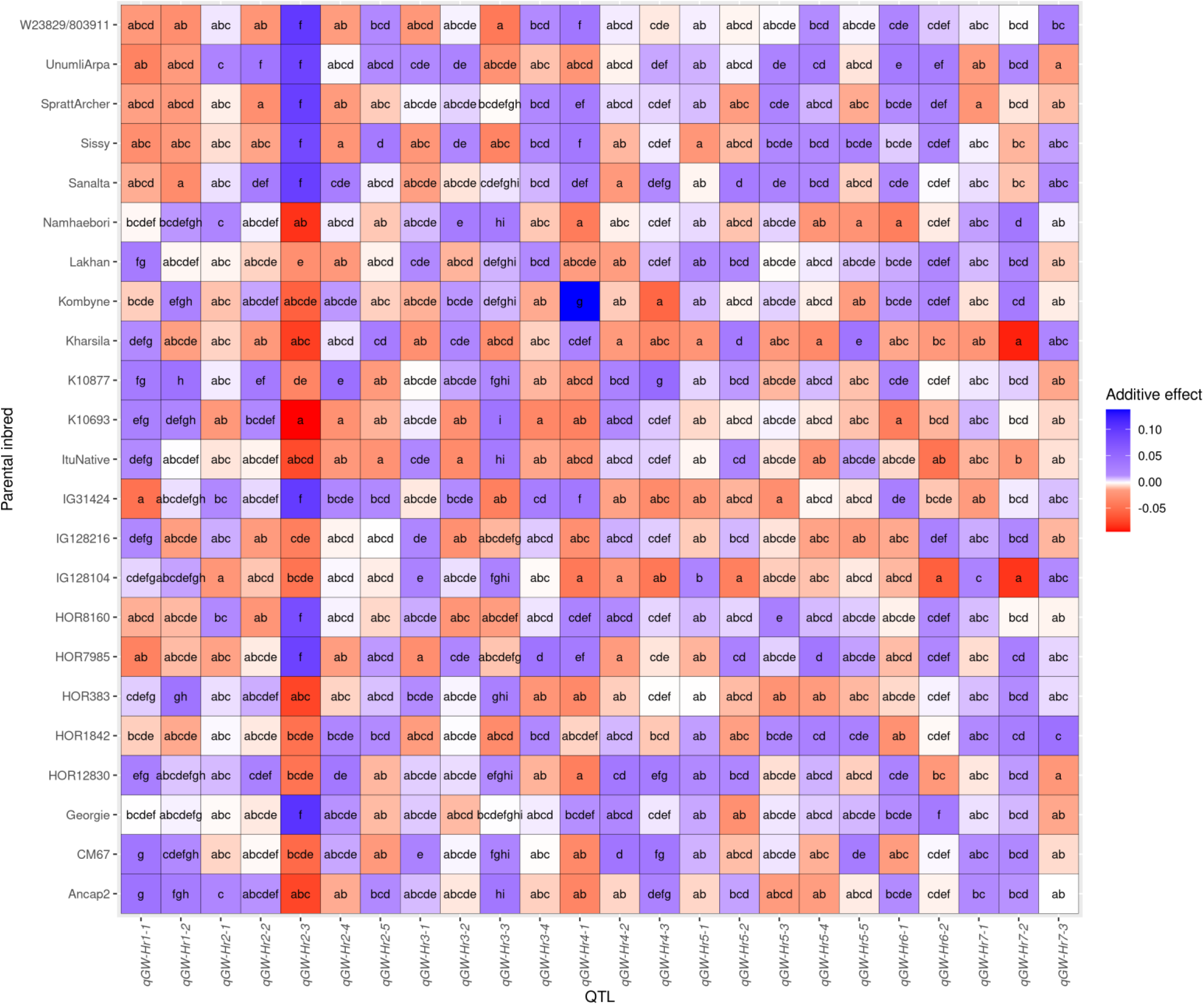
Multiple comparisons of the standardized allele effect for grain width quantitative trait loci (QTLs) detected in a multi-parent population analysis using a parental model. The standardized allele effect for an inbred is the difference between the mean of the estimated allele effect for 23 inbreds and the estimated additive effect of the corresponding inbred. The color code indicates the magnitude of the standardized allele effect. Indexed letters indicate the significant difference (p ≤ 0.05) between the genotypes not sharing the same letter by Tukey’s HSD test.

**Fig. S10:**
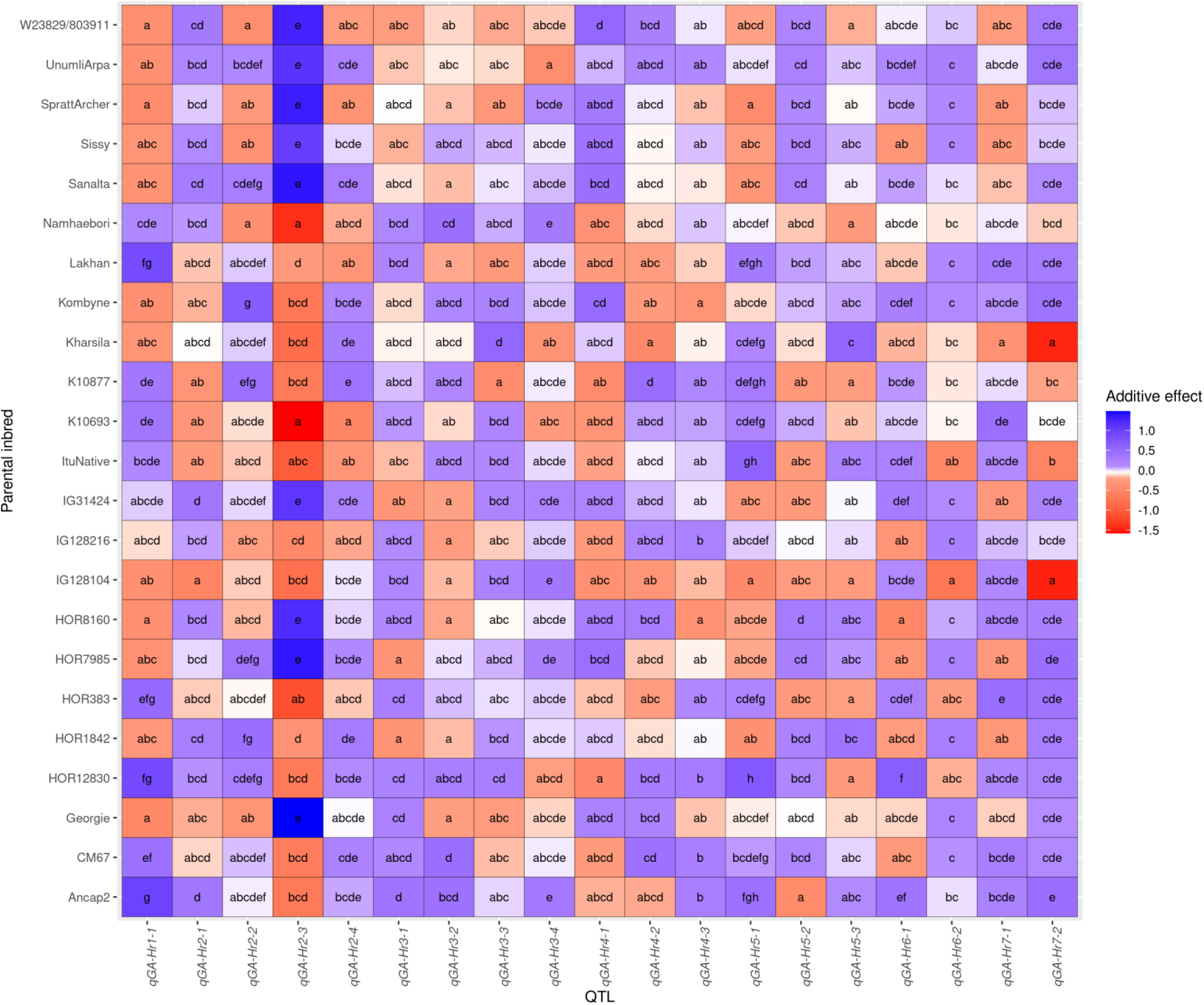
Multiple comparisons of the standardized allele effect for grain area quantitative trait loci (QTLs) detected in a multi-parent population analysis using a parental model. The standardized allele effect for an inbred is the difference between the mean of the estimated allele effect for 23 inbreds and the estimated additive effect of the corresponding inbred. The color code indicates the magnitude of the standardized allele effect. Indexed letters indicate the significant difference (p ≤ 0.05) between the genotypes not sharing the same letter by Tukey’s HSD test.

**Fig. S11:**
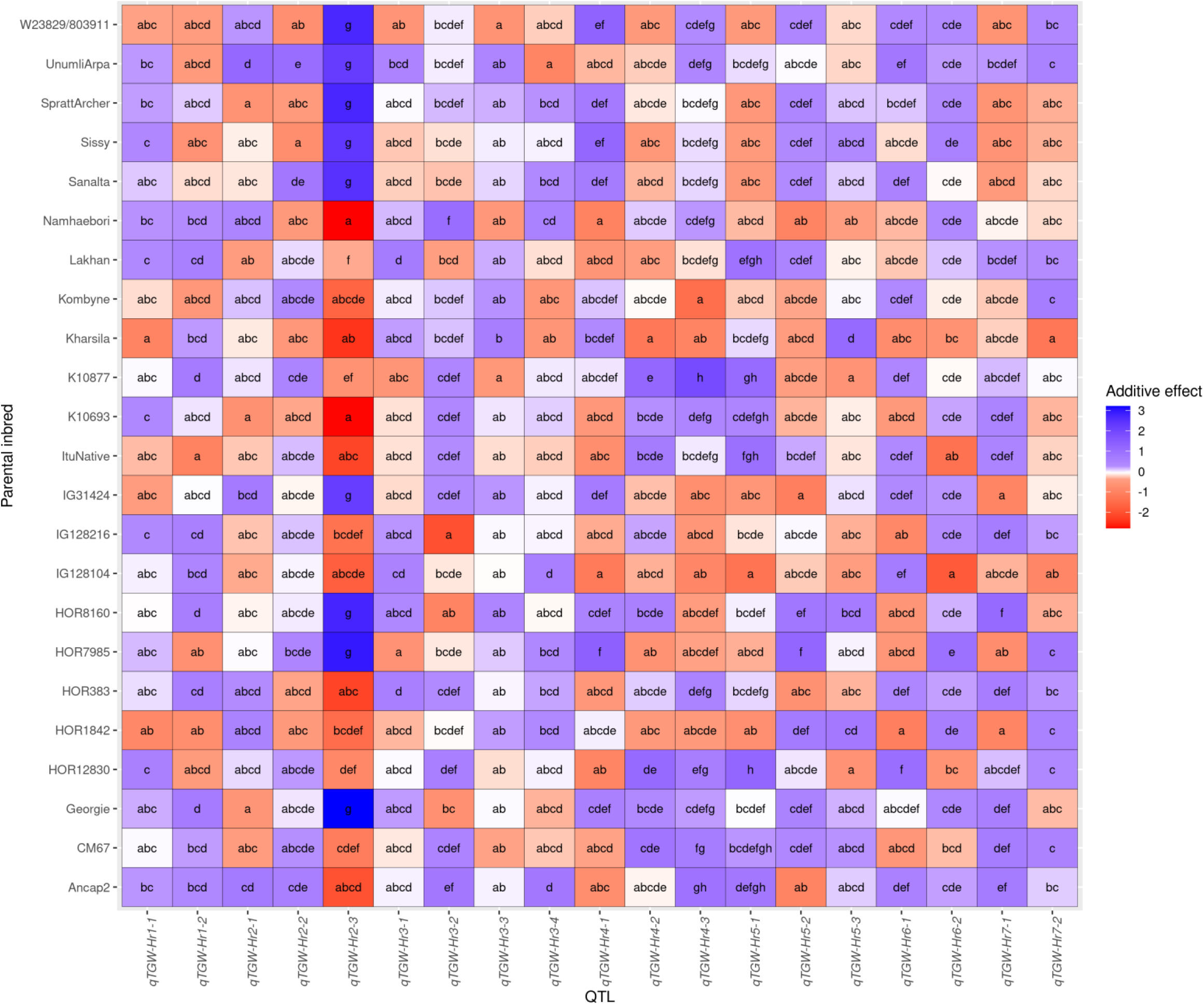
Multiple comparisons of the standardized allele effect for thousand grain weight quantitative trait loci (QTLs) detected in a multi-parent population analysis using a parental model. The standardized allele effect for an inbred is the difference between the mean of the estimated allele effect for 23 inbreds and the estimated additive effect of the corresponding inbred. The color code indicates the magnitude of the standardized allele effect. Indexed letters indicate the significant difference (p ≤ 0.05) between the genotypes not sharing the same letter by Tukey’s HSD test.

**Fig. S12:**
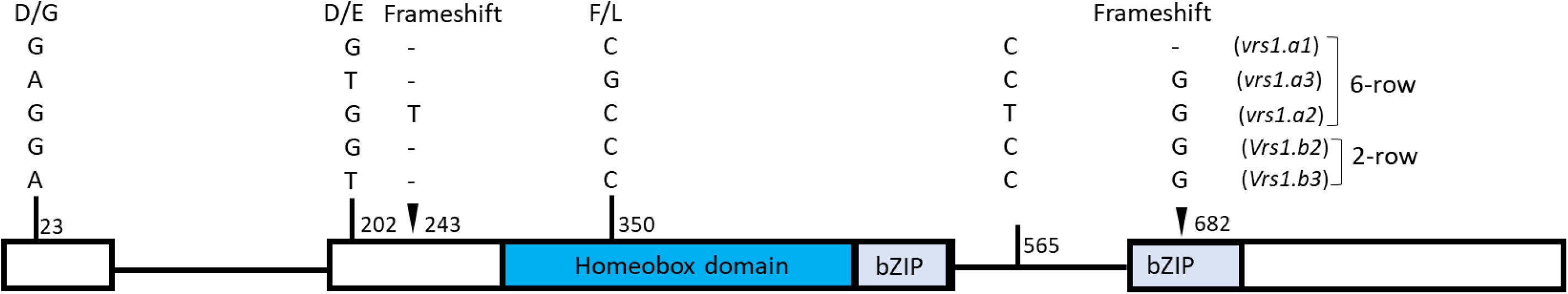
The allelic variant for *Vrs1*. A major QTL at *Vrs1* locus on chromosome 2 associated with grain size and weight (*qHvDRR-GS-14*) was detected in HvDRR populations developed from the genetic cross between 2 and 6-row inbreds. The position adjacent to the polymorphic sites is the relative position from the start of the gene.

**Fig. S13:**
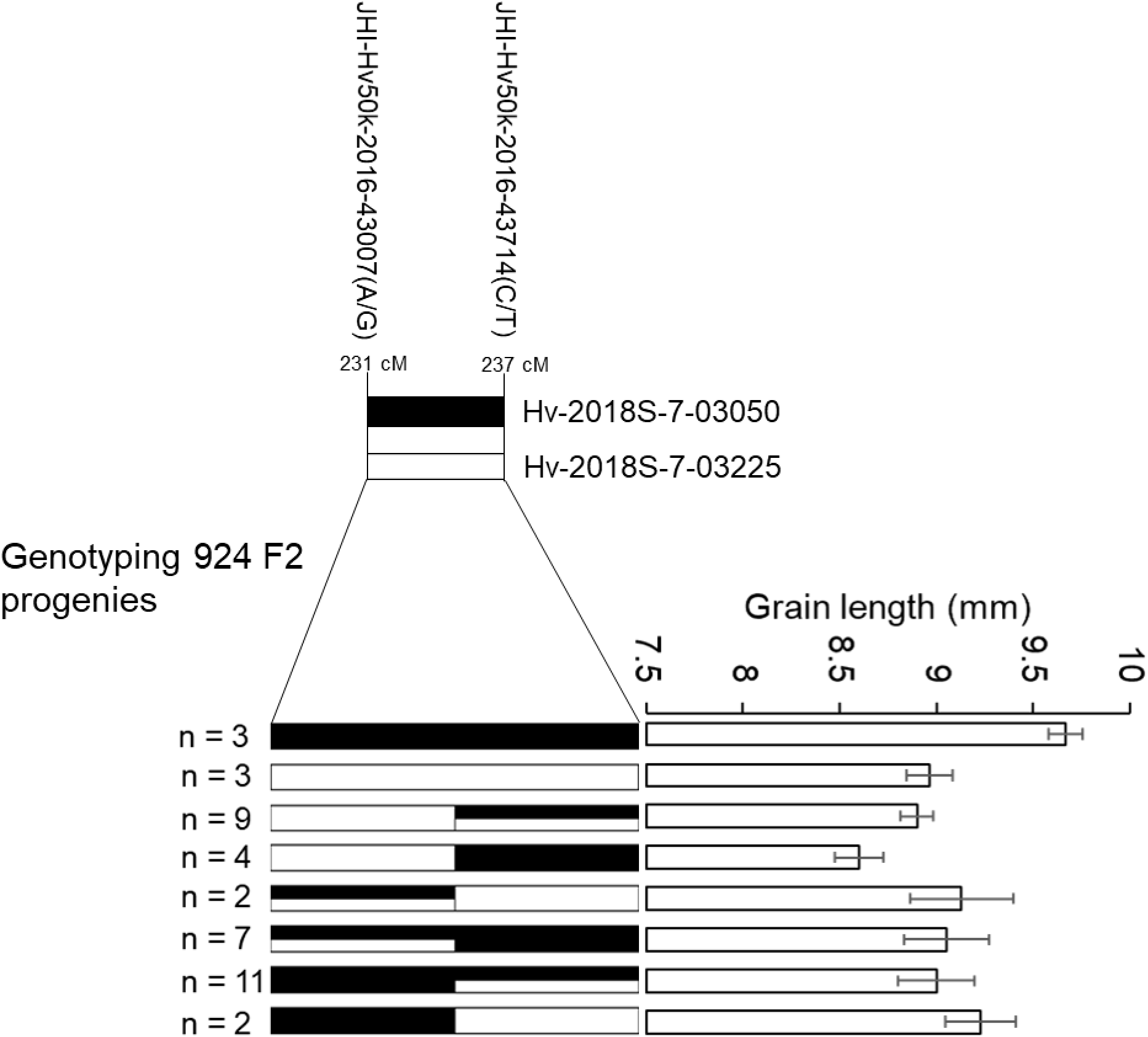
QTL allele effect validation at *qHvDRR-GS-6* on chromosome 1. Two recombinant inbred lines of the HvDRR33 sub-population bearing separate QTL alleles at *qHvDRR-GS-6* but being monomorphic for the other sub-population specific QTLs were selected to produce a high-resolution segregating population. Recombinant F2 progenies segregating for *qHvDRR-GS-6* were selected by genotyping at the left and right border of the QTL. The grain length of the recombinant F2 plants and selected non-recombinants from the segregating population was evaluated. The black and white bar represents the allele from Lakhan and Georgie, respectively.

**Fig. S14:**
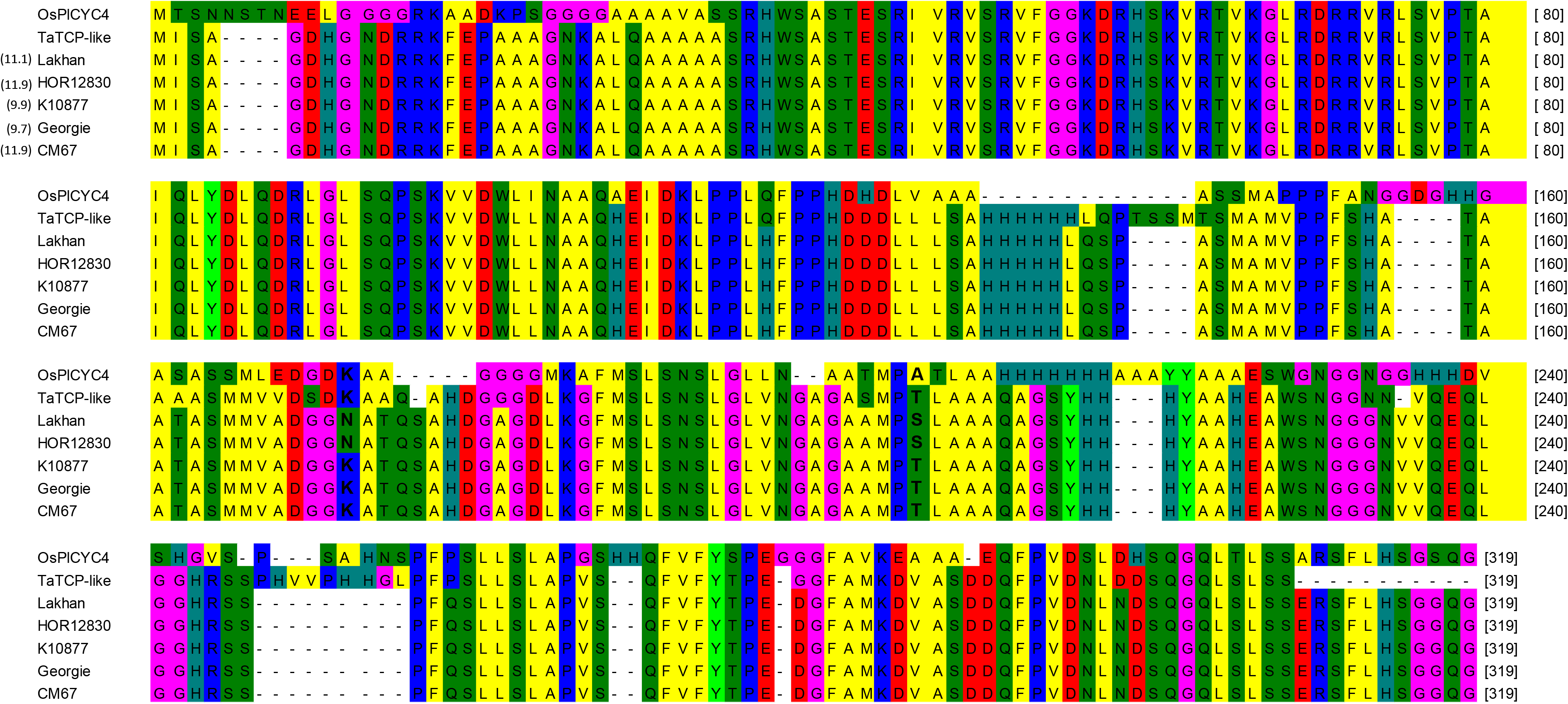
Protein alignment of a candidate gene underlying *qHvDRR-GS-6* on chromosome 1. SNPs caused amino acid substitutions in the conserved domain of HORVU.MOREX.r2.1HG0063230. The protein sequence of the barley gene was highly conserved in rice and wheat. Lakhan and HOR12830 allele at the QTL locus contributed to longer grains. Sites in bold letters indicate amino acid substitutions between parental groups with contrasting additive effects at the *qHvDRR-GS-6*.

